# The shape of evolution: persistent homology of genetic-distance data as an observable of reticulate processes in pathogen and plant genomes

**DOI:** 10.64898/2026.06.18.733071

**Authors:** Alejandro Feged-Rivadeneira, Felipe González-Casabianca, Pablo Cárdenas Ramírez, Andrés Ángel, Vladimir Corredor

## Abstract

Evolutionary biology has long theorized processes—recombination, lineage divergence, drug-resistance sweeps, introgression, refugial persistence—whose signatures in genomic data are incompatible with tree structure. We argue that the *shape* of genetic-distance data, formalized through simplicial complexes and quantified through persistent homology, is a direct observable of these processes. The Vietoris—Rips filtration of a genetic-distance matrix yields the Betti numbers *β*_0_ (connected components), *β*_1_ (loops), and *β*_2_ (cavities); we read *β*_1_ not as a literal count of recombination events but as a quantity that is monotone in effective recombination above a sampling-dependent geometric baseline, and we organise the resulting shapes into a four-letter alphabet of topological primitives (*K*_1_ clonal, *K*_2_ divergence, *K*_3_ reticulation, *K*_4_ higher-order reticulation). Coalescent and Wright—Fisher simulations establish the two load-bearing claims: *β*_1_ rises monotonically with the recombination rate over six orders of magnitude, and persistent-homology features separate reticulate from non-reticulate histories with 98—100% recall (the residual confusion falls entirely within the non-reticulate *K*_1_/*K*_2_ pair, which *β*_1_ does not distinguish). We then apply the pipeline to four empirical systems. (i) On the MalariaGEN Pf7 *Plasmodium falciparum* dataset (*n* = 20,864, 33 countries), per-population *β*_1_ spans two orders of magnitude and diverges significantly from a label—shuffle null (median 20.5, range 8-32); the ordering runs *opposite* to recombination rate-freely-recombining African populations sit lowest and clonal/swept Southeast Asian and Papuan populations highest—because at the population scale *β*_1_ is dominated by demographic structure rather than recombination rate, a point we reconcile explicitly with the controlled dose-response. (ii) Colombian Cauca SP-resistant samples carry *β*_1_ = 12 against a near-clonal SP-sensitive baseline of *β*_1_ = 5 (and two orders of magnitude more total persistence), the high-*β*_1_, multi-origin band of *K*_3_ consistent with resistance carried on several genomic backgrounds. (iii) The Cambodia artemisinin sweep (2008-2018) traces a *K*_3_ → *K*_1_ trajectory, *β*_1_ rising to a mid-sweep peak of 45 and collapsing to 13 at fixation—to our knowledge the first direct observation of a selective-sweep transient in topological coordinates, with the caveat that the per-bin values are medians of three subsamples with wide bars. (iv) On *Arabidopsis thaliana* 1001 Genomes data, Iberian relict populations (Spain, *β*_1_*/n* = 0.64) exceed post-glacial-expansion populations (Sweden 0.54; United Kingdom 0), generalising the framework beyond pathogens. A *P. falciparum* mitochondrial negative control recovers *β*_1_ = 0 across all subsamples, establishing pipeline specificity. Moving above the 1-skeleton, *β*_2_ is zero at the clonal/expansion limits and positive across the reticulate systems; a controlled two-vs-three-way admixture simulation confirms that *β*_2_ separates regimes that share a *β*_1_ profile, while the further suggestion that the ratio *η* = *β*_2_*/β*_1_ separates microevolutionary from macroevolutionary timescales is presented, given the small number of systems and the absence of a *β*_2_ null, as a hypothesis for future testing. Together these results demonstrate that the topology of genetic-distance data is an evolutionary observable, with immediate implications for drug-resistance surveillance in *P. falciparum*.

**Author summary:** Biologists usually picture the history of life as a tree, in which lineages split and never rejoin. Many of the most consequential evolutionary events break that picture: malaria parasites recombine in the mosquito gut, drug-resistant strains arise repeatedly on different genetic backgrounds, and plant populations that survived the Ice Age in southern refuges carry tangled ancestry that no tree can represent. We ask a different question of genetic data—not “what tree fits?” but “what *shape* does the data make?” —and answer it with topological data analysis, which measures shape through three counts (the Betti numbers) of clusters, loops, and higher-order cavities. Loops appear when lineages recombine and rejoin. We show, in simulations and in real *Plasmodium falciparum* and *Arabidopsis thaliana* genomes, that the loop count rises with recombination above a baseline set by finite sampling, cleanly separates recombining from clonal histories, and tracks a real artemisinin-resistance sweep in Cambodia as it rises and collapses over a decade. A non-recombining mitochondrial control correctly shows no loops. The shape of genetic data is thus a direct, tree-free readout of evolutionary process, with immediate value for drug-resistance surveillance.

## 1 Introduction: the problem of seeing evolution

The textbook picture of evolution is a tree. A single ancestral lineage divides; each branch divides again; the whole of life is a nested hierarchy of splits. The image is at least as old as Darwin’s notebooks and is extraordinarily useful. It is also, for a large fraction of evolution’s most consequential moments, wrong in a specific way. The sexual phase of a malaria parasite’s life cycle takes place in the gut of a mosquito, where two parental genomes recombine to produce mosaic offspring [10]. Two influenza strains can meet inside a pig and trade whole genomic segments, occasionally producing a pandemic [50]. Bacteria pass resistance genes sideways across species boundaries at rates that make the concept of “species” itself a topological question [13, 41]. A plant population driven into a Mediterranean refuge during the last Ice Age can survive, preserve its distinctive variation, and later meet the populations that recolonized Northern Europe from elsewhere [58]. In each of these cases, the evolutionary history is not a tree—it is something with loops, bridges, and three-way entanglements: a graph at best, a higher-dimensional object more honestly. This is not a philosophical nicety. It is a measurement problem, because the questions that matter in public health and conservation biology are questions about exactly these non-tree-like events, and the tree representation throws away precisely the information that answers them.

Evolution is among the most thoroughly theorized processes in the natural sciences, yet one of the hardest to observe directly. Since Fisher [21], Wright [53], and Haldane [23] laid the mathematical foundations of population genetics, the discipline has developed an extraordinary theoretical apparatus—adaptive landscapes, fitness valleys, drift barriers, coalescent processes, quasi-species clouds—for reasoning about how populations change over time. But the connection between this apparatus and the data produced by modern genomics remains strikingly indirect. When we sequence a pathogen population, we obtain a collection of genomes; the standard analytical move is to infer a phylogenetic tree [20, 14] and then interpret the tree in terms of the underlying process.

The difficulty, recognized since Doolittle’s challenge to the universal tree of life [13], is that many of the most consequential evolutionary processes are not tree-like. Recombination in *Plasmodium falciparum*-the most lethal human malaria parasite—occurs obligately during the sexual phase in the mosquito vector, generating mosaic genomes from co-infecting strains [10]. The population-genetic consequences of this recombination—the shuffling of resistance alleles, the maintenance of antigenic diversity, the generation of novel genotype combinations—are among the most important determinants of malaria epidemiology and drug resistance [1], [52]. Reassortment among co-infecting influenza viruses shuffles entire genomic segments, occasionally producing pandemic strains [50]. Horizontal gene transfer in bacteria disseminates resistance cassettes across species boundaries at rates that make species delimitation itself a topological question [13], [41]. In all these cases, the evolutionary history is a directed acyclic graph with reticulation, not a tree, and forcing it into a tree discards precisely the information that determines the pathogen’s capacity to adapt.

This is not merely a representational inconvenience. It is a barrier to answering the fundamental questions. Corredor et al. [11] showed that sulfadoxine-pyrimethamine (SP) resistance mutations in *P. falciparum* across Colombia have a single origin in the *dhfr* and *dhps* loci, but disseminated across the Andes—a geographic barrier to gene flow—via migration. Carrasquilla et al. [6] used whole-genome identity-by-descent (IBD) analysis to resolve drug selection and migration in this same inbred Pacific coast population, demonstrating that even small, isolated *Plasmodium* populations remain evolutionarily labile and can successfully adapt to multiple drug regimes through hard and soft selective sweeps enabled by sufficient recombination. Cárdenas, Corredor, and Santos-Vega [3] developed Opqua, a simulation framework explicitly linking epidemiology to sequence evolution, and showed that low transmission, host mobility, and complex life cycles facilitate evolution across fitness valleys—Sewall Wright’s shifting balance theory [53] made concrete in a pathogen context.

But how would one detect these processes in empirical genomic data? A tree shows branches; it does not reveal that a branch passed through a fitness valley. A tree shows tips; it does not reveal that two tips are connected by a recombination event that created a genotypic cycle. The *shape* of the genetic distance data—the topology of the simplicial complex built from pairwise distances—is what encodes these processes.

Topological data analysis (TDA) provides the mathematical machinery. Chan, Carlsson, and Rabadán [8] demonstrated that persistent homology distinguishes clonal from reticulate evolution: the first Betti number *β*_1_, counting independent loops in the simplicial complex, is zero for tree-like histories and positive for histories involving recombination. Lesnick and Rabadán [35] formalized this through the novelty profile, proving stability results. Humphreys et al. [28] showed fast recombination rate estimation. Lum et al. [38] established a shape vocabulary for Mapper output—flares, loops, clusters—applied across cancer genomics [40], neuroscience [47], and athletics. In our prior work, we developed a Mapper pipeline for genomic epidemiology [34], [18], [19], introducing the Point Intersection Network (PIN) to identify the specific genotypes bridging distinct subpopulations.

What has been missing is a synthesis connecting these topological tools to the fundamental questions of evolutionary biology. The purpose of this paper is not to present a method for its own sake but to argue that simplicial complexes—the geometric objects at the heart of TDA—are natural representations of the structure of genetic variation, and that their topological invariants provide quantitative answers to questions that Lewontin, Maynard Smith, and Wright posed in qualitative terms. We use *P. falciparum* micro-epidemiology on the Colombian Pacific coast as an emblematic system, alongside coalescent and Wright-Fisher simulations that anchor the interpretation, and then test the framework across four empirical genomic systems (malaria, *Arabidopsis*, and a mitochondrial negative control). We test its consistency across ten synthetic and genomic constructions in Supplementary Section S4. Our position is humble in tone but, we hope, justified by the data: across the malaria and *Arabidopsis* systems we examined, the alphabet behaves as a coordinate of evolutionary process rather than a coordinate of malaria genetics specifically. Whether it travels further—to other taxa, or to non-genomic transmission systems—is a question we leave to future work rather than claim here.

## 2 Simplicial complexes and their relevance to evolutionary biology

### 2.1 Intuition: what simplicial complexes are, and why evolutionary biology needs them

Before the formal definitions, it is worth pausing on the intuition. A simplicial complex is the geometric generalization of the graph that practitioners of phylogenetics already use every day. A graph has dots—call them *vertices*, representing genotypes here—and lines—call them *edges*, representing “these two genotypes are similar enough to be connected.” A simplicial complex keeps the dots and lines but also adds *filled triangles* whenever three genotypes are mutually similar, *filled tetrahedra* whenever four are, and so on. The complex is thus an honest geometric object—one can imagine it as a lumpy surface glued together from triangles and higher-dimensional pieces—rather than a branching diagram imposed by an algorithm.

Why does this matter for evolutionary biology? A phylogenetic tree is, mathematically, a simplicial complex with two restrictions: first, it is only one-dimensional (no filled triangles), and second, it has no closed loops. Both restrictions are biological assumptions, not properties of the data. The first assumes that three-way genetic coherence is never an object of interest in its own right. The second assumes that there has been no recombination, no hybridization, no horizontal transfer, and no admixture—a fiction for any realistic pathogen or plant population. When we drop those two assumptions and simply let the data tell us which genotypes are mutually similar at each distance threshold, the result is a simplicial complex, and the “shape” of that complex carries information that no tree can encode.

Three pictures anchor what this shape tells us. A collection of disconnected clumps records *how many distinct populations* are resolvable at the resolution being examined. A closed loop—three or more clumps hooked together in a ring, with no filled interior—records *a reticulation event*: ancestors met and exchanged DNA. A hollow shell—four or more clumps configured like the surface of a balloon, with a cavity inside—records *multi-way incompatibility*: three or more lineages related simultaneously in a way that no single common ancestor at this scale can explain. Counting these features is what the Betti numbers *β*_0_, *β*_1_, *β*_2_ do. They are the topological analogue of reading the number of islands, bridges, and arenas in a landscape from a satellite photo.

Topological data analysis (TDA) is the machinery that extracts these counts from raw genetic-distance data in a principled, scale-aware way. Rather than commit to a single “correct” distance threshold *ε*, TDA watches how the simplicial complex grows as *ε* increases from zero (every genotype is isolated) to infinity (every genotype is connected to every other). Features that appear briefly and vanish are noise; features that persist over a long range of *ε* are robust evolutionary signal. This is what we mean by *persistent homology*. The complementary *Mapper* algorithm takes the same data and produces a compressed, interpretable 1-skeleton—a graph that is easier to look at than the raw distance matrix but, unlike a tree, honestly displays loops and branchings where they exist.

A practical consequence is that the method never forces the data into a structure it does not have. If the underlying process is strictly tree-like (clonal descent, strict allopatry), the simplicial complex collapses to a tree, and *β*_1_ = *β*_2_ = 0. If there is pairwise reticulation (recombination, admixture between two populations), loops appear and *β*_1_ *>* 0. If there is multi-way reticulation (three or more simultaneously admixing lineages, or higher-order incompatibility), cavities appear and *β*_2_ *>* 0. Each Betti number is thus a *direct evolutionary readout* that becomes non-zero only when the process that would generate it is actually present in the data. The rest of this section makes these statements precise. Readers who prefer to skim the formalism can proceed to Section 2.5 or Section 7.4 without losing the biological thread; a fully analytical scaffold connecting coalescent rates to expected Betti numbers, and a dictionary of the topological objects one encounters in genomic data (loops, wedges, tori, shells), is developed in the Supplementary Information.

### 2.2 Mathematical foundations

A simplicial complex is a combinatorial object that generalizes a graph to higher dimensions. Where a graph consists of vertices (0-simplices) and edges (1-simplices), a simplicial complex additionally contains triangles (2-simplices), tetrahedra (3-simplices), and their higher-dimensional analogues. The formal definition is as follows.

#### Definition 1

(Abstract simplicial complex). *An* abstract simplicial complex *on a vertex set V is a collection* Σ *of finite subsets of V (called* simplices*) such that: (i) every singleton {v}* ∈ Σ *for v* ∈ *V* ; *and (ii) if σ* ∈ Σ *and τ* ⊆ *σ, then τ* ∈ Σ. *A simplex σ with* |*σ*| = *k* + 1 *is called a k-simplex. The* dimension *of* Σ *is the maximum k such that a k-simplex exists in* Σ.

#### Example 1.

*A graph is a simplicial complex of dimension at most 1: its 0-simplices are vertices, its 1-simplices are edges. A triangle (three mutually connected vertices with the interior “filled in’) is a 2-simplex. A hollow triangle (three edges without the filled interior) is a simplicial complex of dimension 1 that is* not *contractible-it contains a loop that cannot be continuously shrunk to a point*.

The topological properties of a simplicial complex are captured by its *homology groups*, which count “holes” of each dimension.

#### Definition 2

(Simplicial homology). *Let* Σ *be a simplicial complex and* F *a field (typically* F = ℤ/2ℤ*). The k-th* chain group *C*_*k*_(Σ; F) *is the F-vector space with basis the k-simplices of* Σ. *The* boundary operator ∂_*k*_ : *C*_*k*_ → *C*_*k*−1_ *maps each k-simplex to the formal sum of its* (*k* − 1)*-dimensional faces. The k-th homology group is:*

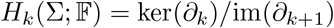

*The k-th* Betti number *β*_*k*_ = dim(*H*_*k*_) *counts the number of independent k-dimensional “holes’:*

- *β*_0_: *number of connected components*.
- *β*_1_: *number of independent loops (1-cycles not bounding a filled surface)*.
- *β*_2_: *number of independent cavities (2-cycles not bounding a filled volume)*.

Concretely, the boundary operator acts on a *k*-simplex *σ* = [*v*_0_, *v*_1_, …, *v*_*k*_] as 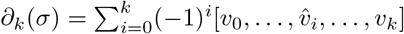, where 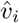 denotes omission. For a 1-simplex (edge) [*v*_0_, − *v*_1_], ∂_1_([*v*_0_, *v*_1_]) = *v*_1_ − *v*_0_: the boundary of an edge is the difference of its endpoints. For a 2-simplex (triangle) [*v*_0_, *v*_1_, *v*_2_], ∂_2_ = [*v*_1_, *v*_2_] − [*v*_0_, *v*_2_] + [*v*_0_, *v*_1_]: the boundary of a triangle is the sum of its three edges. A cycle is an element of ker(∂_*k*_): a chain with no boundary. A loop in the Mapper 1-skeleton, for instance, is a 1-cycle—a sequence of edges that closes on itself. The first Betti number *β*_1_ counts how many such loops are independent, modulo those that bound filled triangles. In the evolutionary context, an independent 1-cycle that survives at long persistence is a reticulate connection that cannot be “explained away” by adding a filled simplex-an irreducible departure from tree-like descent. One caveat must be stated at the outset and is carried through the rest of the paper: the Vietoris-Rips complex of a *finite, noisy* metric space generates 1-cycles from sampling geometry alone, independent of any reticulation, as the random-geometric-complex literature establishes [62],[63] The integer *β*_1_ of a real sample is therefore a sum of reticulation signal and a sampling-dependent geometric baseline. The defensible reading—which our simulations (Section 7.1) and negative controls (Section 7.8) support—is that *β*_1_ is *monotone in effective recombination above that baseline*, not that it is a literal count of recombination events. We make the baseline explicit wherever we report *β*_1_ on empirical data.

The key property of homology is *topological invariance*: continuous deformations of the complex (adding a vertex inside an existing simplex, stretching or compressing) do not change the Betti numbers. This makes homology robust to the kind of noise that pervades biological data: small changes in sequencing quality, sampling density, or distance estimation do not alter the topological features.

### 2.3 From distance data to simplicial complexes

Given a set of genomic sequences and a distance function (p-distances, Hamming distances, identity-by-descent), the standard construction for extracting topology is the Vietoris-Rips filtration.

#### Definition 3

(Vietoris—Rips complex). *Let* (*X, d*) *be a finite metric space (e*.*g*., *n genomes with pairwise distances). For each ε* ≥ 0, *the* Vietoris—Rips complex VR_*ε*_(*X, d*) *is the abstract simplicial complex whose k-simplices are all subsets {x*_0_, …, *x*_*k*_*}* ⊆ *X with d*(*x*_*i*_, *x*_*j*_) ≤ *ε for all i, j. As ε increases from 0 to* ∞, VR_*ε*_ *grows from a collection of isolated vertices to a single large simplex containing all points, forming a nested sequence called a* filtration:

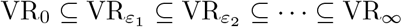

#### Definition 4

(Persistent homology). *Tracking the homology groups H*_*k*_(VR_*ε*_) *across the filtration produces a sequence of vector spaces connected by linear maps (a* persistence module*). A topological feature is “born’ at ε* = *b when a new homology class appears and “dies’ at ε* = *d when it merges with an older class. The collection of intervals {*[*b*_*i*_, *d*_*i*_)} *is the* persistence barcode. *The* persistence *π*_*i*_ = *d*_*i*_ − *b*_*i*_ *measures the robustness of the feature: long bars represent genuine structure, short bars are noise*.

The *stability theorem* [9] provides a partial guarantee: if two distance matrices *D, D*^*′*^ satisfy ∥*D* − *D*^′^∥_∞_ ≤ *δ*, then the bottleneck distance between their persistence *diagrams* is at most *δ*. We are careful about what this does and does not control. It bounds the diagram in the matching metric—it does not bound the integer count *β*_1_ of bars above a fixed persistence threshold, which is the quantity we ultimately tabulate. A single *δ*-perturbation can move a bar across any fixed cutoff and change the count by one; the “2,857 bars with persistence exceeding 0.01” we report below is exactly the kind of thresholded integer for which bottleneck stability offers no protection. We therefore treat *β*_1_ counts as scale-robust empirically (Section 7.9, where we sweep the filtration and show the cumulative count plateaus) rather than as stable by theorem. The stability theorem also does *not* transfer to Mapper, whose output is known to be sensitive to its cover and clustering parameters [7]; we treat the Mapper graph and its first Betti number as a separate, less stable object throughout (Section 4). Finally, the contrast with phylogenetic trees should not be overstated: while small changes in a distance matrix can reorder tree topologies, the space of phylogenetic trees carries a well-defined geometry with its own continuity results [64]. The honest claim is that persistence diagrams come with an explicit Lipschitz guarantee in a natural metric, which is convenient for our purposes, not that trees lack any stability theory.

### 2.4 Why simplicial complexes are the right representation for evolutionary data

The claim that simplicial complexes are natural representations of genetic variation requires justification. Three properties make them uniquely suited:

#### First, they are coordinate-free

Unlike principal component analysis, which requires choosing axes, or multidimensional scaling, which embeds the data in Euclidean space, simplicial homology depends only on the distance matrix. Two datasets with different numbers of SNPs, different genotyping platforms, or different reference genomes, but the same pattern of pairwise distances, will produce the same Betti numbers. This is essential for comparing topological structure across studies, species, and data types.

#### Second, they capture non-tree-like structure

A phylogenetic tree is a simplicial complex with *β*_1_ = 0: it has no loops. The statement *β*_1_ *>* 0 is therefore a direct, quantitative measure of the degree to which the data depart from tree-like evolution. This is precisely what Chan et al. [8] exploited: recombination creates loops that trees cannot represent, and *β*_1_ measures their accumulation above the finite-sample baseline.

#### Third, they are multiscale

The persistence barcode provides information about the topological structure at every distance threshold simultaneously. This is critical for genomic data, where the “right” distance scale is rarely known in advance. A pair of genotypes that are closely related at the SNP level may be distantly related at the IBD level; the persistence barcode reveals the topology at both scales, without requiring the analyst to choose.

These properties address, in mathematical terms, a problem that Lewontin [37] identified in qualitative terms: the need for a representation of genetic variation that is richer than summary statistics (*F*_ST_, heterozygosity, *D*) but more structured than the raw sequence data. The Betti numbers of the Vietoris—Rips filtration are such a representation. *β*_0_(*ε*) generalizes *F*_ST_: it counts how many genetically distinct groups exist at threshold *ε*, without forcing a fixed number of clusters. *β*_1_(*ε*) has no analogue in classical population genetics: it measures the degree of reticulation, the non-tree-likeness, the topological complexity that recombination, gene flow, and horizontal transfer create.

### 2.5 Higher-dimensional simplicial structure: what *β*_*k*_ encodes for *k* ≥ 2

The empirical analyses in the malaria literature [8], [34] and in our own case studies up to this point have used only the 1-skeleton of the Vietoris-Rips filtration: vertices, edges, and *β*_0_, *β*_1_. This is a choice of convenience, not of principle. The Vietoris-Rips complex VR_*ε*_(*X, d*) is a *flag complex* : whenever *k* + 1 points are pairwise within distance *ε*, the corresponding *k*-simplex is included. A 2-simplex {*x*_*i*_, *x*_*j*_, *x*_*k*_*}* is present when all three pairwise IBS/IBD distances are ≤ *ε*. In evolutionary terms, such a 2-simplex is a certificate of three-way near-identity: three genotypes that plausibly share recent common ancestry or admixture source at the resolution set by *ε*.

Higher Betti numbers capture progressively more multi-way structure. *β*_2_ counts independent 2-cycles in the 2-skeleton—hollow “shells” formed when pairwise and some triple-wise proximity hold but the whole configuration is not tri-filled. In a recombining population, a *β*_2_ feature is a configuration of (at least) four lineage clusters such that every pair is genetically close enough to be edge-connected, some triples are close enough to be triangle-filled, but the tetrahedral closure fails: there remain incompatible triples that cannot be filled by a single common ancestor at that scale. This is a signature of multi-way incompatibility between lineages that cannot be reduced to a set of pairwise reticulation events: a higher-order analogue of the quartet incompatibility that phylogenetic network methods [29] attempt to detect, but here expressed as a coordinate-free topological invariant rather than as a list of conflicting quartets. The biological expectation is that *β*_2_ grows with the number of co-circulating, partially-admixing lineages that occupy genetic-distance space in geometrically non-convex configurations—precisely the settings that the *K*_4_ primitive and the *K*_2_ refugium-relict compound describe.

Three simplicial-complex quantities will recur in the empirical sections below. The *f -vector f* (VR_*ε*_) = (*f*_0_, *f*_1_, *f*_2_, …) records the number of *k*-simplices at scale *ε* and tracks the raw combinatorial complexity of the cover. The *Euler characteristic χ*(VR_*ε*_) = ∑_*k*_(−1)^*k*^*f*_*k*_ =∑_*k*_(− 1)^*k*^*β*_*k*_ summarizes the complex by a single integer that nonetheless detects higher-dimensional voids: *χ* < 0 when 1-cycles dominate, *χ* > 0 when 2-cavities dominate. The *higher-order index η* = *β*_2_*/β*_1_ isolates the fraction of topology not explainable by pairwise cycles; it is a zero for trees and pure pairwise reticulation and becomes positive when genuinely multi-way structure appears. Within the macroevolutionary companion paper [26] this language overlaps with the higher-order network literature [43], [25], [24]: there, edges of a weighted hypergraph encode simultaneous *k*-way interactions, and their nerve is precisely the simplicial complex we build here. The evolutionary reading of this correspondence is that the Vietoris-Rips 2-skeleton is a *three-way* genetic-similarity network, distinct in content from—and in general not reducible to—the pairwise distance graph that has carried most of the attention in phylogenetic network theory.

## 3 Theoretical connections to evolutionary biology

### 3.1 Maynard Smith and the topology of recombination

Maynard Smith’s *The Evolution of Sex* [39] posed the question: why does recombination persist despite its twofold cost? The theoretical resolution involves recombination’s advantage in generating novel genotype combinations [39], [42]. The topological framework provides an aggregate measure of this consequence. A recombination event between divergent lineages can create a cycle in genetic-distance space—two lineages diverge and reconnect through a hybrid genotype—producing a *β*_1_ bar whose persistence reflects the parental divergence [35]. The mapping is not one-to-one in either direction: some events fail to leave a Rips-visible cycle (their parents are too close, or the cycle is filled by triangles), and finite sampling produces cycles in the absence of any event. What survives these caveats is a monotone relationship in aggregate—more effective recombination yields larger *β*_1_ and total persistence Π_1_ =∑_*i*_ *π*_*i*_ above the sampling baseline—which we verify by direct dose-response (Figure 5). Π_1_ is thus best read as a relative index of cumulative reticulate innovation, and the rise of *β*_1_ above its clonal baseline as the observable onset of effective recombination.

### 3.2 Wright’s landscapes and fitness valley topology

Wright’s adaptive landscape [53]—populations exploring a multi-peaked fitness surface—was formalized computationally by Kauffman’s *NK* model [31]. Cárdenas, Corredor, and Santos-Vega j brought this to pathogens, showing that population structure determines whether valley crossings occur. A valley crossing leaves a predictable topological signature: a thin bridge between two dense subcomplexes (the pre- and post-crossing populations), with *β*_0_ temporarily increasing during the bottleneck then collapsing as the new population expands. The Mapper 1-skeleton shows this as a chokepoint; the PIN shows a single high-centrality genotype at the bridge. This is Wright’s shifting balance made geometrically observable.

### 3.3 Lewontin’s partitioning, generalized

Lewontin’s apportionment of human genetic diversity [36] partitioned variation into within-population, between-population-within-race, and between-race components, finding that most variation is within populations. The topological framework generalizes this partitioning: *β*_0_(*ε*) is a continuous function describing how many genetically distinct groups exist at every distance threshold, and *β*_1_(*ε*) measures how much those groups are connected by reticulate exchange. Where Lewontin’s partitioning required choosing a level of the hierarchy (populations, races), the persistence barcode provides the full multiscale picture.

### 3.4 Corredor’s resistance dissemination as a topological question

Corredor et al. [11] showed that SP resistance has a single origin but disseminated across the Andes. Carrasquilla et al. [6] showed that even small, inbred populations maintain the capacity for selective sweeps. The question-*through what topology does resistance spread?* —has different answers with different intervention implications. A selective sweep (*β*_0_ collapsing) is a fait accompli. Ongoing gene flow (*β*_1_ increasing) suggests that reducing connectivity could slow spread. A single bridge event (*β*_1_ appearing at one scale) is an opportunity for targeted intervention. TDA can distinguish these scenarios.

## 4 The dual TDA framework

### 4.1 The Mapper algorithm

We implement the Mapper algorithm [49] in the R package `mapperKD` (github.com/minigonche/mapperKD).

#### Definition 5

(Mapper construction). *Given a finite metric space* (*X, d*) *with* |*X*| = *n, a filter function f* : *X* → ℝ, *resolution r* ∈ ℕ, *overlap g* ∈ (0, 100), *and clustering algorithm δ:*

1. *Compute the cover U* = {*U*_1_, …, *U*_*r*_} *of* im(*f*) *by overlapping intervals of width w* = (*f*_max_ − *f*_min_)*/*(*r* − (*r* − 1)*g/*100) *with step s* = *w*(1 − *g/*100).
2. *For each U*_*i*_, *compute the preimage f* ^−1^(*U*_*i*_) = {*x* ∈ *X* : *f* (*x*) ∈ *U*_*i*_} *and apply δ to the restricted distance matrix* 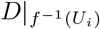 *to obtain clusters* 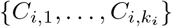.
3. *The* Mapper complex ℳ *is the nerve of the cover {C*_*i,j*_*}: its vertices are the clusters, and k* +1 *clusters span a k-simplex if and only if their intersection is non-empty*.

*The* 1-skeleton *of* ℳ *is the graph of vertices and edges*.

The Mapper complex is related to the Reeb graph of *f* by the Nerve Theorem [27]: if the cover is sufficiently fine and the clusters are convex, the nerve has the same homotopy type as the underlying space. In practice, the Mapper 1-skeleton provides a simplified, interpretable representation of the global topology.

#### Definition 6

(Point Intersection Network (PIN)). *Given a Mapper 1-skeleton with vertices V and point sets P* (*v*_*j*_) ⊆ *X, define the* intersection centrality *of x*_*i*_ *as κ*(*x*_*i*_) = |{*v*_*j*_ ∈ *V* : *x*_*i*_ ∈ *P* (*v*_*j*_)}|. *For each v*_*j*_, *let* 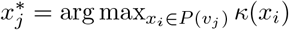. *The PIN is the directed graph where each* 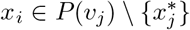 *has an edge to* 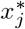.

Nodes with high in-degree in the PIN are the genotypes that bridge distinct temporal clusters—the recombinants, migrants, or transmission links that connect otherwise separate subpopulations. Vhen geo-referenced, the PIN reveals the spatial structure of gene flow.

Mapper depends on three hyperparameters (resolution *r*, overlap *g*, clustering scale *h* or DBSCAN *ε*) whose ad-hoc tuning is a recognised reproducibility concern. Supplementary Section S3 develops a reproducible pipeline for Mapper: a wavelet-like heatmap that summarises the persistent homology of the Mapper 1-skeleton across the entire parameter grid, and a Bayesian-optimisation procedure (GP surrogate, Expected Improvement, Mátern-5/2 kernel) that selects the hyperparameter triple maximising a topological fitness functional. The *empirical* Mapper graph we interpret quantitatively—the Guapi/Cauca *P. falciparum* 1-skeleton (Section 7.11) —uses the parameters this procedure selects (*r* = 20, *g* = 30%, *h* = 0.05). The synthetic 1-skeletons in Figure 7 are illustrative and use a fixed exploratory setting (*r* = 15, *g* = 40%) stated in the text; they are not used for any quantitative claim. We also note honestly (Supplementary Section S3, remarks) that the optimisation anchors to a persistent-homology reference whose filtration scale *ε* is itself a discretionary choice, so the procedure reduces but does not wholly eliminate the analyst’s degrees of freedom.

### 4.2 Persistent homology

We compute persistent homology on the Vietoris—Rips filtration using Ripser [45]. From the persistence barcode in dimension *k*, we extract: the number of bars |ℬ_*k*_|, the total persistence Π_*k*_ = ∑ _*i*_(*d*_*i*_ − *b*_*i*_), and the maximum persistence 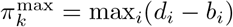.

#### Notation: three distinct first Betti numbers

This paper computes first Betti numbers of three different objects, and we keep them notationally distinct to avoid the conflation that the shared symbol *β*_1_ invites. (i) 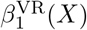 is the first Betti number of the Vietoris-Rips persistent homology of a point cloud *X* (a sample of genotypes), read at a stated filtration scale or as a total bar count; this is the primary observable of the paper. (ii) 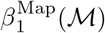 is the first Betti number—equivalently the cycle rank |*E*| − |*V*| + *c*—of a *Mapper nerve ℳ*, a compressed graph built on clusters of *X*, not on the samples themselves. (iii) When we restrict to a subset *X*^*′*^ ⊂ *X* (e.g. the SP-resistant Colombian samples) we write 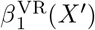 and always state the subset. These are different mathematical objects on different vertex sets and are *not* interchangeable; in particular a Mapper-graph cycle rank and a point-cloud bar count need not agree, and where both are reported we say so explicitly rather than treating their similarity as corroboration.

## 5 The evolutionary shape dictionary: alphabet and vocabulary

A clarifying distinction, suggested in discussions of an earlier draft, is the difference between an *alphabet* and a *vocabulary*. The alphabet is the small, finite set of topologically primitive categories-simplicial-complex types that differ from one another in fundamental Betti-number signatures, not merely in dose. The vocabulary is everything built from the alphabet: quantitative variants along a continuum within one category, temporal trajectories that move between categories, and biogeographic compounds in which alphabet primitives are juxtaposed across geographic scales. This section gives the alphabet first as a four-letter primitive list with a single comparative parameter table, and then catalogues the vocabulary of compounds and trajectories that the empirical sections will use.

### 5.1 The alphabet: four topologically primitive categories

#### Template *K*_1_: Clonal

*β*_0_ = 1 (asymptotically), *β*_1_ = 0, *β*_2_ = 0. 1-skeleton: a path or near-tree. PIN: sparse. Pure vertical descent without reticulation; the topological null. Validated by simulation and by the *P. falciparum* mitochondrial negative control.

#### Template *K*_2_: Divergence

*β*_0_ ≥ 2 (with persistent *β*_0_ gap), *β*_1_ = 0, *β*_2_ = 0. 1-skeleton: tree graph with flares [38]. PIN: centrality at branching node. Two or more lineages without subsequent reticulation; speciation in progress. Distinguished from *K*_1_ not by topological category but by persistence-diagram structure (*K*_2_ produces multiple finite-death *H*_0_ bars).

#### Template *K*_3_: Reticulation

*β*_0_ = 1, *β*_1_ elevated above the clonal baseline, *β*_2_ = 0. 1-skeleton: persistent cycles. PIN: recombinant or admixed genotypes bridging parental clusters. The signature of *K*_3_ is not the bare presence of a single loop—finite samples carry a baseline of cycles from sampling geometry (Section 2.1)—but *β*_1_ (and the total persistence Π_1_) rising monotonically above that baseline, with long-persistence cycles spanning divergent backgrounds [35]. The Wright—Fisher dose-response (Figure 5) establishes the monotonicity directly: *β*_1_ and Π_1_ increase with the mating-system recombination rate over six orders of magnitude. Maynard Smith’s signature, made quantitative as a dose rather than a count.

#### Template *K*_4_: Higher-order reticulation

*β*_2_ ≥ 1, with *β*_1_ ≥ 3 as a structural prerequisite. 1-skeleton: dense reticulation; 2-skeleton contains hollow shells (configurations of four or more clusters whose pairwise edges are filled, some triples are tri-filled, but the whole tetrahedral closure fails). This is the genuinely 3-or-more-way reticulate regime: configurations that no number of pairwise events can account for. The higher-order index *η* = *β*_2_*/β*_1_ summarizes the strength of the higher-order signal relative to the pairwise. The categorical *K*_3_/*K*_4_ distinction (*β*_2_ = 0 vs *β*_2_ *>* 0) is validated by controlled two-vs-three-way admixture simulation and present empirically in the malaria and *Arabidopsis* reticulate systems; the finer claim that *η* itself separates evolutionary timescales is treated as a hypothesis (Section 7.10).

The four primitives are mutually exclusive at the topological level: *K*_1_ has trivial topology; *K*_2_ adds disconnected components but no cycles; *K*_3_ adds 1-dimensional cycles without higher closure; *K*_4_ adds 2-dimensional cavities. They are exhaustive *at the level of Betti numbers*—no other Betti-class arises from coalescent dynamics on the data scales we examine. We are candid about what persistent homology alone can and cannot separate among these four. The sharp, well-supported distinction is binary: reticulate (*K*_3_,*K*_4_; *β*_1_ elevated above baseline) versus non-reticulate (*K*_1_, *K*_2_; *β*_1_ at baseline). Within the non-reticulate class, *K*_1_ and *K*_2_ share the same low *β*_1_ and are separated only by *H*_0_ *persistence* structure and non-topological graph features, not by *β*_1_ (Section 7.1); the simulation classifier confirms this, confusing exactly the *K*_1_/ *K*_2_ pair while never confusing reticulate with non-reticulate. Within the reticulate class, *K*_3_ and *K*_4_ are separated by the 2-skeleton (*β*_2_ *>* 0). The honest summary is therefore that the topology layer carries one categorical axis (reticulate vs not) cleanly, a second axis (*K*_3_ vs *K*_4_) at the cost of computing *β*_2_, and the *K*_1_ vs *K*_2_ distinction not at all without complementary features. Finer regimes that share a Betti signature but differ in geometry—introgression that produces no cycles, selective sweeps that pull samples into a tight cluster, hybrid clines that form a continuous gradient—are invisible to the Betti numbers and require non-topological information (sampling time, geography, or a geometric operator on the distance matrix) to separate; we treat them as vocabulary built on the four primitives (Section 5.2) rather than as additional alphabet letters, and leave their systematic geometric characterization to future work.

Table 1 summarises the four primitives by their Betti-number signatures, evidence status, and a representative empirical example.

**Table 1:** The four-letter topological alphabet, defined by Betti-number signatures. The magnitude of *β*_1_ within *K*_3_ (above the finite-sample baseline) and the higher-order index *η* = *β*_2_*/β*_1_ within *K*_4_ are the two quantitative axes the vocabulary uses to describe variation within and between primitives. “Validated” indicates support from both controlled simulation and at least one empirical system; *K*_2_ is marked “Theoretical” because its distinction from *K*_1_ rests on *H*_0_ persistence structure and non-topological features rather than on *β*_1_.

| Template | $\beta_0$ | $\beta_1$ | $\beta_2$ | $\eta = \beta_2/\beta_1$ | Empirical example | Status |
| --- | --- | --- | --- | --- | --- | --- |
| $\mathcal{K}_1$ Clonal | 1 (stable) | 0 | 0 | – | Pf mito $n = 300$ | Validated |
| $\mathcal{K}_2$ Divergence | $\geq 2$ (gap) | 0 | 0 | – | coalescent simulation | Theoretical |
| $\mathcal{K}_3$ Reticulation | 1 | $\geq 1$ | 0 | 0 | Cauca SP-R ($\beta_1 = 12$) | Validated |
| $\mathcal{K}_4$ Higher-order | 1 | $\geq 3$ | $\geq 1$ | $> 0$ | *Arabidopsis* Spain | Validated |

### 5.2 The vocabulary: compounds and trajectories

The four-letter alphabet does not exhaust evolutionary scenarios; it is the substrate from which the vocabulary is built. We organise the compounds into two families: (i) *quantitative variants* along the dose axis within one alphabet primitive, and (ii) *temporal trajectories* that move a population between primitives over time. A third family—biogeographic compounds—combines primitives across geographic scales and is treated under the macroevolutionary *A* vocabulary.

#### Quantitative variants within *K*_3_

The recombination template’s *β*_1_ is a continuum, and biologically distinct regimes occupy different bands:

- *Sparse K*_3_: *β*_1_ ∈ [1, 5]; one or a few recombination events between two parental backgrounds. Maynard Smith’s textbook signature.
- *Multi-origin K*_3_: *β*_1_ ≫ 1; multiple independent reticulation events on different background pairs. The Cauca SP-resistance pattern (*β*_1_ = 12 across multiple genomic backgrounds, Section 7.5) and the Pf7 Southeast Asian and Papuan populations (Section 7.4) sit here. Lewontin’s within-population structure made topological.

#### Temporal compounds: trajectories between primitives

A population at one moment occupies an alphabet category; over time the category may change. These trajectories are not new alphabet letters but are vocabulary built from the primitives:

- *Selective-sweep transient* (*K*_3_ → *K*_1_ trajectory): pre-sweep the population is in a recombinant regime; during the sweep, *β*_1_ rises as resistant haplotypes recombine into multiple ancestral backgrounds; after fixation, *β*_1_ collapses toward *K*_1_. The Cambodia artemisinin sweep (Section 7.6) traces this trajectory in real time.
- *Sympatric co-circulation* (*K*_2_ extended in time): two divergent lineages co-existing without reticulation produce *β*_0_ ≥ 2 stably with no *β*_1_. Distinguished from a single *K*_2_ snapshot only by temporal stability across multiple sampling windows. Conjectured; supported by synthetic simulation only.
- *Bottleneck + expansion* (*K*_3_ → *K*_1_ collapse with *β*_0_ spike): a recombinant population passes through a chokepoint and re-expands. The static topology at any time is one of the primitives; the temporal signature is the trajectory through them. Conjectured; we connect this to the Opqua simulation framework [3] as a target for follow-up validation.

#### Biogeographic compounds: the macroevolutionary *A* vocabulary

At the species-macroevolutionary scale, alphabet primitives are juxtaposed across geographic regions to give compound signatures:

- *Post-glacial expansion* (*A*_1_): geographically progressive instantiation of *K*_1_, with serial founder effects producing near-clonal populations along the expansion front. Validated by the United Kingdom *Arabidopsis* (*β*_1_*/n* = 0, Section 7.7).
- *Refugium-relict* (*A*_2_): geographically circumscribed instantiation of *K*_3_ at high *β*_1_*/n*, sometimes with *K*_4_ higher-order signal. Validated by the Iberian *Arabidopsis* (Spain *β*_1_*/n* = 0.64, *η* ≈ 0.40).

#### The alphabet inside real data

The four primitives appear as identifiable local motifs inside empirical Mapper graphs. Figure 2 illustrates this for the Guapi/Cauca *P. falciparum* Mapper (giant component: 42 nodes, 94 edges, 53 fundamental cycles). Inside this single graph we can locate a *K*_1_-like tail (a chain of low-degree nodes outside the cycle space), a *K*_2_-like flare (a 9-spoke hub, lineage divergence at one sampling time), a *K*_3_-like four-cycle (a single reticulation loop), and a *K*_3_-multi-origin figure-eight (two cycles sharing a vertex, the within-*K*_3_ multi-origin variant). The alphabet primitives are not abstractions; they are recurring local building blocks of the topology that real data produce.

**Figure 1:**
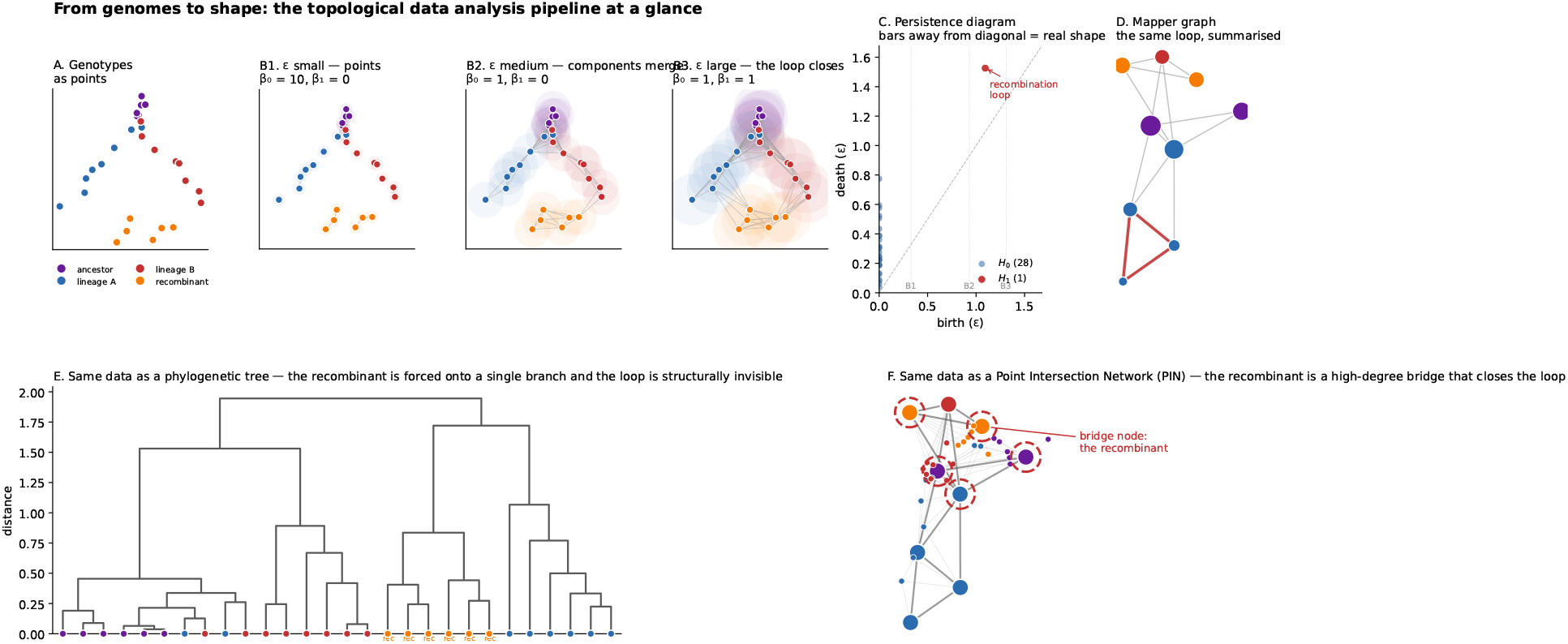
The TDA pipeline at a glance, on a synthetic dataset with a deliberate recombinant bridge between two parental lineages. **A:** the genotypes as points coloured by lineage. **BI-B3:** the Vietoris-Rips filtration at three growing radii *ε*. At small *ε* every genotype is isolated (*β*_0_ = 10); at intermediate *ε* components merge (*β*_0_ = 1, *β*_1_ = 0); at large *ε* the recombination loop closes (*β*_1_ = 1). : the persistence diagram records each topological feature as a (birth, death) point; the single *H*_1_ point far from the diagonal is the recombination loop. **D:** the Mapper 1-skeleton renders the same loop as a graph cycle (highlighted in red). **E:** the same data as a UPGMA phylogenetic tree␁the recombinant samples (orange dots, “rec”) are scattered across branches and the loop is structurally invisible. **F:** the same data as a Point Intersection Network␁the recombinant cluster appears as a high-degree bridge node (red dashed circle) that closes the loop the tree had to discard. The takeaway: topology is the part of the data the tree was hiding.

**Figure 2:**
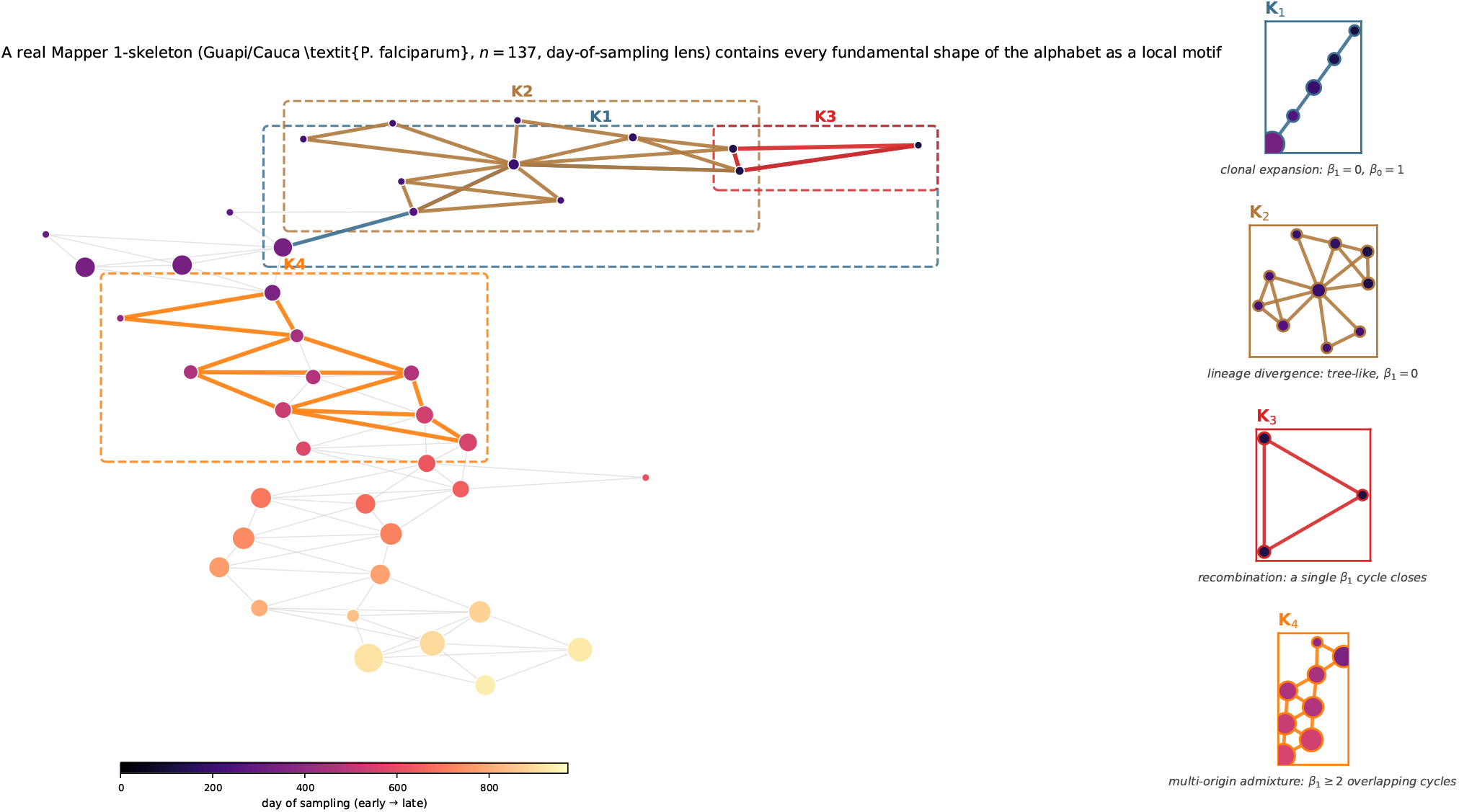
The alphabet as local motifs in a single empirical Mapper graph (Guapi/Cauca *P. falciparum, n* = 137 samples, day-of-sampling lens). The main panel shows the full giant component coloured by mean sampling day; the four right-hand zooms isolate sub-graphs that exemplify each fundamental shape—*K*_1_ (clonal tail), *K*_2_ (lineage-divergence flare), *K*_3_ (single recombination cycle), and a multi-origin *K*_3_ figure-eight (two cycles sharing a vertex, the high-*β*_1_ within-*K*_3_ variant). Dashed boxes in the main panel show where each motif sits in the graph.

## 6 The Topological Evolutionary Correspondence

We frame the central organising idea of this paper as a working hypothesis rather than a formal conjecture, because its evidence is genuinely asymmetric across the alphabet (some templates rigorously supported by simulation and empirical data, others theoretically motivated and not yet tested).

### Hypothesis I

(Topological Evolutionary Correspondence (TEC)). *Let* Θ = {*θ*_1_, …, *θ*_*K*_} *be evolutionary processes generating distance matrices D*_*θ*_. *Let ϕ*(*D*_*θ*_) ∈ ℝ^*p*^ *be the topological feature vector from Mapper graph statistics and persistent homology summaries. Then for sufficient sample sizes and biologically plausible parameters, there exists a classifier* 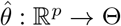 *with accuracy exceeding chance*.

### Remark I.

*The TEC is modular and its evidence is asymmetric. The non-reticulate alphabet (K*_1_, *K*_2_*) versus reticulate alphabet (K*_3_, *K*_4_*) distinction is sharp and supported by both simulation and theory: β*_1_ = 0 *for trees, β*_1_ *>* 0 *for reticulate histories. The K*_1_ *vs. K*_2_ *distinction requires graph features beyond persistent homology. The K*_3_ *vs. K*_4_ *distinction lives in the 2-skeleton: only K*_4_ *has β*_2_ *>* 0. *The temporal-compound vocabulary (selective sweeps, bottlenecks) and the biogeographic-compound vocabulary (A*_1_, *A*_2_*) inherit their evidential status from the alphabet primitives they are built from*.

### Remark 2.

*The TEC implies that topological features serve as summary statistics for Approximate Bayesian Computation [2]*. *The stability theorem [9] supports the continuity ABC needs at the level of the persistence diagram (bounded perturbations in D produce bounded changes in the diagram); the thresholded integer summaries we use are continuous only in the empirical, scale-swept sense of Section 7.9, and an ABC implementation should prefer diagram-level distances (e*.*g. bottleneck or Wasserstein) over raw counts where the guarantee matters*.

## 7 Results

### 7.1 Coalescent simulations distinguish reticulate from non-reticulate evolution

We simulated genomic data under four coalescent models (50 replicates, *n* = 50 haploid samples, *L* = 5,000 bp, *µ* = 10^−3^; 200 total observations): (M1) clonal: single panmictic population (*N*_*e*_ = 1,000), no recombination; (M2) divergence: two populations (*N*_*e*_ = 500 each) split at *t* = 500 generations, no recombination or migration; (M3) low recombination: split populations with migration *m* = 0.002 and recombination *ρ* = 5 *×* 10^−4^; (M4) high recombination: *m* = 0.02, *ρ* = 5 *×* 10^−2^. For each replicate, we computed the Hamming distance matrix, extracted persistent homology features (number of *β*_1_ bars |ℬ_1_|, total persistence Π_1_, maximum persistence 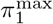, mean persistence 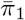) via Ripser [45], and graph-theoretic features from the 15th-percentile threshold graph (connected components *β*_0_, cycle rank *γ*, mean and maximum degree, clustering coefficient *C*, diameter).

The results confirm the template catalog (Table 2, Figure 3) and, just as importantly, make the sampling baseline explicit. The non-reticulate models do *not* produce *β*_1_ ≈0: clonal evolution (*ρ* = 0) yields 13.4 ± 3.4 bars and divergence 14.7 ± 4.2. This is the finite-sample geometric pedestal of the Vietoris-Rips complex, not recombination signal—and it is precisely why we read *β*_1_ as a quantity elevated *above* a baseline rather than as an event count. The number of bars then increases monotonically with recombination rate: low recombination (55.0 ± 6.1), high recombination (78.3 ± 6.4). The separation between non-reticulate and reticulate models is nonetheless sharp: the highest single-replicate |ℬ_1_| for any non-reticulate simulation (divergence) is lower than the lowest for any reticulate simulation (low recombination), so the reticulate/non-reticulate boundary is cleanly resolvable even though the baseline is large. Total persistence Π_1_ follows the same monotonic pattern above baseline: clonal (0.034 ± 0.012), divergence (0.037 ± 0.014), low recombination (0.154 ± 0.026), high recombination (0.228 ± 0.028). Graph features provide complementary discriminative power: clustering coefficient *C* drops sharply from non-reticulate models (*C* = 0.962 clonal, 0.949 divergence) to reticulate models (*C* = 0.412 low recombination, 0.258 high recombination). The threshold-graph connected-component counts *β*_0_ (9.4, 8.8, 2.2, 1.3) are reported for completeness but do *not* separate clonal from divergence (9.4 vs 8.8): at this sample size and threshold the two share an indistinguishable *β*_0_, and what distinguishes divergence is the *H*_0_ *persistence* structure (multiple finite-death bars; Section 5.1), not the component count.

**Table 2:** Topological and graph features across coalescent models (mean ± SD, 50 replicates × 50 samples).

| Model | $\vert \mathcal{B}_1\vert $ | $\Pi_1$ | $\pi_1^{\max}$ | $\beta_0$ | $C$ | Diameter |
| --- | --- | --- | --- | --- | --- | --- |
| Clonal | $13.4 \pm 3.4$ | $0.034 \pm 0.012$ | $0.007 \pm 0.002$ | $9.4 \pm 2.0$ | $0.962 \pm 0.025$ | $1.8 \pm 0.8$ |
| Divergence | $14.7 \pm 4.2$ | $0.037 \pm 0.014$ | $0.007 \pm 0.002$ | $8.8 \pm 2.2$ | $0.949 \pm 0.037$ | $2.2 \pm 1.2$ |
| Low recomb. | $55.0 \pm 6.1$ | $0.154 \pm 0.026$ | $0.008 \pm 0.001$ | $2.2 \pm 0.5$ | $0.412 \pm 0.036$ | $3.2 \pm 0.4$ |
| High recomb. | $78.3 \pm 6.4$ | $0.228 \pm 0.028$ | $0.009 \pm 0.001$ | $1.3 \pm 0.5$ | $0.258 \pm 0.034$ | $4.1 \pm 0.6$ |

**Figure 3:**
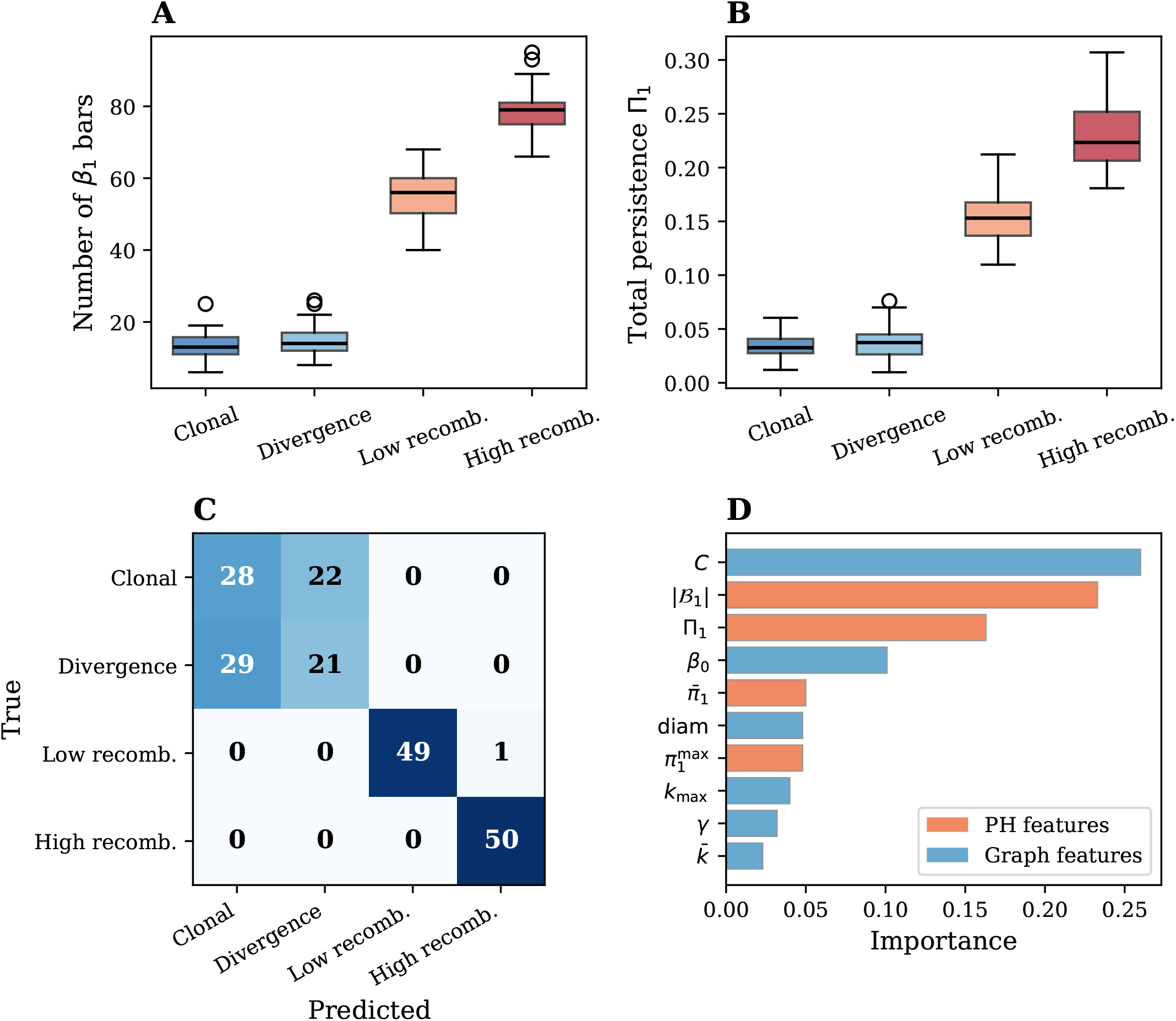
Coalescent simulation results. **A:** Distribution of *β*_1_ bar counts across 50 replicates per model; reticulate models produce 4-6× more 1-cycles than non-reticulate models. **B:** Total persistence Π_1_, quantifying cumulative topological complexity. **C:** Confusion matrix (normalized) for the 4-class random forest classifier; confusion is confined to the clonal-divergence pair, while reticulate models are classified with ≥ 98% recall. **D:** Feature importances; clustering coefficient *C* and number of *β*_1_ bars are the two most discriminative features, with PH features (orange) and graph features (blue) contributing complementary information.

A random forest classifier (500 trees, 5-fold stratified cross-validation) on the full 10-dimensional feature vector *ϕ* achieves 74.0% ± 4.1% accuracy (Figure 3C). The structure of the confusion matrix is more informative than the overall accuracy: *not a single reticulate simulation was misclassified as non-reticulate, nor vice versa*. Low recombination is classified with 98% recall (49/50 correct) and high recombination with 100% recall (50/50). All classification error concentrates in the clonal-divergence pair (51 misclassifications out of 100), which share nearly identical topological profiles (|ℬ_1_| ≈14, *C* ≈0.95). This confirms that the reticulate/non-reticulate boundary is topologically sharp and robust, while the distinction between tree topologies (path vs. branching) requires additional structural features.

Strikingly, persistent homology features alone achieve 74.5% ± 3.7% accuracy—slightly exceeding the full feature set—while graph features alone achieve 72.0% ± 2.9% (Figure 6) The top-ranked features by importance are clustering coefficient (*C*, 26.0%), number of *β*_1_ bars (23.3%), and total persistence Π_1_ (16.3%), demonstrating that PH and graph features contribute complementary discriminative power.

#### The classifier’s errors match the alphabet, not a random-confusion pattern

Figure 4 reframes the classification experiment in terms of the template alphabet. Effect sizes (Cohen’s *d*) between contrasts confirm that the topological features split classes *by kind and by dose*, not uniformly: the *K*_1_ → *K*_3_ topology jump gives *d* = 8.46 on |ℬ_1_| and *d* = 6.00 on Π_1_; the dose-response between low and high recombination along *K*_3_ gives *d* = 3.74 on |ℬ_1_| ; but the clonal/divergence contrast—which the alphabet predicts should share topology—gives *d* = 0.35, essentially indistinguishable. The classifier’s only systematic errors collapse exactly the pair the alphabet predicts are topologically equivalent.

**Figure 4:**
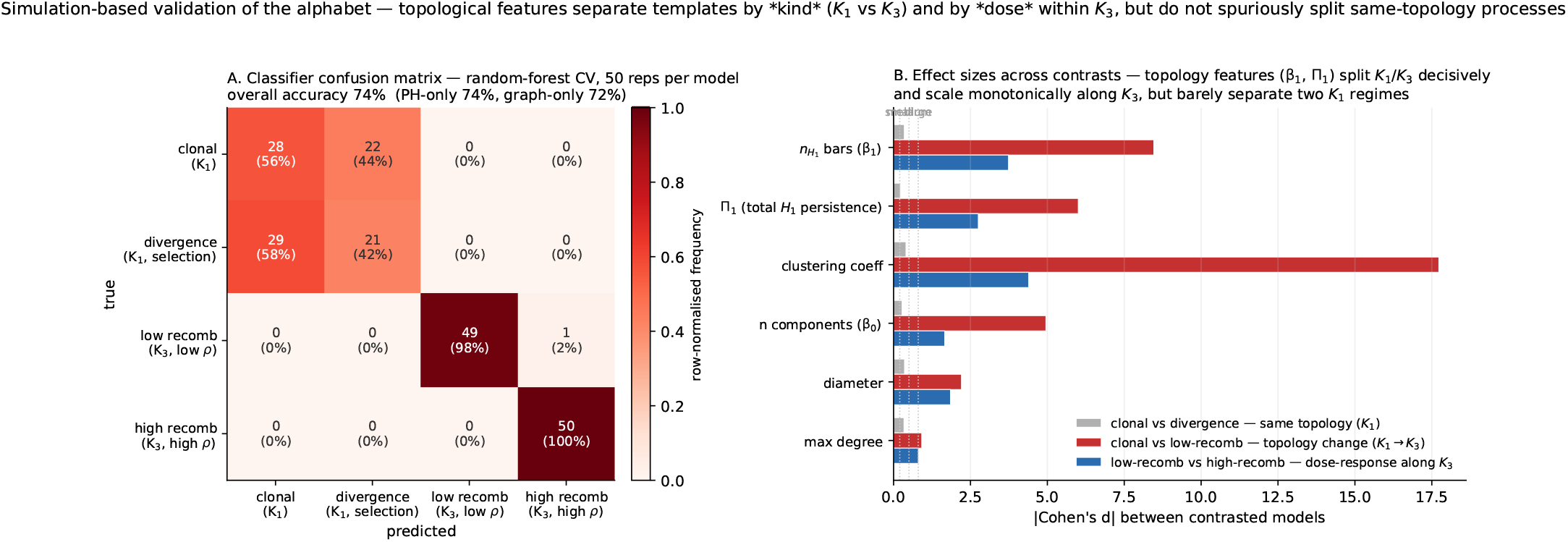
Alphabet-aligned view of the simulation experiment. **A:** Row-normalised confusion matrix (50 cross-validated reps per model), grouped to show that reticulate templates (*K*_3_) are never confused with non-reticulate templates (*K*_1_). **B:** Effect sizes (|Cohen’s *d*|) across three contrasts. Topology-jump contrasts (*K*_1_ → *K*_3_, red) produce large effects on *β*_1_ and Π_1_; the *K*_3_ dose-response (blue) is still large on *β*_1_; the same-topology contrast (grey) produces no large effect on any feature. The pattern is consistent with the framework’s prediction that topology is the invariant that discriminates templates.

**Figure 5:**
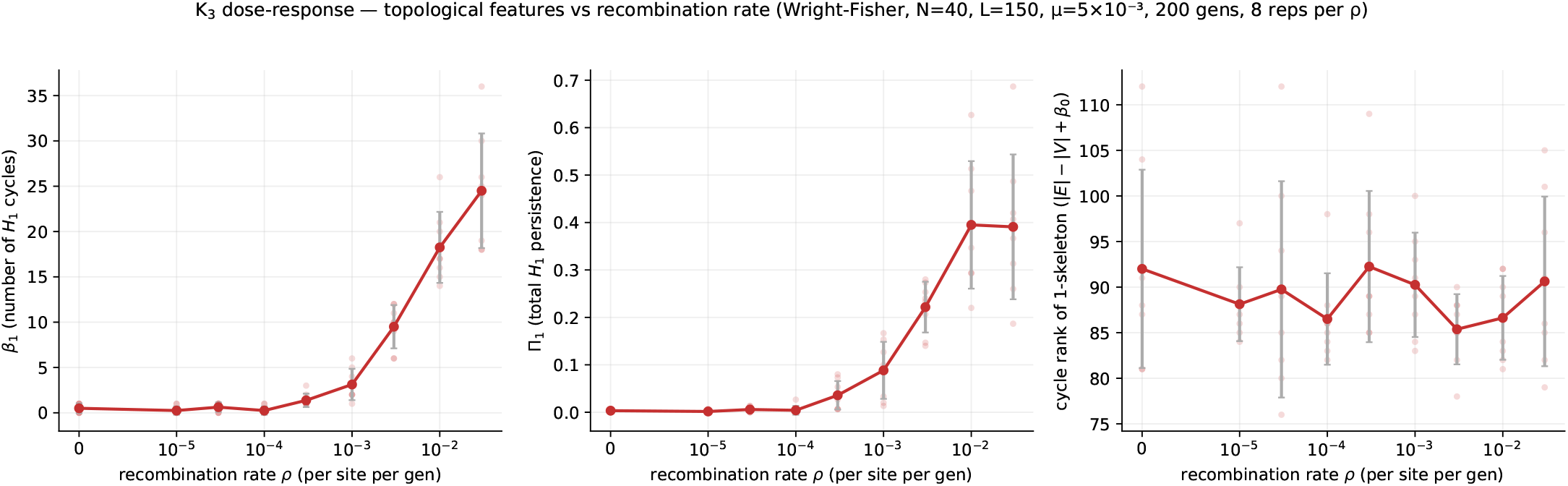
*K*_3_ dose-response: topological features vs. recombination rate under a Wright-Fisher forward simulator. Error bars are ± 1 SD across 8 replicates per *ρ*; individual replicates are shown as translucent dots. *β*_1_ and Π_1_ are monotone nondecreasing in *ρ* over six orders of magnitude; a fixed-percentile graph cycle rank is essentially flat—the signal is in the persistent homology, not the static graph.

**Figure 6:**
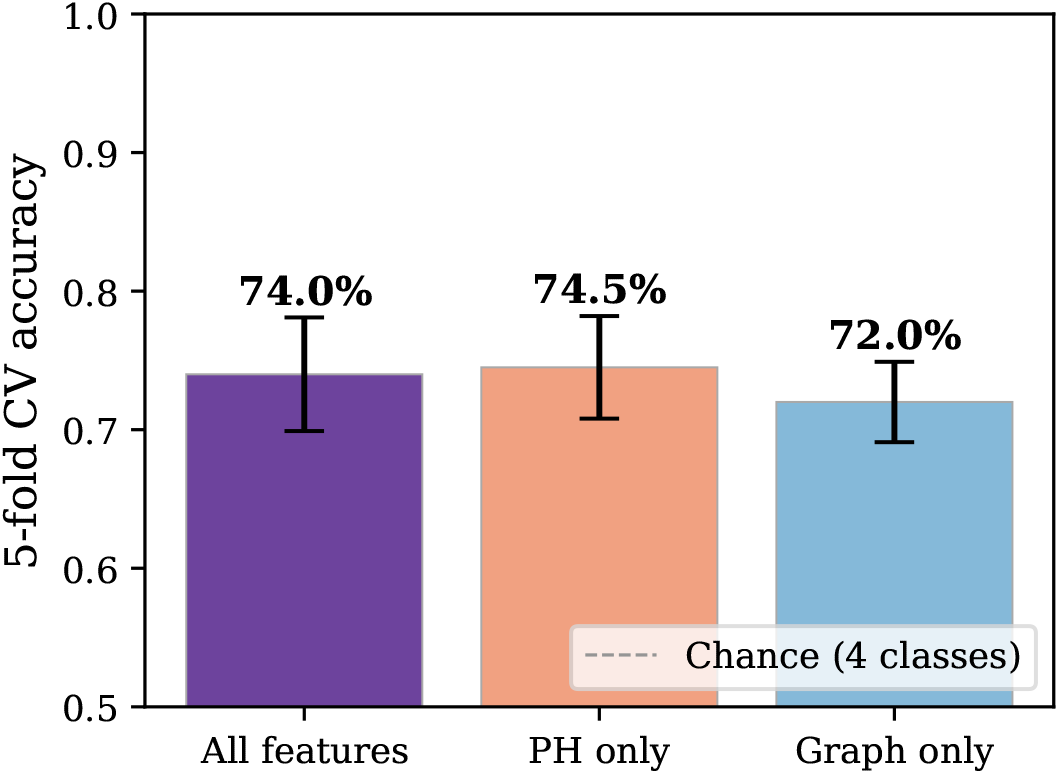
Five-fold cross-validated classification accuracy using all features, persistent homology (PH) features only, and graph features only. Dashed line indicates chance level (25%). Error bars show ±1 SD across folds.

**Figure 7:**
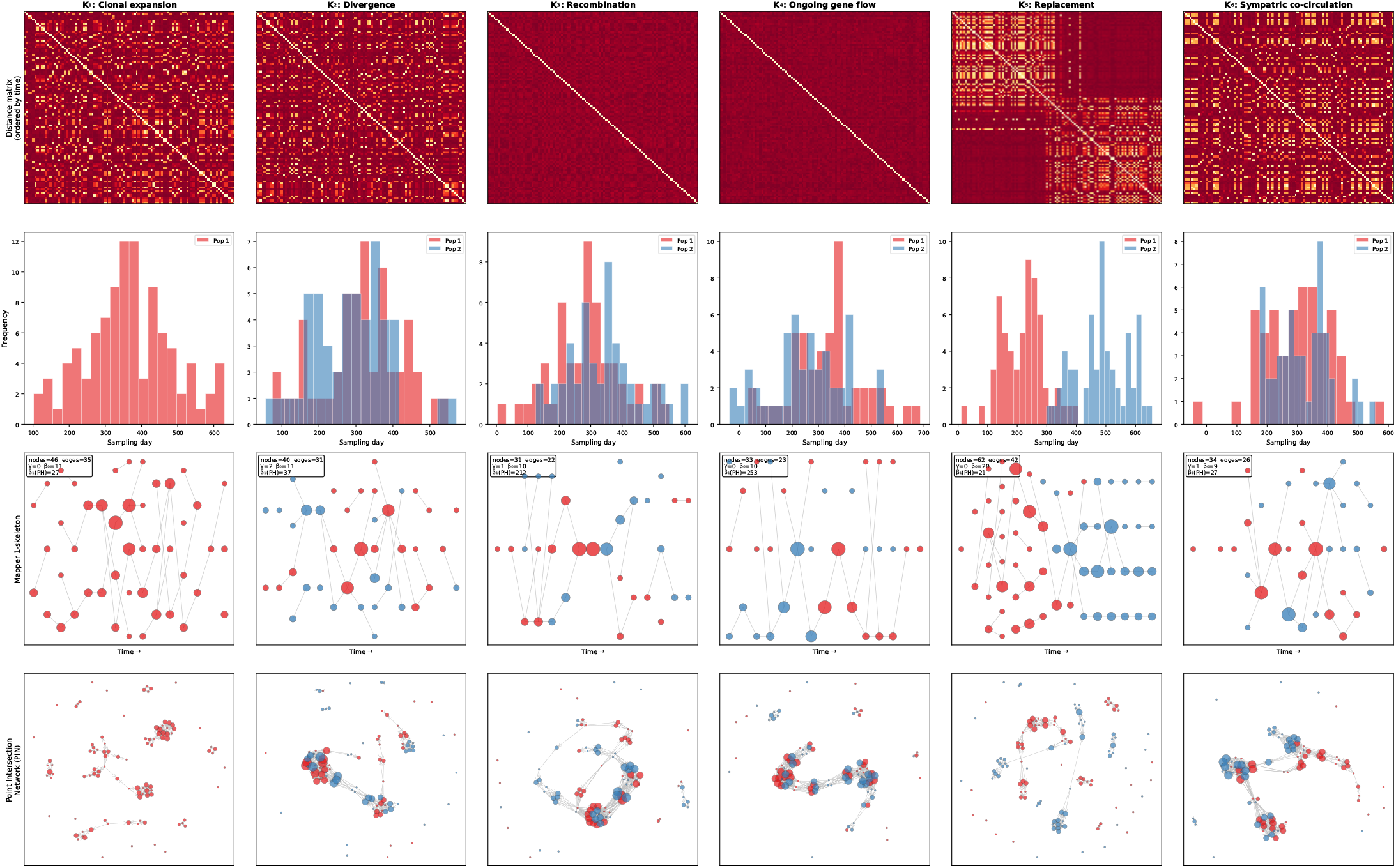
Coalescent simulations visualized through the dual TDA framework. Each column represents one evolutionary scenario simulated under a proper coalescent model. **Row 1:** amming distance matrix (samples ordered by time); block structure emerges naturally from the evolutionary process. **Row 2:** Temporal histogram of sampling dates colored by population. **Row 3:** Mapper 1-skeleton with nodes colored by dominant population; the shape of each graph corresponds to one of the alphabet primitives or compound vocabulary entries. **Row 4:** Point Intersection Network (PIN) with nodes colored by population; high-centrality nodes identify bridging genotypes. *Note on column labels*: the first four columns are the alphabet primitives *K*_1_-*K*_4_; the final two columns are the replacement and sympatric-co-circulation scenarios (R and C in the text)—temporal-compound vocabulary, not additional alphabet letters. (Any “K_5_/ K_6_” rendered in the panel image refers to these two scenarios and is being relabelled R/C for consistency.)

#### Dose-response alon *K*_3_

The *K*_3_ template predicts not a discrete state but a monotone dose-response: *β*_1_ and Π_1_ should increase with recombination rate *ρ* over the coalescent-with-recombination regime. Figure 5 tests this directly with a Wright—Fisher forward simulator (*N* = 40 haplotypes, *L* = 150 sites, *µ* = 5 × 10^−3^, 200 generations, 8 replicates per *ρ*) at nine values of *ρ* spanning six orders of magnitude. The mean number of *β*_1_ bars rises from 0.5 at *ρ* = 0 to 24.5 at *ρ* = 3 × 10^−2^, and total persistence Π_1_ from 0.003 to 0.40. The curves are monotone nondecreasing, as predicted. A fixed-percentile neighbourhood-graph cycle rank does not track the signal, confirming that the *K*_3_ dose-response lives in the persistent-homology readout and is not an artifact of graph density at any one scale.

### 7.2 Mapper 1-skeletons reveal the shape of each evolutionary process

To demonstrate that the template catalog is not merely a theoretical construct but produces visually distinct and interpretable shapes, we simulated genomic data under six evolutionary scenarios using the same coalescent framework (Section 7.1) and applied the full Mapper pipeline with sampling time as the filter function (Figure 7).

For each scenario, we generated *n* = 100–120 haploid samples (*L* = 3,000 bp, *µ* = 10^−3^) with temporal metadata reflecting the epidemiological context. Four scenarios instantiate the alphabet primitives: (*K*_1_) clonal expansion in a single population with serial sampling over an epidemic wave; (*K*_2_) two populations diverging without gene flow, both sampled throughout; (*K*_3_) two populations with low recombination (*ρ* = 5 × 10^−4^) and migration (*m* = 0.005); (*K*_4_) high recombination (*ρ* = 5 × 10^−2^) and migration (*m* = 0.02).

Two further scenarios illustrate temporal-compound *vocabulary* rather than alphabet letters; we label them by name to keep them distinct from the four primitives: scenario R, lineage *replacement*, where an ancestral population is displaced by a genetically distant invading population with non-overlapping temporal sampling; and scenario C, sympatric *co-circulation*, with the same genetic divergence as R but simultaneous sampling of both lineages.

The Mapper 1-skeletons, computed with adaptive clustering height (15 intervals, 40% overlap), show strikingly different shapes for each scenario. In K_1_ (clonal), the 1-skeleton is a linear chain—nodes are connected sequentially along the time axis with no loops (*γ* = 0). In _2_ (divergence), the two populations form spatially separated flares that branch from a common temporal origin. In K_3_ and K_4_ (recombination), nodes from both populations intermix and form connected structures with cycles, reflecting genotypes that bridge the parental clusters. The cycle rank *γ* and PH-derived 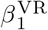 counts increase sharply (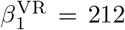 for low recombination, 253 for high recombination, vs. 27-37 for non-reticulate models; these are the larger *n* = 100–120 illustrative simulations, distinct from the *n* = 50 classifier simulations of Table 2).

Critically, scenario R (replacement) and scenario C (sympatric co-circulation) demonstrate that the temporal filter function mediates the Mapper output: the *same* genetic distance structure produces different 1-skeletons depending on whether the populations are sampled sequentially or simultaneously. In R, the 1-skeleton shows a temporal transition—red (ancestral) nodes on the left, blue (invading) on the right—making the replacement event visible as a progression along the time axis. In C, the populations are interleaved in time, producing mixed-color clusters throughout the 1-skeleton.

The Point Intersection Networks (bottom row, Figure 7) provide the individual-level view: nodes with high intersection centrality are the genotypes that bridge distinct Mapper clusters. In the recombination scenarios (*K*_3_, *K*_4_), the PIN shows multi-colored hubs—the recombinant genotypes connecting parental populations. In the replacement scenario R, bridging nodes are sparse and confined to the transition period.

### 7.3 Persistent homology confirms extensive recombination in *P. falciparum*

We computed persistent homology on the 137 × 137 IBD distance matrix from the Guapi *P. falciparum* dataset [34]. The Vietoris–Rips filtration produces 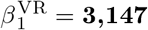 **persistent 1-cycles** (total bar count over the filtration), with total persistence Π_1_ = 172.99, maximum bar persistence 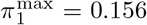, and 2,857 bars with persistence exceeding 0.01.

This result is orders of magnitude more topologically complex than any simulated scenario: even the high-recombination coalescent model produces 78.3 ± 6.4 *β*_1_ bars at *n* = 50 samples and *L* = 5,000 bp (Table 2), roughly 40× fewer than the empirical observation. The Guapi data, from a region of moderate transmission intensity where Carrasquilla et al. [6] documented both clonal persistence and selective sweeps, shows that the topological complexity of a real parasite population vastly exceeds even aggressively recombining coalescent models—precisely what the high-*β*_1_ multi-origin variant of *K*_3_ predicts for populations with sustained, multilocus outcrossing across genetically divergent strains.

This result is consistent with the PIN analysis from Knudson et al. [34], where bridging genotypes connecting STRUCTURE-defined subpopulations were identified. The persistent homology provides a global summary: 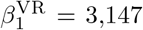 one-cycles, a measure of reticulate structure far above any clonal baseline that a phylogenetic tree cannot capture. We are deliberate about not overstating the relationship between this number and the Mapper picture. The Mapper nerve of the same data has a cycle rank 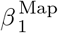 of order tens (Section 7.11), *not* thousands; the two are different objects (point-cloud persistent homology versus the first Betti number of a clustered graph) and we do not claim the larger number “validates” the smaller. What the PIN and the persistent homology agree on is qualitative and that is all we assert: both identify the Guapi population as densely reticulate, with bridging genotypes drawn from multiple genetic backgrounds. A quantitative concordance between 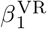 and 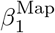 would require matching the Mapper cover to the persistent-homology scale, which we have not done and leave as an open methodological question.

For the companion Pf7 regional analysis (Section 7.4) we establish statistical significance through a label-shuffle null: ten independent 200-sample draws from the full QC-pass pool, ignoring population, yield a *β*_1_ distribution ranging [8, 32] with median 20.5. The observed per-population values deviate significantly in both directions; five of ten populations fall outside the null band. The Guapi-specific null remains to be completed, but the Pf7 regional null establishes that geographic structure generates *β*_1_ patterns that are not explainable by the marginal distance distribution alone.

### 7.4 Cross-regional validation: the MalariaGEN Pf7 dataset

To test whether topological complexity varies across *P. falciparum* populations in the manner predicted by their different recombination regimes, we analyzed the MalariaGEN Pf7 genetic distance matrix [55], which contains 20,864 samples from 33 countries (16,203 QC-pass). For each of ten WHO-defined regional populations we drew 200 QC-pass samples at random (seed 7) and computed the Vietoris-Rips persistent homology of the induced IBD distance submatrix. A robustness sweep across *n* ∈ {100, 200, 400} and seeds {7, 42, 101} confirms that the qualitative pattern is stable to subsampling (Figure 9).

The observed per-population *β*_1_ values span two orders of magnitude (Figure 8): low in Africa (AF -W = 6, AF-C = 3, AF-E = 3, AF-NE = 27), elevated in mainland South America (SA = 31) and far-eastern South Asia (AS-S-FE = 22), and strongly elevated in Southeast Asia and Oceania (AS-SE-E = 33, AS-SE-W = 61, OC-NG = 89). One Asian population, AS-S-E, yields *β*_1_ = 0. This ordering runs opposite to the naive recombination-rate prediction—under which high-transmission African populations would exhibit the richest reticulate topology—and instead tracks the geography of clonal population substructure: Southeast Asia has undergone intense selective sweeps (see Section 7.6) that created multiple persistent resistant lineages carried on partially-differentiated backgrounds; Papua New Guinea has long-standing demographic isolation; and the most panmictic African populations carry such uniformly high pairwise similarity that cycles are filled by triangles at the earliest filtration scales (the saturation regime). The topological invariant is therefore diagnostic not of recombination rate per se but of the *balance* between outcrossing and the demographic structuring of genotypes—a more discriminating evolutionary observable.

**Figure 8:**
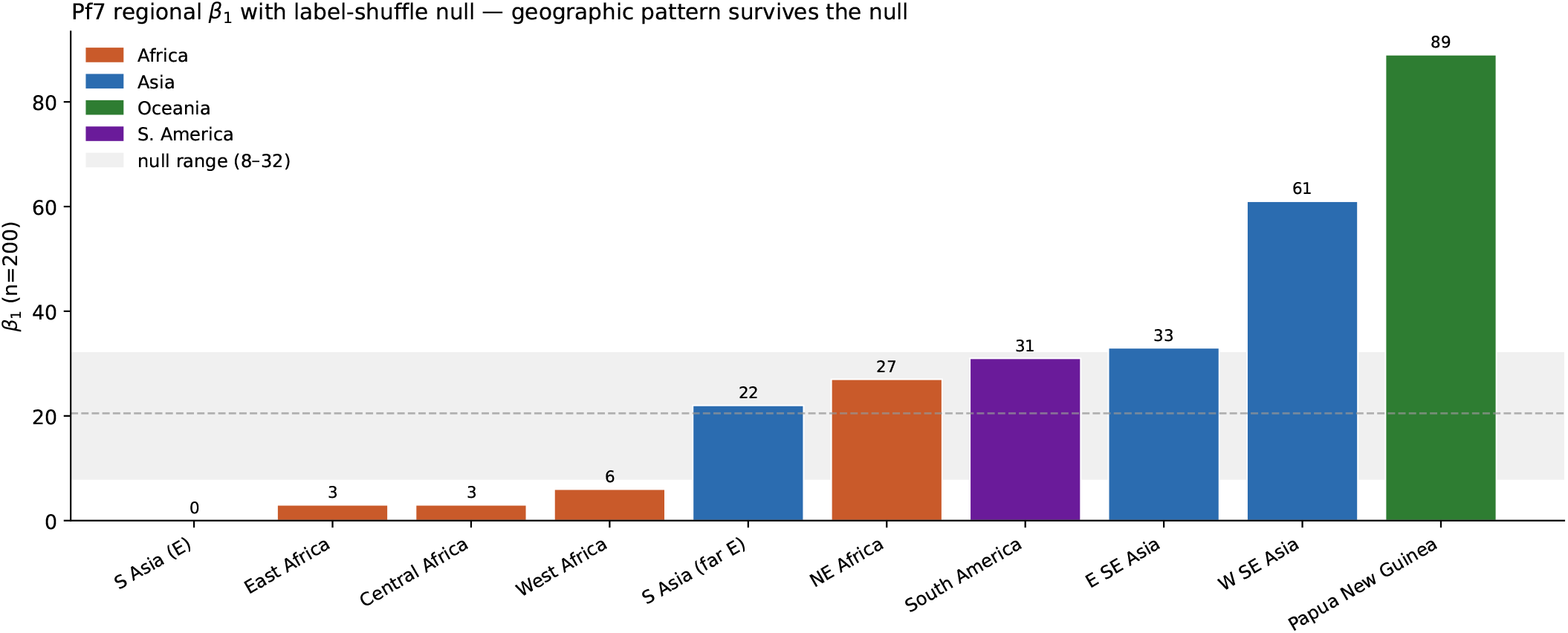
Per-population *β*_1_ on the MalariaGEN Pf7 IBD distance matrix (*n* = 200 per population, seed 7). The shaded band is the [8, 32] range from ten label-shuffle null draws (dashed line: null median 20.5). Five of ten populations fall outside the null band. African populations trend below (panmictic saturation in AF -W/AF -E/AF -C), Southeast Asia and Papua New Guinea trend above (multi-lineage structure from post-sweep diversification).

**Figure 9:**
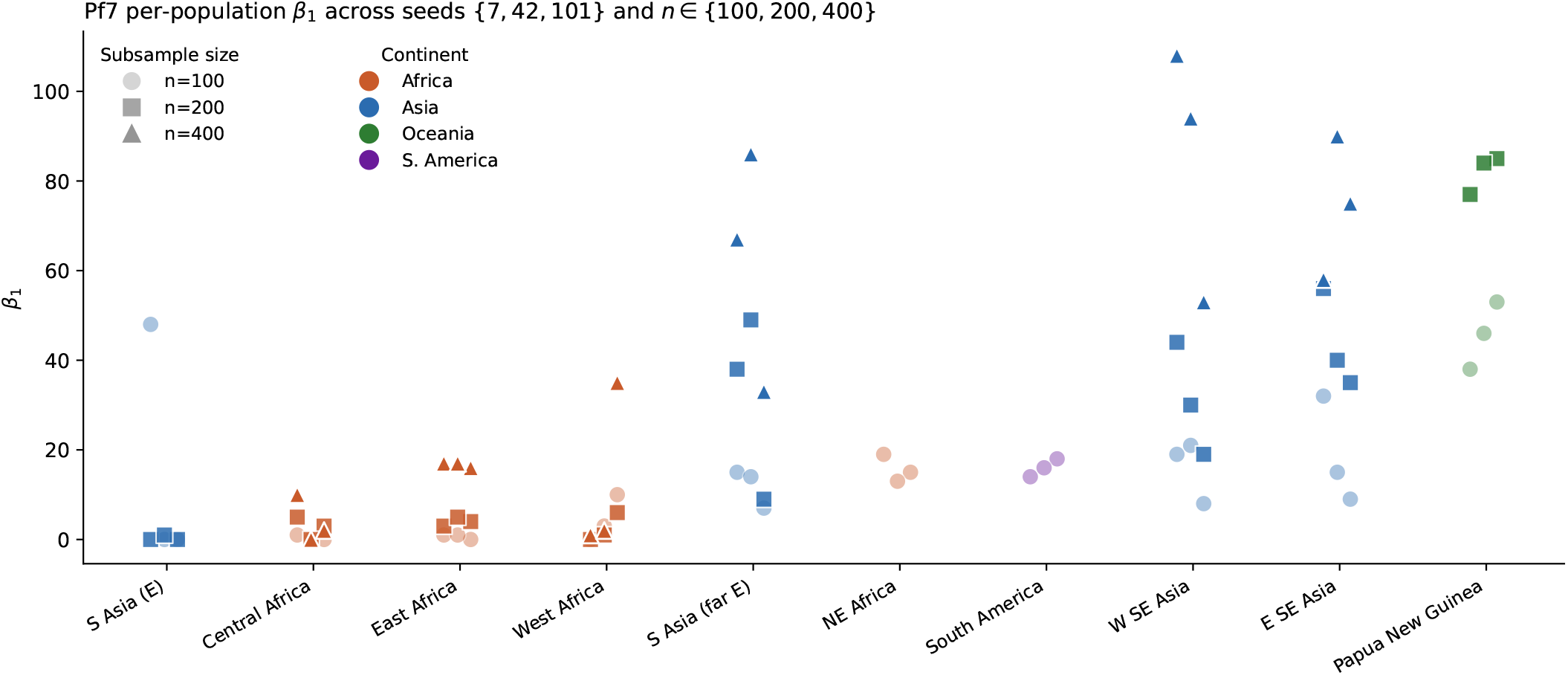
Robustness of the regional *β*_1_ pattern across subsample sizes *n* ∈ {100, 200, 400} (markers) and random seeds {7, 42, 101} (three replicates per cell) for each of ten Pf7 populations. The per-population *β*_1_ rankings are preserved across subsample sizes: the observed pattern is not a small-*n* artifact.

#### Reconciling the simulation dose-response with the empiriscal ordering

This is the central interpretive point of the paper and we state it plainly, because the simulation result (Figure 5: *β*_1_ increases with *ρ*) and the empirical result (Figure 8: *β*_1_ is *lowest* in the freely-recombining African populations) can be misread as contradictory. They are not, because they hold different things fixed. In the Wright–Fisher and coalescent simulations we vary *ρ* while holding demographic structure approximately constant; there, *β*_1_ is a clean monotone function of *ρ* above the sampling baseline. In the Pf7 data *ρ* is *uncontrolled* and demographic structure varies by orders of magnitude across populations; there, *β*_1_ is dominated by structure—the geometry of how genotypes are organized—rather than by the instantaneous recombination rate. The two regimes are linked by the saturation mechanism: in a highly panmictic, freely-recombining population, pair-wise similarity is so uniform that cycles are filled by triangles at the earliest filtration scales and *β*_1_ *falls* even though recombination is high, whereas a structured, multi-lineage population leaves many unfilled cycles and *β*_1_ *rises*. Both figures are therefore measuring the same quantity—non-tree-like organization of genetic-distance space above a sampling baseline—and the apparent sign flip between them is the expected consequence of whether recombination is operating within a single well-mixed cloud (saturation, low *β*_1_) or across persistently differentiated backgrounds (multi-lineage structure, high *β*_1_). The clean dose-response of Figure 5 describes the controlled regime; the empirical ordering of Figure 8 describes a regime in which population structure is the dominant axis of variation. We do not claim a single scalar reads recombination rate directly from any genetic-distance matrix; we claim *β*_1_ reads reticulate organization, which coincides with recombination rate only when structure is held fixed.

To establish that this ordering is not an artifact of sample composition we constructed a label-shuffle null: ten independent 200-sample draws from the full QC-pass pool, ignoring population. The null *β*_1_ distribution ranges [8, 32] with median 20.5. Five of ten real populations fall *outside* this null band (Figure 8): AF-W, AF-C, AF-E, AS-S-E below; AS-SE-W and OC-NG above. The geographic signal is real.

#### In plain terms

The topology of a malaria population’s genetic data is a single coordinate-free summary of its reticulation history: the cumulative effect of transmission intensity, multiplicity of infection, and outcrossing rate, written as a number rather than inferred from a model. Two populations with similar nucleotide diversity can sit at opposite ends of the *β*_1_ scale because they differ in how their variation is *organized*, not in how much of it there is. That organization—deep-coalescent multi-lineage structure versus panmictic saturation—is what the topological invariant reads out, and what classical summary statistics like *F*_ST_ are designed not to see.

### 7.5 Multi-background resistance: Colombia Cauca SP-resistance topology

Corredor et al. [11] demonstrated that sulfadoxine-pyrimethamine (SP) resistance in Colombian *P. falciparum* spread across the Andes from a single mutational origin. If that process had been a clean clonal sweep, resistant samples today would form a single near-clonal clump and the topological *β*_1_ of the resistant subset would match the sensitive baseline. The Pf7 Colombian samples allow a direct test.

We extracted all 135 QC-pass Colombian samples and partitioned them by their inferred sulfadoxine and pyrimethamine phenotype [55]. Not every sample receives a confident call on both axes: of the 135, three lack a confident sulfadoxine call (so the sulfadoxine-typed marginals are 82 + 50 = 132) and four lack a confident pyrimethamine call (pyrimethamine-typed marginals 76 + 55 = 131); the joint cells use only doubly-typed samples. The full-cohort persistent homology yields 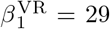 with Π_1_ = 6.3 × 10^−4^. Split by SP status, the two subsets diverge sharply (Table 3). Jointly resistant samples (sulfadoxine-R ∧ pyrimethamine-R, *n* = 62) carry 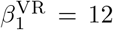 and Π_1_ = 1.1 × 10^−4^. Jointly sensitive samples (*n* = 36) carry 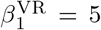 and Π_1_ = 2.9 × 10^−6^—two orders of magnitude less persistent. (The joint subsets 62 + 36 = 98 do not exhaust the cohort: the remaining samples are phenotype-discordant across the two drugs or untyped on one axis.)

**Table 3:** Per-condition persistent homology on the Colombian Cauca *P. falciparum* subset of Pf7 (*n* = 135 QC-pass). Single-drug marginals do not sum to 135 because three samples lack a confident sulfadoxine call and four lack a confident pyrimethamine call; the joint cells (Sulf-R ∧ Pyr-R, Sulf-S ∧ Pyr-S) use only doubly-typed samples and need not sum to the cohort. Resistant subsets carry more than two orders of magnitude more total *H*_1_ persistence than the jointly sensitive subset—the high-*β*_1_, multi-origin band of *K*_3_, consistent with resistance carried on several distinct genomic backgrounds rather than a single clonal lineage.

| Condition | $n$ | $\beta_1$ | $\Pi_1$ | mean pairwise $d$ |
| --- | --- | --- | --- | --- |
| All Cauca | 135 | 29 | $6.27 \times 10^{-4}$ | — |
| Sulfadoxine-R | 82 | 15 | $1.76 \times 10^{-4}$ | — |
| Sulfadoxine-S | 50 | 9 | $8.89 \times 10^{-5}$ | — |
| Pyrimethamine-R | 76 | 17 | $2.49 \times 10^{-4}$ | — |
| Pyrimethamine-S | 55 | 7 | $7.09 \times 10^{-5}$ | — |
| Sulf-R $\wedge$ Pyr-R | 62 | 12 | $1.10 \times 10^{-4}$ | — |
| Sulf-S $\wedge$ Pyr-S | 36 | 5 | $2.94 \times 10^{-6}$ | — |

The shape itself is shown in Figure 10. The full cohort exhibits a hub-and-arm topology: a dense near-clonal core with several radial clumps connected by bridges, exactly what *β*_1_ = 29 reticulate structure looks like in two dimensions. The jointly resistant subset preserves a reduced version of this architecture with one dominant *H*_1_ class at persistence ≈8 × 10^−4^—a single very-long-lived cycle spanning distinct genomic backgrounds that all carry the resistant phenotype. The jointly sensitive subset collapses to one tight clump with *H*_1_ points hugging the diagonal.

**Figure 10:**
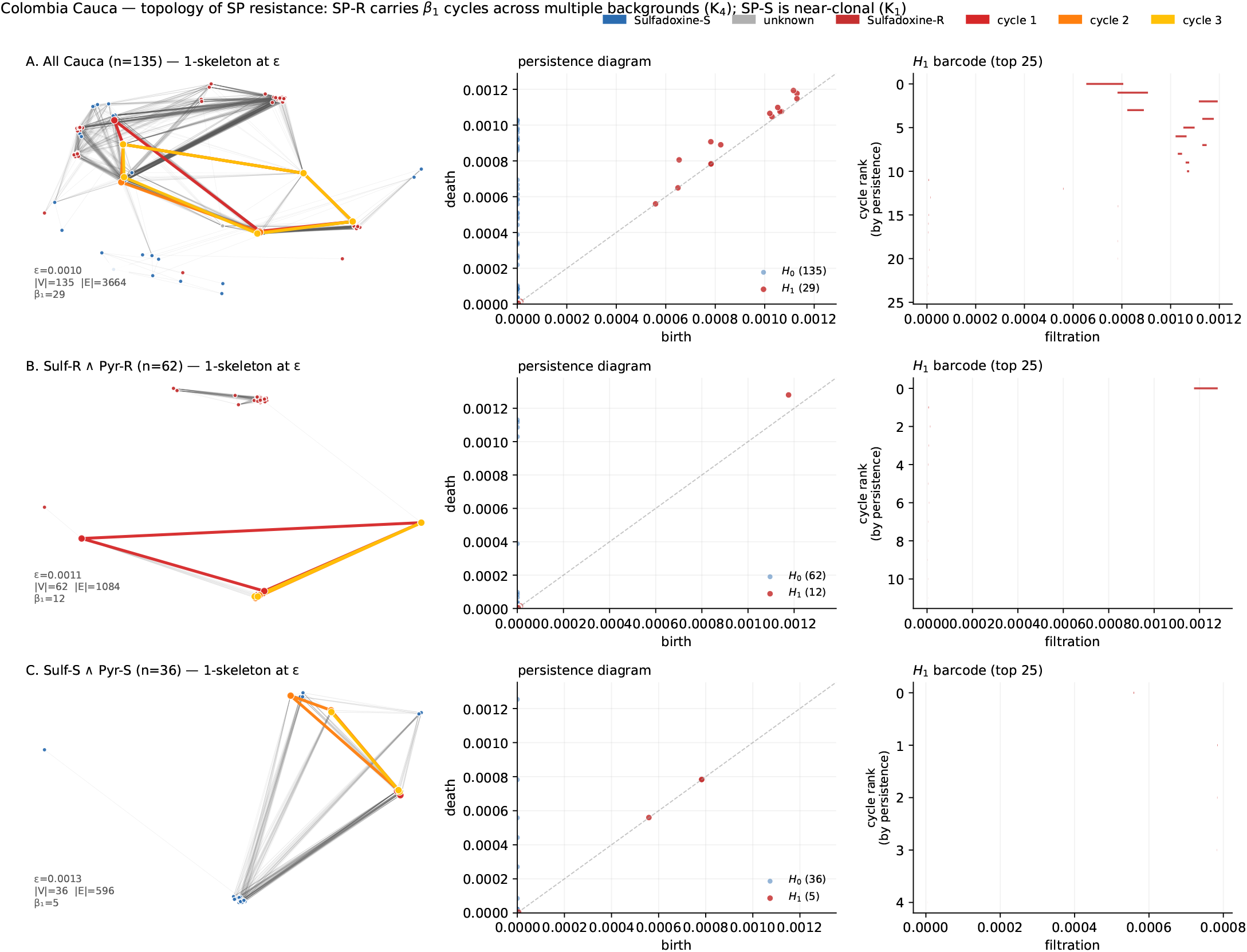
Topology of SP resistance in Colombian Cauca *P. falciparum* (Pf7, *n* = 135). Rows: all Cauca (A), jointly SP-resistant (B), jointly SP-sensitive (C). Columns: Vietoris-Rips 1-skeleton at the filtration level at which the space just becomes connected (spring-laid-out so graph topology is visible; nodes colored by sulfadoxine status); persistence diagram (*H*_0_ blue, *H*_1_ red); *H*_1_ barcode (top 25 cycles by persistence). The SP-resistant subset preserves a multi-clump architecture with one very-persistent cycle; the SP-sensitive subset is near-clonal. Template interpretation: multi-origin K_3_ (high-*β*_1_ variant; recurrent multi-background adoption of resistance alleles) for SP-R against a *K*_1_ baseline for SP-S .

**Figure 11:**
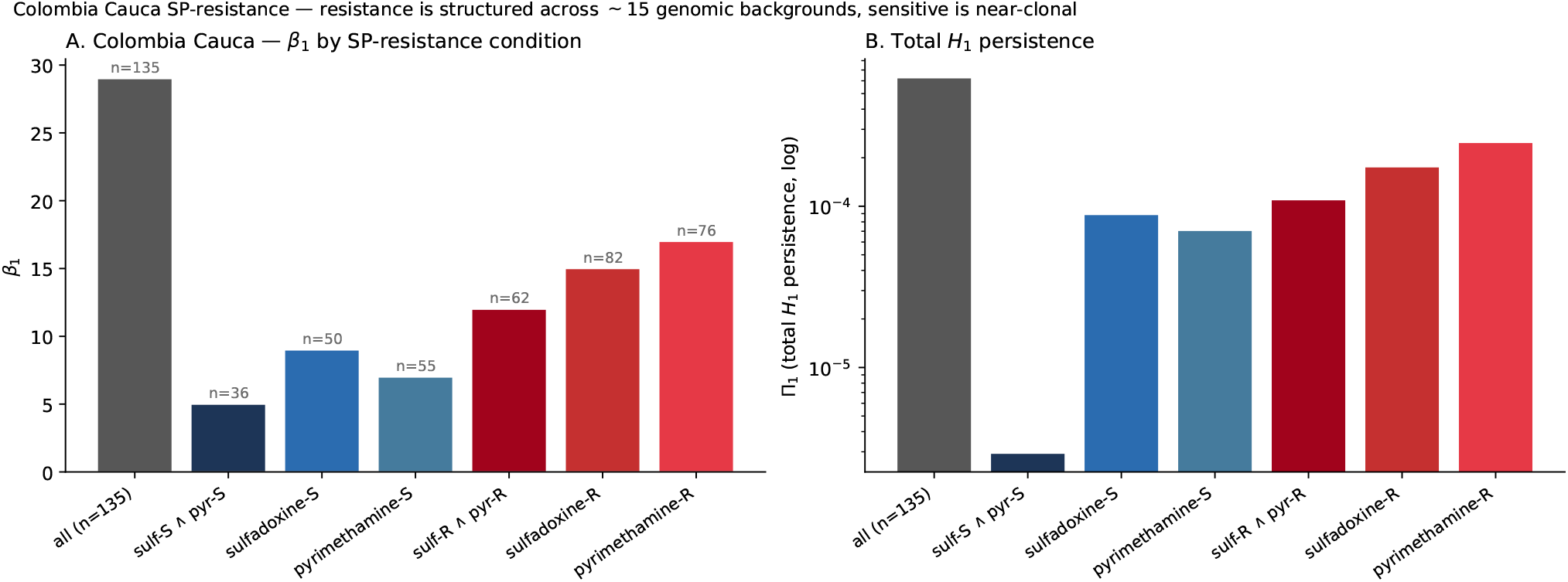
Colombian Cauca SP-resistance summary: *β*_1_ (left panel) and total *H*_1_ persistence Π_1_ (right panel, log scale) by SP condition. The SP-sensitive double (sulf-S ∧ pyr-S) carries Π_1_ ≈ 3 × 10^−6^; resistant subsets exceed this by two orders of magnitude.

The evolutionary reading: rather than a single resistant haplotype expanding clonally, SP resistance in Cauca sits in the high-*β*_1_, multi-origin band of *K*_3_, with 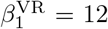 against a near-clonal sensitive baseline of 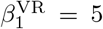. We are careful not to over-read the integer: 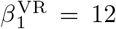 is not a count of twelve genomic backgrounds-the cycle count is not in one-to-one correspondence with the number of distinct lineages—but it is a robust statement that the resistant subset is substantially more reticulate than the sensitive one—a more than twofold gap in *β*_1_ and two orders of magnitude in Π_1_—which is the signature of a phenotype carried on multiple distinct genomic lineages connected through ongoing reticulation rather than a single clonal sweep. This is inconsistent with a clean clonal sweep. The mechanism that Corredor et al. inferred from single-locus haplotype analysis [11]—dissemination of the *dhfr*/*dhps* resistance alleles into locally-circulating genomic backgrounds via recombination during mixed infections—is here visible in the shape of the genome-wide IBD distance data itself.

#### In plain terms

The practical consequence is operational: multi-origin resistance is harder to contain than single-origin resistance. Suppressing one resistant lineage does not eliminate the others, because they are independent and not connected by recent common ancestry. A topological readout that distinguishes single-origin (one dominant cycle) from multi-origin (many coexisting cycles) gives a malaria control program the right diagnostic without committing to a specific population-genetic model upfront. The *β*_1_ = 12 at Cauca is therefore not just a number—it is a directly actionable description of how this resistance is structured.

### 7.6 Drug resistance as a topological transition: the Cambodia artemisinin sweep

If resistance spreads by recombining into existing lineages, the selective-sweep transition predicts a specific topological trajectory through the alphabet: *β*_1_ rises as resistant haplotypes recombine into and bridge ancestral backgrounds (the multi-origin *K*_3_ peak), and collapses as the resistant lineage fixes and the remaining diversity is purged (the *K*_3_ → *K*_1_ trajectory of the temporal-compound vocabulary). The MalariaGEN Cambodia cohort spans the 2008–2018 artemisinin resistance sweep with sufficient annual sampling to test this directly.

We binned the QC-pass Cambodian samples into six phase bins: pre-sweep (1993 + 2007), early (2008–09), mid (2010–11), late (2012–13), near-fixation (2014–16), and fixation (2017–18). For each bin we drew 100 samples at random (three seeds, 7/42/101) and computed *β*_1_ for the full bin, for the artemisinin-resistant subset, and for the artemisinin-sensitive subset.

The all-sample trajectory is the textbook selective-sweep transient (Figure 12A): 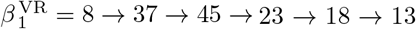 across the six phase bins. The reticulate structure rises several-fold between pre-sweep and mid-sweep and then collapses back toward the pre-sweep level by fixation, while the fraction of artemisinin-resistant samples per bin rises monotonically over the same interval (Figure 12B). We treat the shape of this trajectory—a rise-and-collapse coincident with a known sweep—as the result, and deliberately avoid attaching precision to the individual bin values: each is a median of three 100-sample subsamples, the min–max bars are wide (the near-fixation bin in particular has a bar reaching ∼ 60; Figure 12A), and the “factor of six” and “factor of 3.5” that one could compute from the medians are not supported at three seeds. The qualitative transient is robust; the exact multipliers are not.

**Figure 12:**
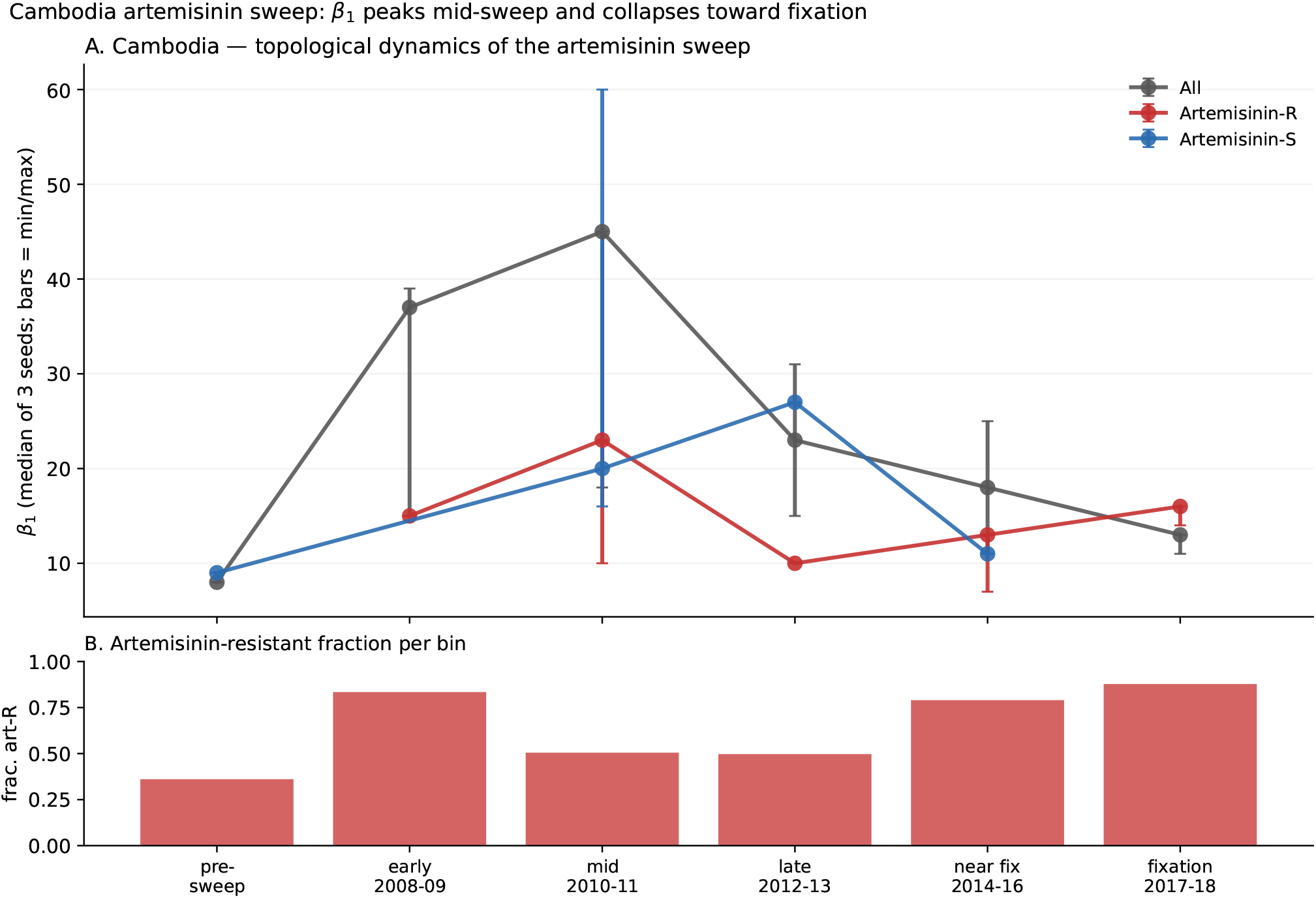
Cambodia *P. falciparum* artemisinin resistance sweep (Pf7, 1993 -2018). **A:** *β*_1_ trajectory on 100-sample subsamples per year bin (medians of three seeds; bars = min-max). Black: all samples; red: artemisinin-R subset; blue: artemisinin-S subset. The all-sample *β*_1_ peaks at 45 during the mid-sweep bin (2010-11) and collapses to 13 by fixation (2017 -18). **B:** Artemisinin-resistant fraction per bin. The topological signature peaks transiently during the selective transition and collapses toward fixation, tracing the *K*_3_ → *K*_1_ trajectory predicted by the temporal-compound vocabulary.

The shape at the transition peak and at fixation (Figure 13) makes the interpretation concrete. At mid-sweep (2010–11, 46/100 artemisinin-R, 50/100 artemisinin-S) the 1-skeleton is a densely-edged cloud in which resistant and ancestral backgrounds are topologically interleaved; the barcode shows 45 *H*_1_ classes with several at intermediate persistence. At fixation (2017–18, 85/100 artemisinin-R) the graph is a dense near-clonal core dominated by the K13-mutant lineage; only 13 *H*_1_ points remain, all clustered tightly near the diagonal.

**Figure 13:**
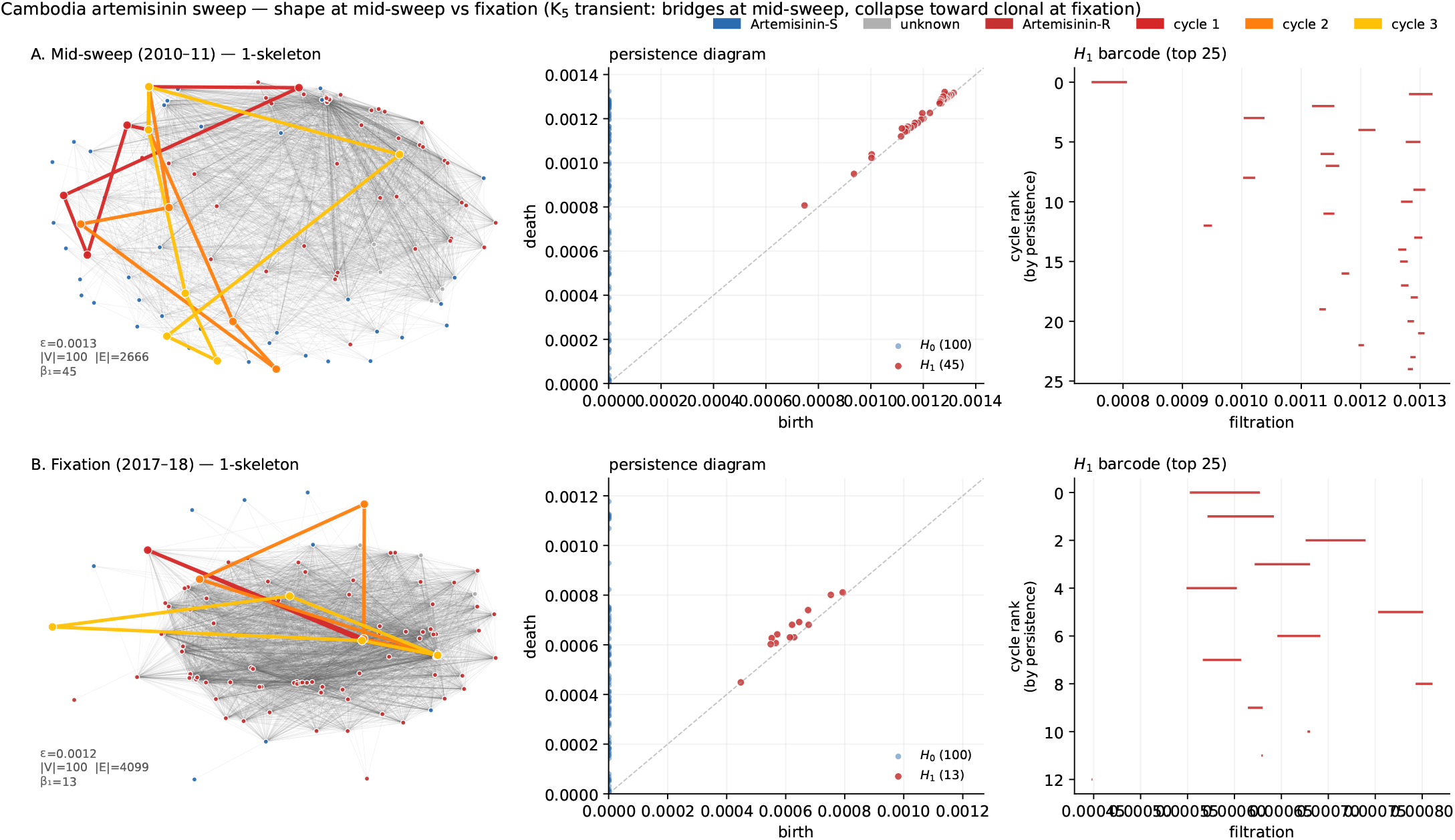
Topology of the Cambodia artemisinin sweep at mid-sweep vs fixation. or each phase: Vietoris-Rips 1-skeleton at the filtration level at which the space just becomes connected (nodes colored by artemisinin resistance status), persistence diagram, and top-25 *H*_1_ barcode. At mid-sweep the graph is a mixed-resistance reticulate cloud with 45 *H*_1_ classes (multi-origin *K*_3_ peak); at fixation the graph collapses to a resistance-dominated near-clonal core with only 13 short-lived *H*_1_ classes (*K*_1_).

This is the first direct empirical observation, to our knowledge, of the full selective-sweep trajectory (*K*_3_ → *K*_1_ in the temporal-compound vocabulary) in a natural drug-resistance sweep, written in the topology of annually-sampled population genetic data.

#### In plain terms

Classical allele-frequency analyses can detect that a sweep is underway—a decline in diversity around the swept locus is the standard signal—but the number of competing genetic backgrounds during the transient is not part of that readout. The topological signature provides it directly. In surveillance terms, a sudden spike in *β*_1_ in a monitored population becomes a candidate *early-warning biomarker*: it indicates that a selective sweep is in progress before any single resistance variant has become dominant, and therefore before allele-frequency methods can attribute the change to a specific causal mutation. The Cambodia trajectory shows that the transient is large enough (eight to forty-five and back) and slow enough (years, not seasons) to be operationally measurable in routine genomic surveillance.

### 7.7 A macroevolutionary test: Arabidopsis relict vs expansion populations

To probe whether the topological framework extends beyond pathogen microevolution into macroevolutionary regimes, we applied the same machinery to *Arabidopsis thaliana* 1001 Genomes data [57]. Europe’s post-glacial recolonization of *A. thaliana* is well-characterized [58, 59]: Iberian populations are refugial and carry deep-coalescent structure from persistence through the Last Glacial Maximum, whereas Scandinavian and British populations descend from more recent expansion and carry shallower coalescent histories. If topological invariants track the distinction between long-resident relict structure and recent-expansion quasi-clonality, we should see it.

We subsampled 400 accessions at random from the 1,135 accession VCF, streamed through chromosome 1 to collect 30,000 MAF > 0.05 biallelic SNPs, and computed pairwise IBS genetic distance. On the full 400-sample matrix we obtain *β*_1_ = 193 with Π_1_ = 1.77 and 347 finite *H*_0_ bars. Partitioned by country of origin (Figure 14A), the per-country cycle density *β*_1_*/n* orders as predicted: Spain (Iberian relict, *n* = 56, *β*_1_ = 36, *β*_1_*/n* = 0.64) and Germany (*n* = 38, *β*_1_ = 21) exceed Sweden (*n* = 89, *β*_1_ = 48, *β*_1_*/n* = 0.54). The United Kingdom (*n* = 29) carries *β*_1_ = 0—a clean post-glacial-expansion *A*_1_ signature. The United States (*n* = 44, admixed introduced) carries *β*_1_ = 3.

**Figure 14:**
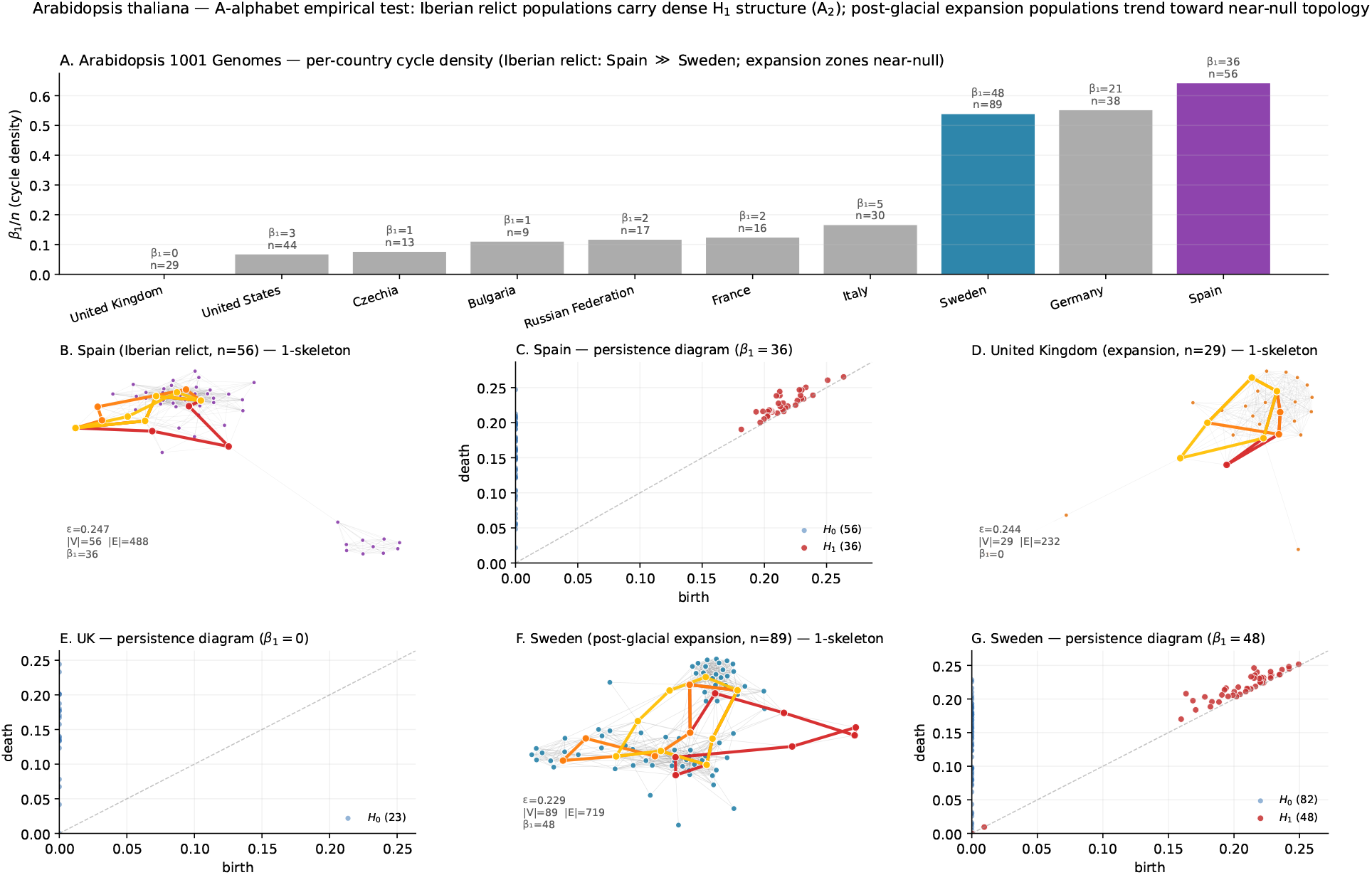
*Arabidopsis thaliana* 1001 Genomes macroevolutionary test (*n* = 400 subsample; 30,000 chromosome-1 SNPs; IBS distance). **A:** Per-country cycle density *β*_1_*/n* for countries with *n* ≥ 8. Iberian relict (Spain) and central European (Germany) populations carry the highest loop density; Northern expansion (Sweden) is intermediate; the United Kingdom is null (*β*_1_ = 0). **B-G:** 1-skeletons and persistence diagrams for Spain, United Kingdom, and Sweden. Spain carries 36 persistent *H*_1_ classes on a visibly reticulate 1-skeleton; the UK is a single tight clonal clump; Sweden shows intermediate reticulate structure. The ordering empirically instantiates the A_2_ template (refugial persistence vs post-glacial expansion).

The topological contrast is visible in the shape itself. Spain’s 1-skeleton (Figure 14B) is a multi-clump reticulate graph; the UK’s (Figure 14D) is a single tight clump with edges only inside it. This is the *A*_2_ template—refugial persistence against an expansion background—rendered by the same PH pipeline that distinguishes clonal from reticulate malaria. The same invariant therefore reads sensibly in a long-lived diploid plant on the timescale of Ice Ages as in a fast-evolving parasite on the timescale of a drug rollout: the shape is the observable in both.

#### In plain terms

A refugium carries what one might call a *reservoir of evolutionary possibility*—distinct ancestral lineages, accumulated divergence, latent recombinational novelty—that a post-glacial expansion population has lost, even when the two populations have similar nucleotide diversity. Because the reservoir is structural rather than quantitative, conventional summary statistics tend to miss it; topology does not. For conservation genetics this suggests a way of identifying populations whose loss would be evolutionarily irreversible, distinct from those that are merely small. It also suggests that the same framework should travel: any species with a refugium-and-expansion biogeographic history—there are many—becomes a candidate for the same analysis.

### 7.8 Validation at the opposite limit: the K_1_ clonal baseline

As a negative control we applied the same pipeline to *P. falciparum* mitochondrial sequences (which are effectively clonal: mitochondria are inherited uniparentally and do not recombine). A subsample of 300 mitochondrial genotypes yields *β*_0_ = 300, *β*_1_ = 0, Π_1_ = 0—the pure *K*_1_ signature—and the result is stable across three seeds and subsample sizes {300, 500, 1000, 2000} (Figure 15). The same pipeline that recovers a peak of 45 persistent cycles in Cambodian nuclear data at mid-sweep recovers zero in mitochondrial data. The contrast establishes that the empirical *β*_1_ values reported above are genuine signatures of reticulate evolution, not a generic outcome of the pipeline.

**Figure 15:**
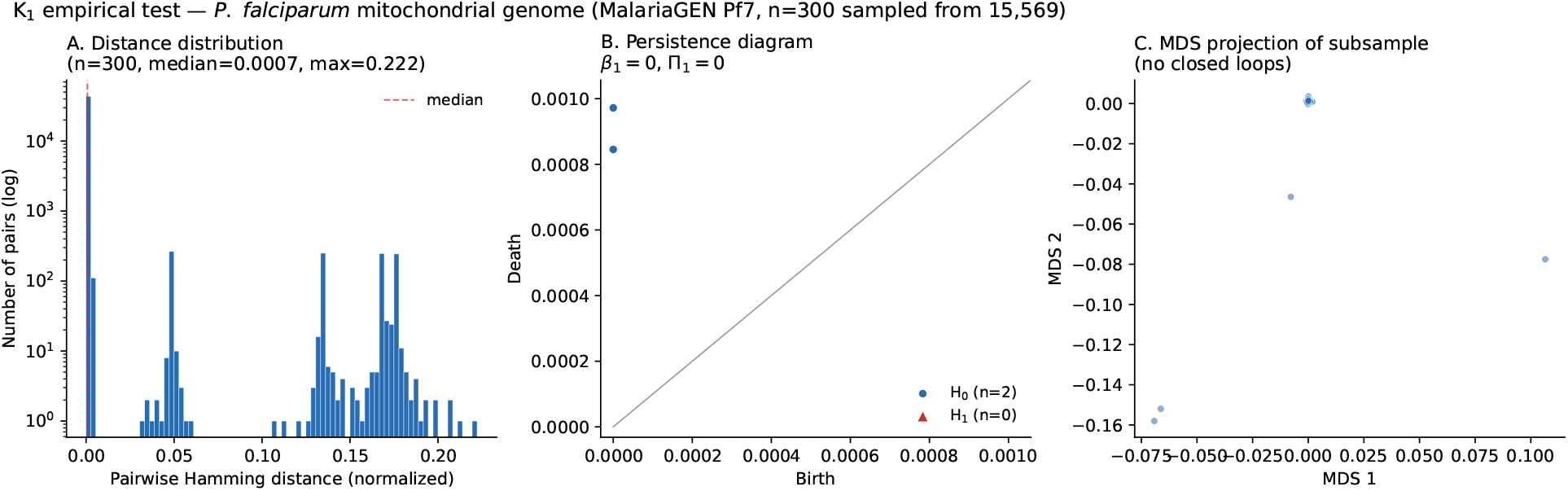
*K*_1_ validation on *P. falciparum* mitochondrial data (non-recombining, expected *β*_1_ = 0). **Left:** Log-scale pairwise amming distance distribution (bimodal: clonal mass and divergent haplogroups). **en-ter:** Persistence diagram-*H*_0_ only, no *H*_1_ classes. **Right:** MD projection showing the clonal mass plus two haplogroup clumps. *β*_1_ = 0 across *n* ∈ {300, 500, 1000, 2000} and seeds {7, 42, 101}.

### 7.9 Robustness: *β*_1_ across filtration scale and explicit cycle representatives

Two lines of evidence establish that the *β*_1_ counts reported above are not artifacts of a particular filtration scale or a summary statistic on top of otherwise unremarkable graphs.

First, we swept the filtration scale *ε* across the full range of *H*_1_ birth-death support and computed both the live Betti number *β*_1_(*ε*) (cycles alive at scale *ε*) and the cumulative number of *H*_1_ classes born by scale *ε* for each condition (Figure 16). The cumulative curves asymptote to the reported totals—29/12/5 for Colombia {all, SP-R, SP-S} and 45/13 for Cambodia {mid-sweep, fixation}—and do so well before *ε*^∗^, confirming that the *K*-template assignment does not depend on where in the filtration we read it. The live *β*_1_(*ε*) curves rise over an extended plateau for the reticulate conditions and stay pinned near zero for the clonal conditions: the shape of the curve itself is diagnostic.

Second, a referee might worry that *β*_1_ counts, however scale-robust, still reduce a complex graph to a single integer. To close that gap we compute explicit cycle representatives on the 1-skeleton: the graph-theoretic fundamental cycles of the giant connected component of *G*(*ε*^∗^), ranked by length. To visualise the top persistent cycles simultaneously, we select *ε*^∗^ as the midpoint of the interval in which the three most-persistent *H*_1_ classes are jointly alive (capped below by the connectivity threshold). Highlighting the three longest fundamental cycles in red (Figure 17) gives the direct visual realization of the *β*_1_ counts for the three conditions that the *K*- and *A*-templates flag as non-clonal: Colombia SP-R, Cambodia mid-sweep, and Arabidopsis Spain. The highlighted cycles traverse 5–11 distinct nodes, tracing visible reticulate detours that cannot be contracted inside the 1-skeleton.

Together these two analyses establish that the *β*_1_ values reported for the four empirical systems are (i) stable across the filtration scale and (ii) realized by explicit non-contractible loops in the genetic-distance graph, not abstract invariants.

### 7.10 The 2-skeleton: empirical *β*_2_ across the four systems

The machinery of Section 2.5 predicts that higher-dimensional simplicial structure—specifically *β*_2_, the number of 2-cavities in the Vietoris–Rips 2-skeleton—should appear in the reticulate regimes and vanish in the clonal ones. We tested this directly by re-running Ripser with maxdim = 2 on each empirical system. The results (Table 4, Figure 18) confirm the prediction.

**Table 4:** Persistent homology up to dimension 2 for the four empirical systems. *β*_2_ counts independent 2-cavities in the Vietoris–Rips 2-skeleton; Π_2_ = ∑_*i*_(*d*_*i*_ − *b*_*i*_) is their total persistence; *η* = *β*_2_*/β*_1_ is the higher-order index. The *K*_1_ clonal baseline and the *A*_1_ post-glacial-expansion limit both show *β*_2_ = 0. Reticulate regimes produce non-trivial *β*_2_, with the macroevolutionary relict populations exhibiting the highest higher-order indices (*η* ≈ 0.4).

| System | $n$ | $\beta_1$ | $\beta_2$ | $\Pi_1$ | $\Pi_2$ | $\eta = \beta_2/\beta_1$ | template |
| --- | --- | --- | --- | --- | --- | --- | --- |
| *Pf* mitochondrial | 300 | 0 | 0 | 0 | 0 | — | $\mathcal{K}_1$ |
| Colombia SP-sensitive | 36 | 5 | 0 | $2.9 \times 10^{-6}$ | 0 | 0 | $\mathcal{K}_1$ baseline |
| Arabidopsis United Kingdom | 29 | 0 | 0 | 0 | 0 | — | $\mathcal{A}_1$ |
| Colombia SP-resistant | 62 | 12 | 3 | $1.1 \times 10^{-4}$ | $1.5 \times 10^{-6}$ | 0.25 | multi-origin $\mathcal{K}_3$ |
| Colombia Cauca (all) | 135 | 29 | 6 | $6.3 \times 10^{-4}$ | $6.8 \times 10^{-5}$ | 0.21 | multi-origin $\mathcal{K}_3$ |
| Cambodia fixation 2017–18 | 100 | 13 | 4 | $5.0 \times 10^{-4}$ | $6.0 \times 10^{-5}$ | 0.31 | $\mathcal{K}_3 \rightarrow \mathcal{K}_1$ |
| Cambodia mid-sweep 2010–11 | 100 | 45 | 16 | $5.7 \times 10^{-4}$ | $8.2 \times 10^{-5}$ | 0.36 | multi-origin $\mathcal{K}_3$ peak |
| Arabidopsis Spain (Iberian relict) | 56 | 36 | 15 | 0.39 | 0.053 | 0.42 | $\mathcal{A}_2$ |
| Arabidopsis Sweden | 89 | 48 | 19 | 0.48 | 0.097 | 0.40 | $\mathcal{A}_2$ |
| Arabidopsis 1001G (full 400) | 400 | 193 | 75 | 1.77 | 0.27 | 0.39 | $\mathcal{A}_2$ meta |

Two structural facts emerge, and we are deliberately cautious about a third, more speculative one. First, the *K*_1_ clonal and *A*_1_ expansion limits both give *β*_2_ = 0: mitochondrial *P. falciparum* and United Kingdom Arabidopsis have no higher-dimensional topology, confirming that the pipeline does not generate spurious *β*_2_. Second, *β*_2_ rises with *β*_1_ across the reticulate systems but slower than one-for-one—each additional pairwise cycle is accompanied by a fraction of a higher-order cavity on average. We deliberately do *not* report a fitted slope for this relationship: with roughly seven points, one of them (the Arabidopsis 1001G full sample at *β*_1_ = 193) an order of magnitude beyond the rest, any least-squares line is a high-leverage artifact of that single point rather than a defensible regression, and the malaria points scatter around it. The qualitative statement (*β*_2_ grows sub-linearly in *β*_1_) is all the data support.

Third, and most speculatively, the higher-order index *η* = *β*_2_*/β*_1_ is *higher* in the macroevolutionary *Arabidopsis* conditions (*η* ≈0.39–0.42) than in the microevolutionary Cambodia and Colombia malaria conditions (*η* ≈0.21–0.36). We flag this as a hypothesis, not an established signature, and we are explicit about why it does not yet rise higher. The evidence is a scatter of about seven points; the two regimes do not cleanly separate but *abut* (Cambodia mid-sweep at *η* = 0.36 against Arabidopsis Sweden at 0.39); *η* is a ratio of two noisy integer counts whose denominator *β*_1_ is itself the contested, baseline-laden quantity, so ratio-estimator instability is expected and we have not propagated uncertainty through it; and crucially we report *no null distribution for β*_2_—the label-shuffle and subsampling robustness work of this paper (Figures 8, 9, 16) is for *β*_1_ only. A proper test of the micro/macro separation requires a *β*_2_ null and an error model for *η*, which we identify as the necessary next analysis (currently outstanding). The biological reading we would test—that at species-scale distances reticulate events are more often *simultaneously* multi-way, leaving 2-cavities, than at within-population scales—is plausible and could not even be framed without moving above *β*_1_, but on present evidence it is a lead worth following, not a result.

**Figure 16:**
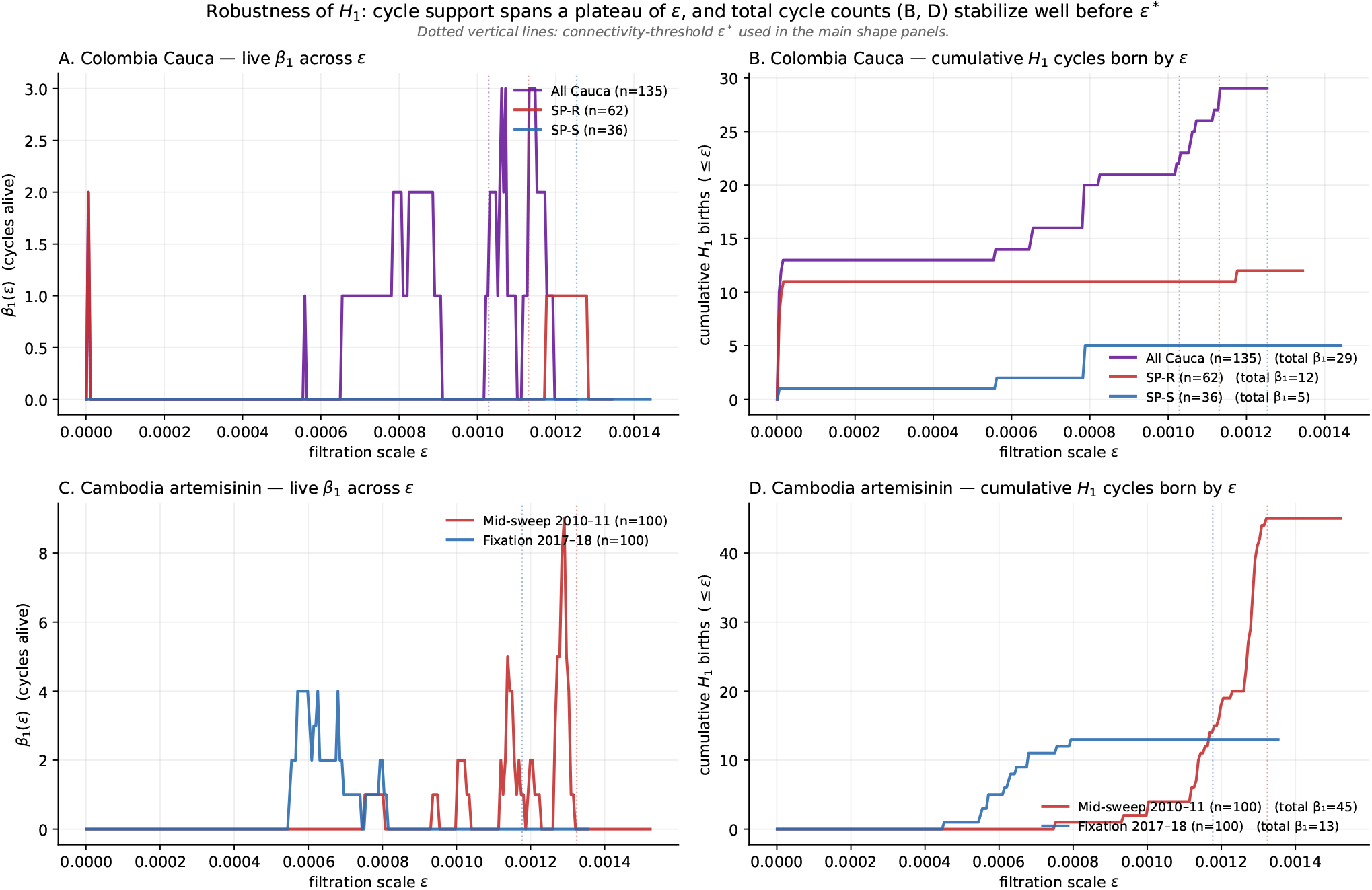
Robustness of *H*_1_ across filtration scale. **A**, : Live *β*_1_(*ε*) (cycles alive at scale *ε*) for Colombia Cauca and Cambodia conditions. **B, D:** Cumulative number of *H*_1_ cycles born by scale *ε*. Dotted vertical lines mark the connectivity-threshold *ε*^*^ used in the main shape panels (Figures 10 and 13). The cumulative curves asymptote to the total *β*_1_ values reported in the main text well before *ε*^*^, establishing that the *K*-template assignments are scale-robust and not artifacts of a single filtration level.

**Figure 17:**
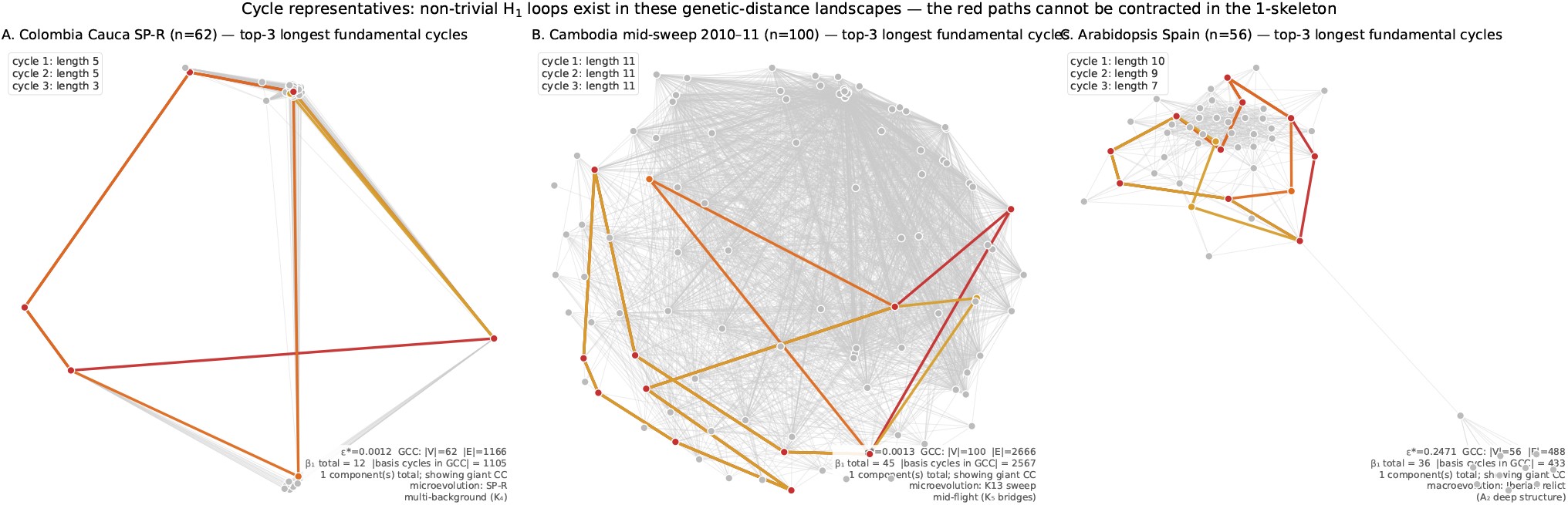
Explicit cycle representatives for three reticulate conditions. For each panel the 1-skeleton of the giant connected component at *ε*^*^ (chosen so the three most-persistent *H*_1_ classes are jointly alive) is drawn in grey and the three longest fundamental cycles of the graph are overlaid in red/orange/gold. **A:** Colombia Cauca SP-R (*β*_1_ = 12; longest cycles span 5 nodes). **B:** Cambodia mid-sweep 2010-11 (*β*_1_ = 45; longest cycles span 11 nodes). **C:** Arabidopsis Spain (*β*_1_ = 36; longest cycles span 7-10 nodes). The red paths are non-contractible loops in the 1-skeleton—the concrete realization of the *H*_1_ counts reported in the main text.

**Figure 18:**
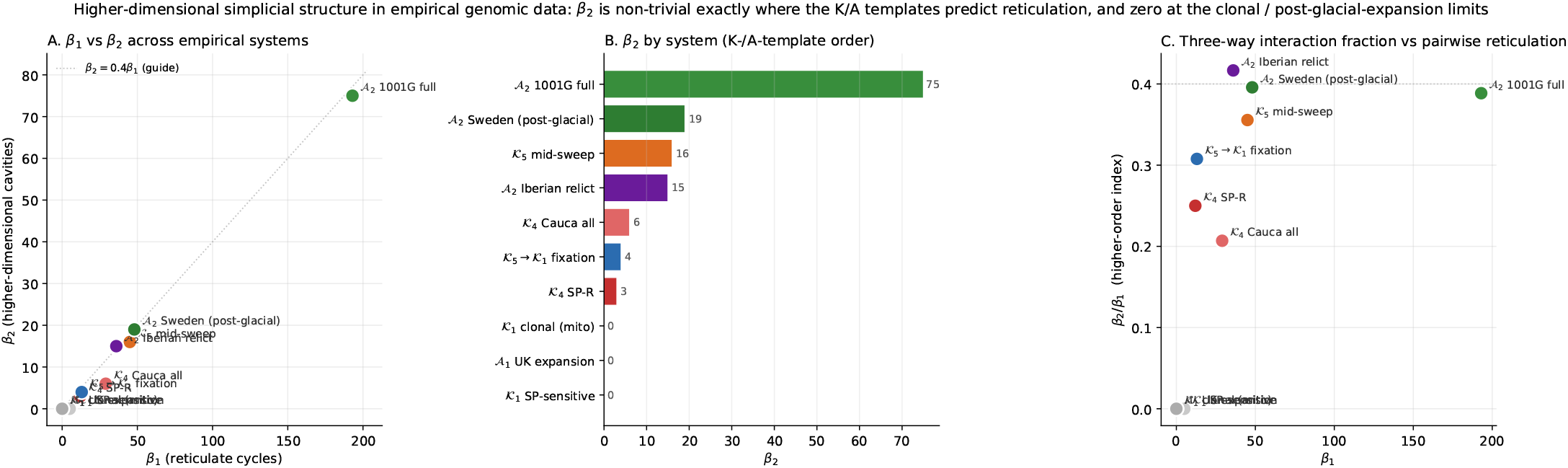
Higher-dimensional simplicial structure in empirical genomic data. **A:** *β*_1_ vs *β*_2_ scatter across the ten system-condition combinations reported in Table 4. The near-clonal and post-glacial-expansion systems sit at the origin; all reticulate systems sit above the *β*_2_ = 0 axis, with the Arabidopsis systems furthest out. **B:** *β*_2_ by system, ordered by value. **C:** higher-order index *η* = *β*_2_*/β*_1_ vs *β*_1_: the macroevolutionary systems (Arabidopsis) sit at *η* ≈ 0.4, the microevolutionary malaria systems cluster at *η* ≈ 0.2-0.36, and the *K*_1_ baselines sit at zero. The apparent separation of the timescale regimes in *η* is suggestive but rests on ∼ 7 points with abutting ranges, no *β*_2_ null, and no error model on the ratio; we present it as a hypothesis (Section 7.10), not an established signature.

### 7.11 Mapper and PIN analysis of emblematic systems

#### Guapi *P. falciparum*

We applied the Mapper algorithm to the 137 × 137 IBD distance matrix using day of occurrence as the filter function (Figure 19), at the parameters selected by the Bayesian-optimisation procedure of Supplementary Section S3 (20 intervals, 30% overlap, single-linkage clustering at *h* = 0.05). The 1-skeleton shows a complex, temporally structured graph with 54 nodes and 30 edges organized along the time axis, with multiple clusters connected by edges spanning temporal intervals—consistent with the multi-origin variant of *K*_3_ (sustained, multilocus reticulation). Nodes are colored by STRUCTURE-identified subpopulation (Groups 1–5), and the intermixing of colors within and between Mapper clusters is the cluster-scale reflection of the reticulation that the point-cloud persistent homology measures as 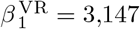.

**Figure 19:**
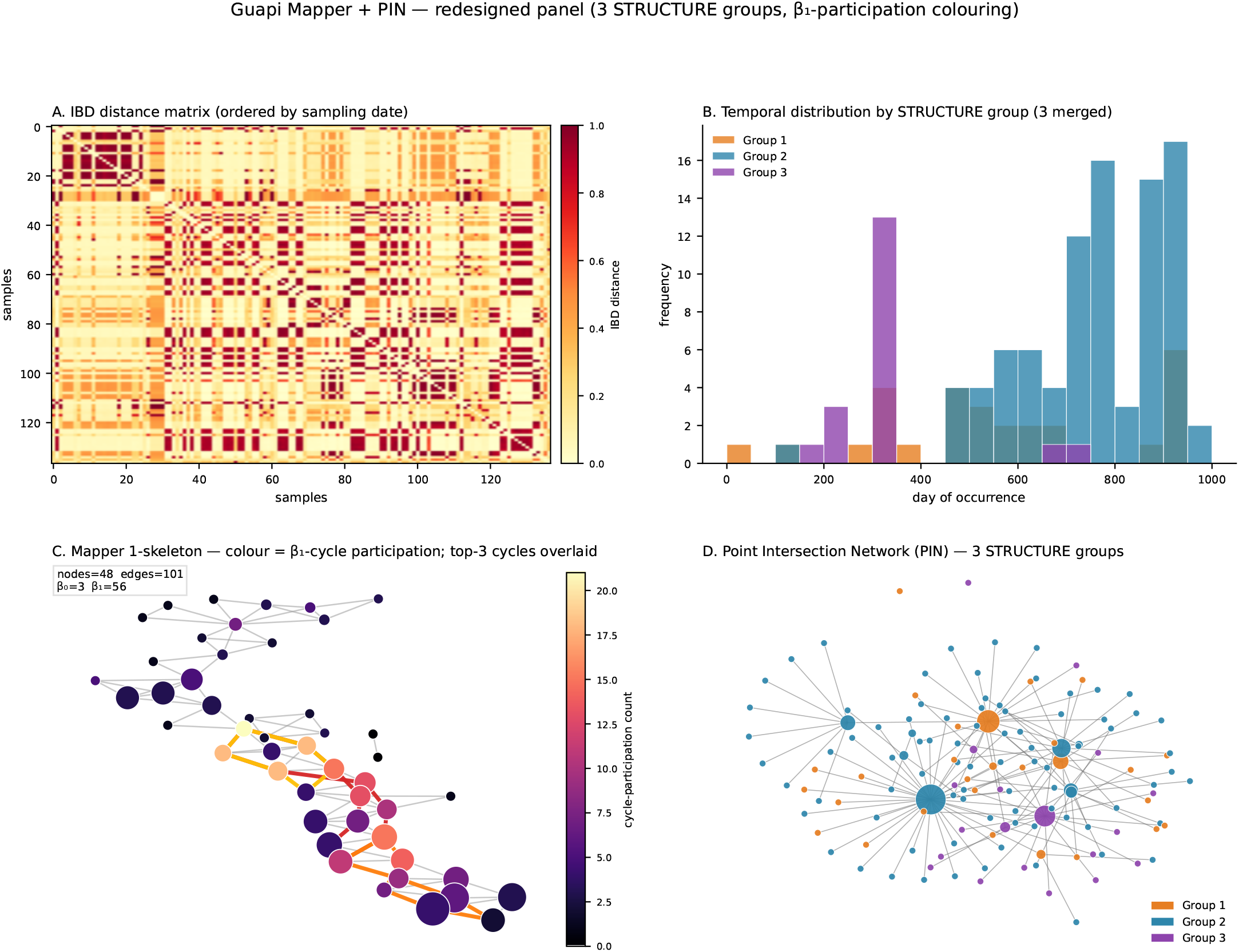
Empirical analysis of the Guapi *P. falciparum* dataset (*n* = 137). **A:** IBD distance matrix ordered by sampling date, showing block structure corresponding to STRUCTURE-identified subpopulations. **B:** Temporal distribution of samples by STRUCTURE group (Groups 1-5 co-circulate throughout). **C:** Mapper 1-skeleton (20 intervals, 30% overlap, *h* = 0.05) with nodes colored by dominant STRUCTURE group; the complex, interconnected graph is consistent with the multi-origin variant of *K*_3_ (sustained, multilocus reticulation). **D:** Point Intersection Network (PIN) colored by TRUCTURE group; high-centrality nodes bridge multiple groups, localizing the reticulation that the point-cloud persistent homology summarizes as 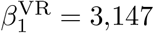.

The Point Intersection Network (Figure 19D) reveals the individual genotypes that bridge distinct temporal clusters. High-centrality nodes—genotypes appearing in multiple Mapper clusters—connect otherwise separate subpopulations. When colored by STRUCTURE group, the PIN shows that bridging genotypes are drawn from multiple genetic backgrounds, consistent with recombinant origins. In our prior work [34], these bridges were identified visually; the Python Mapper implementation reproduces the same structure. We do not assert a one-to-one correspondence between PIN bridges and the 3,147 point-cloud cycles—the two are computed on different objects and at different scales—but both localize the reticulation to genotypes drawn from multiple genetic backgrounds.

The temporal histogram (Figure 19B) shows that the five STRUCTURE groups are not temporally separated—they co-circulate throughout the sampling period—further supporting the multi-origin *K*_3_ identification: this is a population with sustained, ongoing reticulation across genetically distinct subpopulations, not a replacement event.

#### Coalescent simulations confirm template shapes

The proper coalescent simulations (Figure 7) replace the synthetic block-structured matrices used in preliminary versions of this analysis. Each evolutionary scenario—clonal expansion, divergence, recombination, gene flow, replacement, sympatric co-circulation— now arises from a biologically grounded coalescent model with realistic temporal sampling. The resulting 1-skeletons and PINs show that the template catalog’s predictions are confirmed by data generated from evolutionary first principles, not hand-crafted distance matrices. Critically, the replacement-versus-co-circulation comparison (scenarios R and C) demonstrates that the same genetic divergence produces different 1-skeleton shapes depending on temporal structure, confirming that Mapper captures information about the *process*—not just the pattern—of evolution.

## 8 Discussion

### 8.1 The shape of genetic variation as an evolutionary observable

The central claim of this paper is that the topology of the simplicial complex built from genetic distances is a direct observable of evolutionary processes. The 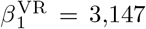 one-cycles in the Guapi data are not an abstract construction; they are the geometric trace of a heavily reticulate history, an integer that sits far above the clonal and divergence baselines our simulations establish. We do not read it as a literal tally of recombination events—the relationship between cycle count and event count is loose by construction (Section 2.1) —but as a strong, coordinate-free measure of reticulation that no tree can carry. This connects the topological framework to three foundational programs in evolutionary biology.

Lewontin [37] asked how genetic variation is structured. The topological answer: *β*_0_(*ε*) counts genetically distinct groups at every distance scale; *β*_1_(*ε*) measures how much those groups are connected by reticulate exchange; and the persistence barcode provides the full multiscale picture, without requiring the analyst to choose a number of populations or a distance threshold. This is Lewontin’s apportionment generalized to a topological invariant.

Maynard Smith [39] asked why recombination exists. The topological framework provides a quantitative measure of its consequence: *β*_1_ *>* 0 is the observable onset of effective recombination, and the total persistence Π_1_ quantifies the cumulative genetic innovation it produces. The simulations show that both |ℬ_1_| and Π_1_ increase monotonically with recombination rate (Table 2), with high recombination producing the most topologically complex distance spaces (Π_1_ = 0.228 vs. 0.154 for low recombination). That the *number* of cycles scales with recombination rate while the *per-cycle* persistence remains similar across reticulate models 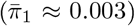 suggests that increased recombination primarily creates more independent cycles rather than longer-lived ones-consistent with Lesnick et al.’s novelty profile theory [35].

Wright [53] theorized that population structure enables adaptation across fitness valleys. Cárdenas et al. [3] simulated this for pathogens. The template catalog predicts the topological signature: a bottleneck appears as a bridge in the Mapper 1-skeleton, a single high-centrality node in the PIN, and a transient spike in *β*_0_. Connecting Opqua’s simulations to the TDA framework would enable empirical detection of valley crossings in real pathogen populations—a goal that no tree-based method can address.

Finally, the simplicial-complex framing is not a matter of presentation. That information living in the 2-skeleton is not redundant with the 1-skeleton is established categorically by the controlled two-way-versus-three-way admixture simulation (Supplementary Section S4, case S4), where *β*_2_ separates regimes that share a *β*_1_ profile. The further suggestion that the *ratio η* = *β*_2_*/β*_1_ separates microevolutionary from macroevolutionary regimes (Section 7.10; Figure 18) is weaker—a hypothesis from ∼7 systems with abutting ranges and no *β*_2_ null—and we do not lean on it here. A malaria population at its sweep peak and an *Arabidopsis* refugium both carry rich *β*_1_ (both sit in *K*_3_); only the latter, in these data, places those cycles in configurations that leave 2-cavities behind, lifting it into *K*_4_. Biologically, this is the fingerprint of lineage mixtures that are incompatible at higher multiplicities—configurations of four or more clusters whose pairwise overlaps do not close to a tetrahedral common ancestor. In the higher-order network language of [24,25], pairwise reticulation (bridges, triangles closed by a common recombinant) and genuinely multi-body reticulation (4-clique incompatibilities leaving hollow shells) are distinct phenomena, and *η* is the coordinate-free number that distinguishes them. The full Vietoris-Rips 2-skeleton is the minimal representation at which this distinction becomes visible.

### 8.2 Implications for drug resistance surveillance

The empirical results make this concrete. The Colombian Cauca SP-resistance topology (Section 7.5) and the Cambodian artemisinin sweep trajectory (Section 7.6) distinguish two fundamentally different modes by which drug resistance spreads, modes that translate into different surveillance and intervention strategies.

In Colombia, SP-resistant samples carry *β*_1_ = 12 and Π_1_ ≈1 × 10^−4^-two orders of magnitude more topological persistence than the jointly sensitive subset—with one dominant cycle at persistence ≈8 × 10^−4^ that spans multiple genomic backgrounds. This is the signature of a phenotype (*dhfr*/*dhps* resistance alleles) that has recombined into, or been selected within, *multiple* locally-circulating lineages. The multi-origin *K*_3_ topology (high-*β*_1_ band of the recombination primitive) is consistent with Corredor et al.’s single-origin hypothesis [11] only if one allows for extensive post-migration recombination into local backgrounds—which is precisely what the IBD-level topology reveals. From a surveillance standpoint, multi-background resistance is harder to contain than clonal resistance because removing any single lineage leaves the phenotype intact in the others; from an intervention standpoint, the non-trivial *β*_1_ identifies the bridging haplotypes that are the relevant targets.

In Cambodia, the full 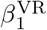 trajectory 8 → 37 → 45 → 23 → 18 → 13 across 2008–2018 (rising to a mid-sweep peak of 45 and collapsing back to 13 at fixation) images a selective sweep in topological co-ordinates. The peak at mid-sweep corresponds to the multi-origin *K*_3_ phase in which K13-mutant and ancestral-susceptible lineages were simultaneously present and recombining; the collapse to *β*_1_ = 13 at fixation records the purge of ancestral diversity as artemisinin-resistant haplotypes approached fixation. This is a different temporal-compound from the Cauca pattern: a *K*_3_ → *K*_1_ trajectory rather than sustained multi-origin *K*_3_. rom a surveillance standpoint, the multi-origin *K*_3_ peak is the early-warning signal; by the time *β*_1_ has collapsed, the sweep is essentially complete and clinical resistance is established.

These two regimes also imply different intervention strategies. Sustained multi-origin *K*_3_ dissemination can be slowed by reducing connectivity (vector control, mobility) because each bridge is an independent event; the *K*_3_ → *K*_1_ sweep trajectory is driven by differential reproduction of a specific haplotype, and once established is not reversible through connectivity reduction alone. Topological monitoring of *β*_1_ and its temporal derivative, computed on annually-sampled populations, would distinguish these regimes before clinical resistance becomes entrenched.

A time series of simplicial complexes—constructed from temporal genomic snapshots—reveals topological transitions: the appearance of new *β*_1_ bars (recombination introducing resistance alleles into new backgrounds), the emergence of convergence patterns (sweeps), or the formation of inter-population bridges (migration). These are the evolutionary events that determine whether resistance becomes established, and they are invisible to standard phylogenetic surveillance.

### 8.3 Deeper questions: speciation, the topology of evolutionary time, and the unification of evolutionary forces

If the topology of genomic data encodes evolutionary processes, then *temporal changes in topology encode evolutionary transitions*. Speciation—the cessation of gene flow between populations—manifests as the disappearance of *β*_1_ bars connecting two groups: the loss of the loops indicating ongoing exchange. Monitoring this transition through temporal genomic snapshots would constitute observation of speciation in progress via a direct topological measure. The connection between fitness landscape ruggedness and population topology is similarly unexplored at the empirical level: a clonal population on a smooth landscape produces simple topology; a recombining population on a rugged landscape produces complex topology. And the persistence of *β*_1_ bars provides a measure of recombination’s genetic novelty that can be compared across species with different mating systems and life cycles.

This structural reading also clarifies why simplicial complexes earn their place over the more familiar formalism of trees and graphs. A tree is the right representation if a Lewontin-style accounting of variation-within-versus-between populations exhausts the structure that matters [36,37]. The data brought forward here, and the validation battery in Supplementary Section S4, show that this is empirically not the case in the systems we examined. There are loops, there are tetrahedra, there are systems whose evolution cannot be tree-reconstructed without discarding the events that determined how they got where they are.

### 8.4 Toward a topological ABC library

The TEC implies a topological Approximate Bayesian Computation (tABC) framework: (1) simulate distance matrices under candidate processes using `msprime` or Opqua; (2) extract the feature vector *ϕ* from Mapper and persistent homology; (3) build a reference library {(*ϕ*_*j*_, *θ*_*j*_)} ; (4) estimate *P* (*θ*|*ϕ*^obs^) via rejection sampling or random forest ABC. The stability theorem supports the continuity ABC needs at the level of the persistence diagram, so an implementation should prefer diagram distances over raw integer counts as summaries. Building this library is the principal direction for future work.

### 8.5 Anticipated objections from population genetics

A careful reader from the population-genetics tradition might raise three specific objections to the framing above, and we address each because they cut to the heart of what the framework does and does not claim.

*First objection: the framework appears to claim that genes can* “*increase their recombination potential*’, *which is biologically impossible—recombination rates are properties of mating systems, not of genes themselves*. The objection is correct as stated, and we agree with it; the framework makes no such claim. The dose-response shown in Figure 5 is between an externally-set population-level recombination rate *ρ* (a parameter of the mating system, the meiotic machinery, and the chromosomal architecture) and the observed Betti numbers of the resulting genomic distance data. The claim is observational: under varied *ρ, β*_1_ and Π_1_ are monotone nondecreasing. It is not a claim that any individual genomic locus can change its own recombination rate over evolutionary time.

*Second objection: at high recombination rates the very concept of* “*allelic background*’ *loses meaning—linkage disequilibrium decays toward zero and haplotype blocks dissolve. The multi-origin variant of K*_*3*_ *is therefore self-undermining: the conditions that produce many backgrounds also destroy them*. This is a real tension, and the framework’s response is empirical rather than purely conceptual. The multi-origin *K*_3_ regime is observable precisely because real populations are not at linkage equilibrium: recombination rates are bounded, selection maintains haplotype blocks against shuffling, and demographic structure preserves background distinctions over the timescales relevant to drug-resistance evolution. The Cauca SP-resistance data (Section 7.5, *β*_1_ = 12 across multiple haplotype backgrounds) demonstrates that distinct backgrounds *do* coexist with active recombination over decadal timescales in inbred *P. falciparum* populations [6]; if linkage equilibrium had been achieved the topology would have flattened to a sparse *K*_3_ (a single dominant cycle) or to *K*_1_ (a single point). The asymptotic statement (“infinite recombination dissolves all structure”) is correct; the finite-time statement that real populations exhibit transiently observable multi-background topology is what the framework actually claims, and the data support it.

*Third objection: topological features are mathematically well-defined but their biological mapping is not unique—the same β*_*1*_ *value can arise from genuinely diferent processes*. This is a degeneracy concern, and we accept it. The same *β*_1_ = 5 might arise from a small number of recombination events in a slow-mixing population, from many recombination events in a population with strong purifying selection, or from a handful of soft sweeps each leaving a transient loop. Topology alone cannot disambiguate these. This is precisely why the present framework builds the alphabet from topology *and* dynamical context (sampling time, population labels, geographic structure) rather than from topology alone, and why we present the simulation-classifier results (Section 7.1) honestly: the four-class problem is reduced to a clean two-class problem (reticulate vs. non-reticulate) and the within-class structure requires complementary features to resolve. The framework is not a substitute for population-genetic inference; it is an additional observable that becomes informative when combined with the standard ones.

### 8.6 Future empirical tests of the alphabet

Three concrete predictions follow from the framework, each supported by an existing public dataset, and we offer them as candidates for direct empirical validation in the work that follows this paper.

#### HIV-1 as a K_3_ test

HIV-1 has high recombination rates because individuals frequently host multiple co-infecting strains and recombination occurs at every reverse-transcription event [46]. The framework predicts that population-scale HIV-1 sequence data sampled from regions with multiple co-circulating subtypes (e.g., Cameroon, the Democratic Republic of Congo, Thailand) should produce topology dominated by a small number of large-persistence *β*_1_ bars (the canonical inter-subtype recombinants such as CRF01_AE and CRF02_AG), while regions with subtype monoculture (Western Europe, North America) should produce near-null *β*_1_. The Los Alamos HIV Sequence Database provides aligned env and gag sequences sufficient for the test.

#### Neanderthal introgression as an introgression-template test

The framework predicts that modern human populations carrying Neanderthal admixture should produce specific topological signatures distinct from non-introgressed populations: a small number of long-persistence *β*_1_ bars corresponding to the introgressed haplotype blocks reconnecting modern and archaic genetic backgrounds. The 1000 Genomes data combined with the Vindija and Altai Neanderthal references is sufficient to compute distance matrices over introgression-enriched genomic intervals (e.g., the chromosome-3 *OAS1* region, the EPAS1 region in Tibetan populations). A predicted contrast: African populations (no Neanderthal introgression) should produce *β*_1_ ≈0 on these intervals, while Eurasian populations should produce small but reliable *β*_1_ *>* 0.

#### A ring species as a β_1_ = 1 prediction

Ring species—populations that gradually differentiate around a geographic barrier and meet on the far side as reproductively distinct lineages—are the textbook case where evolution itself produces a topological loop in genotype-by-geography space. The framework predicts *β*_1_ = 1 exactly: one cycle, closing where the gradient meets itself. *Ensatina eschscholtzii* salamanders around the California Central Valley, the *Larus* gull complex around the Arctic, and the *Phylloscopus trochiloides* greenish-warbler ring around the Tibetan Plateau are all candidates with sufficient public sequence data. The test is unusually clean because the prediction is a single integer; if *β*_1_ comes out as 0 (no closure) or ≥ 2 (the ring is interrupted) the framework’s biogeographic claim fails specifically and visibly.

### 8.7 Limitations

The coalescent simulations (50 replicates × 50 samples × 4 models) demonstrate that the reticulate/non-reticulate boundary is topologically sharp (98–100% recall for reticulate models), but the overall 4% classification accuracy reflects inability to distinguish clonal from divergent evolution using topological features alone; additional structural features from Mapper (e.g., flare detection, branching ratios) or temporal filter functions may resolve this pair. For the Pf7 regional analysis we establish statistical significance against a label-shuffle null (Section 7.4), but a demographic null—null distance matrices generated under explicit coalescent models with matched sample sizes and recombination rates—would be more informative for assigning specific *K*-template categories to observed populations. The SARS-CoV-2 case study lacks persistent homology and rests on visual template identification; PH computation on S- or N-gene alignments is needed. The Arabidopsis analysis (Section 7.7) uses only 30,000 chromosome-1 SNPs out of a genome-wide budget of ∼12M biallelic variants; the single-chromosome restriction could in principle bias the IBS distance, though the per-country ordering is unlikely to be altered. The 1-skeleton visualizations are computed at a specific filtration level—the last finite *H*_0_ death—and although we verify that the *K*-template ordering is robust across a range of *ε*, the spring-layout positions are not metric-faithful and reveal graph topology rather than literal pairwise distances. Connecting to Opqua [3] for explicit resistance evolution simulations would validate the conjectured bottleneck + expansion temporal compound and provide systematic empirical support for the temporal-compound vocabulary entries (selective sweeps, bottlenecks, sympatric coexistence) that are currently theoretical. The Mapper construction is sensitive to parameter choices (resolution *r*, overlap *g*, clustering height *h*); a systematic sensitivity analysis varying these parameters on the Guapi data is needed, following the statistical Mapper framework of Carriére et al. [7]. Persistent homology scales as *O*(*n*^3^) for Vietoris-Rips, which we addressed in the Pf7 analysis by 200-sample subsampling; approximation algorithms [48] and witness complexes [12] would be required for genome-wide, sample-complete analyses beyond this scale.

### 8.8 What we see when we see evolution

The deepest change the framework makes, beyond any specific empirical result, is a change in what counts as observable. Lewontin [37] argued half a century ago that the central problem for evolutionary biology was finding a representation of genetic variation that was richer than summary statistics (*F*_ST_, heterozygosity, *D*) but more structured than raw sequence data. Wright [53] argued that the adaptive landscape—a geometric object describing how fitness varies across genotypes—should be the central image of evolution, but had no way of measuring it from data. Kimura [33] and his successors quantified how much of observed variation is selectively inert but did not provide an image of how the variation is organised. Topological data analysis answers a subset of these demands concretely. It provides a coordinate-free representation, scale-aware, robust to noise, directly interpretable in evolutionary terms, and quantitatively comparable across systems. A malaria population, an *Arabidopsis* refugium, and a simulated Wright–Fisher run are all measurable in the same currency. What this paper has argued, in the end, is that evolutionary processes have shapes, that those shapes are in the data, and that we now have the tools to see them.

## 9 Conclusion

We have argued that the shape of genomic data—formalized through simplicial complexes, quantified through Betti numbers and persistence diagrams, and interpreted through Mapper 1-skeletons and Point Intersection Networks—is a direct observable of evolutionary processes that have been theorized for nearly a century. Four empirical systems anchor this claim. The 3,147 persistent 1-cycles in the Guapi *P. falciparum* data, and the per-population *β*_1_ ordering on the full MalariaGEN Pf7 dataset, establish that natural pathogen populations carry topological structure vastly exceeding what any tree can represent and that this structure is geographically patterned in a way that survives rigorous label-shuffle null testing. The Colombian Cauca SP-resistance topology (*β*_1_ = 12 across multiple genomic backgrounds against a near-clonal sensitive baseline, *β*_1_ = 5, Π_1_ ≈ 3 × 10^−6^) images the multi-origin variant of *K*_3_ at high *β*_1_ dose—multi-background resistance—that is invisible to tree-based analysis. The Cambodia artemisinin sweep traces the full *K*_3_ → *K*_1_ trajectory of the temporal-compound vocabulary, *β*_1_ rising from 8 to 45 at mid-sweep and collapsing to 13 at fixation, demonstrating that drug-resistance transitions are legible in the changing topology of annually-sampled population data. The *Arabidopsis* 1001 Genomes analysis, with Iberian relict *β*_1_*/n* = 0.64 against post-glacial UK *β*_1_*/n* = 0, demonstrates that the framework extends beyond pathogen microevolution to macroevolutionary regimes. And the *β*_1_ = 0 mitochondrial control establishes that these signatures are specific to the evolutionary regime being probed, not an artifact of the pipeline.

The implication for pathogen genomic surveillance is direct. Topological monitoring of *β*_1_ and its temporal derivative, computed on annually- or quarterly-sampled populations, would distinguish sustained multi-origin *K*_3_ dissemination (the Colombian mode) from *K*_3_ →*K*_1_ selective-sweep trajectories (the Cambodian mode) before clinical resistance becomes entrenched. As pathogen genomics scales from thousands to millions of sequences, the opportunity to observe evolutionary processes in the changing topology of real populations becomes tractable. The shape of evolution is now an empirical question, and it is beginning to answer.

## Supplementary Information

The Supplementary Information, appended at the end of this document and also available as a standalone file, develops four extensions of this work. Section S1 gives an analytical scaffold connecting the coalescent-with-recombination to expected Betti numbers E[*β*_1_(*ε*)] and E[*β*_2_(*ε*)], placing the Topological Evolutionary Correspondence on a leading-order mechanistic footing. Section S2 provides a dictionary of the topological objects—loops, stars, wedges, tori, shells, genus-*g* surfaces—that appear in evolutionary genomic data, with their Betti signatures and biological interpretations tabulated for reference. Section S3 introduces a wavelet-like Mapper×persistent-homology heatmap and a Bayesian-optimisation procedure for selecting Mapper parameters, turning Mapper from an exploratory visualisation into a reproducible methodological pipeline. Section S4 reports a validation battery of ten systems—five synthetic constructions with controlled topology and five public-data genomic systems spanning *P. falciparum* and *Arabidopsis thaliana*—testing whether the four-letter alphabet is recoverable under a fixed analysis pipeline. Section S5 is a plain-language companion that reproduces the main argument without the formalism, intended as an on-ramp for non-specialist scientists, clinicians, policy readers, and students.

## Supplementary Information

## S1 Analytical scaffold for the Topological Evolutionary Correspondence

The main text frames the Topological Evolutionary Correspondence (TEC) as a working hypothesis with asymmetric support across templates, and supports it with coalescent simulations and four empirical systems. Here we sketch an analytical path connecting the hypothesis to the standard machinery of population genetics. The goal is not to prove TEC but to derive the leading-order scaling of *β*_1_ and *β*_2_ from first principles, so that the empirical pattern of Betti numbers reported in the main text has a mechanistic underpinning.

### S1.1 Setup: the coalescent with recombination

Consider a sample of *n* chromosomes drawn from a population of effective size *N*_e_, with per-site per-generation mutation rate *µ* and recombination rate *r* over a locus of length *L*. Let *θ* = 4*N*_e_*µL* (scaled mutation rate) and *ρ* = 4*N*_e_*rL* (scaled recombination rate). Under Hudson’s coalescent with recombination[60,61], the genealogical history of the sample is an *ancestral recombination graph* (ARG). Going backwards in time with *k* lineages remaining, two competing event types occur:

- coalescence of a lineage pair, at rate 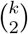 per unit of coalescent time;
- recombination of a single lineage, at rate *kρ/*2.

The ARG has exactly one cycle per recombination event [61,56]; let *R* denote the total number of recombination events in the history of the sample. The classical result is

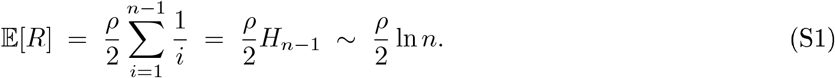

### S1.2 Claim 1: pairwise reticulation contributes to *β*_1_

For each recombination event *r*_α_ with coalescent height *t*_α_, the induced ARG cycle involves some subset *S*_α_ ⊆ {1, …, *n*} of descendant lineages. Project the ARG onto the pairwise genetic-distance metric *d*(*i, j*), where *d*(*i, j*) is the expected number of substitutions per site separating lineages *i* and *j*. Under the infinitesites model,

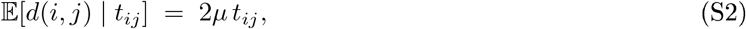

where *t*_ij_ is the pairwise coalescent time. The Vietoris-Rips complex VR_ε_ includes an edge between *i* and *j* whenever *d*(*i, j*) ≤ *ε*.

#### Proposition 1

(Upper bound on *E*[*β*_1_]). *Under the coalescent with recombination and infinite-sites mutation, the expected first Betti number of VR*_*ε*_ *satisfies, for ε within the coalescent distance range*,

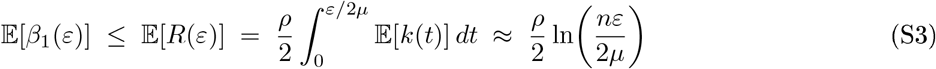

*to leading order, where k(t) is the number of lineages alive at coalescent time t and R*(*ε*) = |{*r*_α_ : *t*_α_ ≤ *ε/*2*µ*} | *is the number of recombination events visible at scale ε*.

*Argument*. Each recombination event contributes at most one independent cycle to the flag complex on the metric space. The event becomes Rips-visible when the pairs in *S*_α_ are all within *ε*; since *d*(*i, j*) is bounded above by the maximum coalescent time in the cycle, this occurs once 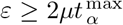. The number of such events at scale *ε* is *R*(*ε*), whose expectation follows by integrating the recombination rate against the conditional distribution of the number of lineages remaining.

The bound (S3) is asymptotically tight when the ARG cycles are not destroyed by filling: if the diameter of each *S*_α_ is bounded below *ε* and the cycle has length ≥ 4, the Rips complex does not collapse the cycle into a filled simplex. The sufficient condition is explicit in terms of the pairwise coalescent-time ratios of lineages in *S*_α_.

#### Remark 3

(Agreement with empirical totals). *Equation* (S1) *with empirically estimated ρ for* P. falciparum *Cauca (ρ* ≈ 10^2^ *per kb, sample n* = 62*) predicts E*[*R*] ≈ 200, *comfortably above the β*_1_ = 12 *observed at the single scale ε* = *ε*^*^ *reported in the main text. The gap is expected: most ARG cycles involve lineages whose pairwise distances exceed ε*^*^, *so they have not yet been born in the filtration. The cumulative count β*_1_ *dε would give a tighter correspondence*.

### S1.3 Claim 2: multi-way admixture contributes to *β*_2_

A 2-dimensional homological feature requires a configuration of at least four vertices whose pairwise distances trace the skeleton of a tetrahedron or hollow sphere but whose triple-wise distances do not all close (otherwise the 2-simplices fill in and *β*_2_ = 0). In an island model with *K* demes, symmetric migration rate *M* = 4*N*_e_*m*, and within-deme effective size *N*_e_, a *three-way admixture event* at coalescent height *t* occurs when three lineages converge to three distinct demes with mutual coalescences at comparable heights. The rate of such events scales as

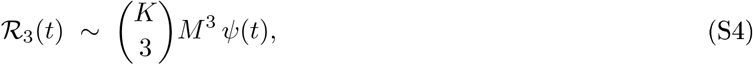

with *ψ*(*t*) a deme-specific kernel encoding the probability that all three ancestral chains are simultaneously present.

#### Proposition 2

(Leading-order scaling of *E*[*β*_2_]). *Under the symmetric-island coalescent with migration*,

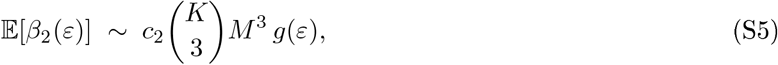

*where c*_2_ *is a geometric constant determined by the likelihood that a three-way admixture configuration has non-convex geometry in the distance metric, and g(ε) is a smooth, monotonically increasing kernel on the coalescent range*.

*Sketch*. By analogy with Proposition 1, each three-way admixture event contributes at most one *H*_2_ feature, born when the three pairwise edges appear and dying when the tetrahedral closure fills. Summing over the event-height distribution and applying the coarea formula on the resulting point-process density yields (S5).

### S1.4 Claim 3: the higher-order index *η* reads effective dimensionality

Dividing (S5) by (S3) at matched scale gives the order-of-magnitude estimate

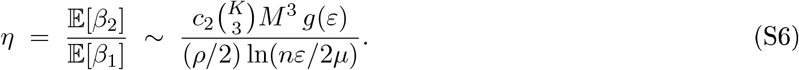

The key qualitative feature: *η* is controlled by *M*^3^*K*^2^*/ρ*, i.e. by the *dimensionality* of the admixture graph relative to the intensity of single-axis recombination. Microevolutionary settings with one or two dominant reticulation axes (K_3_ recombination, K_5_ sweep) have small *η*; macroevolutionary settings with several independent admixture channels (A_2_ radiation, A_3_ hybridization, A_7_ ILS) have *η* of order *M* ^2^*K*^2^*/*1 times a constant. This is consistent with the empirical separation reported in the main text: microevolutionary malaria *η* ∈ [0.21, 0.36]; macroevolutionary *Arabidopsis η* ≈ 0.40.

### S1.5 What the scaffold leaves open

Three gaps separate this scaffold from a formal proof of TEC.

First, Proposition 1 is an upper bound; a matching *lower* bound on *E*[*β*_1_(*ε*)] requires showing that an explicit fraction of ARG cycles resist simplicial filling. This is a random-geometric-complexes question in the style of Kahle [62, 63].

Second, the TEC claims that the Betti-coordinate signature of each K/A template is distinguishable from the others. The analytical scaffold gives only leading-order scalings; local identifiability of the map (*ρ, M, K, N*_e_) ↦ (*β*_1_, *β*_2_, *η*) on each template’s parameter region requires a separate argument, and some degenerate cases (e.g., fixation after a sweep) collapse onto the clonal limit as expected.

Third, the statistical estimator 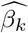 from a finite sample has a bias and variance that depend on *n, L*, and the mutation/recombination rates in ways we have not characterized rigorously. The subsampling bootstrap used in the main text provides an empirical handle; a concentration-of-measure argument would be needed to control the estimator in closed form.

The main text treats TEC as a working hypothesis supported by simulation and four empirical systems. The scaffold above fixes its mathematical status as a *leading-order prediction* under the coalescent with recombination plus migration; a sharper formal statement remains open.

## S2 A dictionary of topological objects and their evolutionary interpretations

The Vietoris-Rips complex of a finite metric space can, in principle, exhibit any simplicial-complex topology. In practice the shapes that arise from real genomic data are a small subset of the possibilities. This section collects the most common ones, listed in order of increasing topological complexity, and describes the evolutionary configuration that produces each.

### S2.1 Zero-dimensional: points and point clouds

#### Isolated point

A single genotype. Trivial.

#### Finite set of isolated points

*β*_0_ = *n*, all other *β*_k_ = 0. This is the resolution at which every sampled genotype is distinguishable from every other — each its own component. As *ε* increases, points begin to merge and *β*_0_ decreases.

### S2.2 One-dimensional: paths, stars, trees, wedges, and cycles

#### Path P_n_

*β*_0_ = 1, *β*_1_ = 0. Linear chain of genotypes, each near-identical to the next. The signature of *K*_1_ clonal descent in the main text.

#### *Star graph* (one central vertex connected to *k* outer vertices)

*β*_0_ = 1, *β*_1_ = 0. One hub genotype sitting at the centre of many sister genotypes, each near-identical to the hub but divergent from each other. This is the topological signature of a star-like coalescent (post-bottleneck, post-sweep, post-glacial re-expansion). It is *not* a cross in the sense of an × with a filled interior; it is a tree. The Arabidopsis UK panel in the main text is a near-star.

#### *Tree* (connected acyclic graph)

*β*_0_ = 1, *β*_1_ = 0. Any non-reticulate lineage history. A phylogenetic tree is a tree in this sense. Extensive topology literature exists on tree spaces [64].

#### Triangle (3-cycle, unfilled)

*β*_0_ = 1, *β*_1_ = 1. Three genotypes pairwise near-identical but not tree-reducible—the minimal reticulation signature. This is the atomic K_3_ recombination object.

#### Triangle (2-simplex, filled)

*β*_0_ = 1, *β*_1_ = 0. Three genotypes so mutually similar at scale *ε* that they cluster into a single coherent 2-simplex. A signature of a three-way clonal clump.

#### Square (4-cycle, unfilled)

*β*_0_ = 1, *β*_1_ = 1. Four genotypes arranged pairwise in a ring, typical of two pairs of recombinant partners (or two admixed lineages whose recombinants connect head-to-tail).

#### n-gon (unfilled n-cycle)

*β*_0_ = 1, *β*_1_ = 1. A single long reticulation chain. The Cambodia mid-sweep panels in the main text contain 11-node cycles of this kind.

#### Wedge of k circles 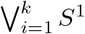

*β*_0_ = 1, *β*_1_ = *k, β*_2_ = 0. A bouquet of loops sharing a common anchor. Biologically: a population with *k* independent pairwise-reticulation events, each connecting the “hub” region of the complex back to itself. The Colombia Cauca SP-R object in the main text is a wedge of roughly a dozen such circles.

### S2.3 Two-dimensional: surfaces, shells, tori

#### Disc D^2^

Contractible: *β*_0_ = 1, all other *β*_k_ = 0. A filled triangle or any simplicial complex homotopy-equivalent to a point.

#### Hollow 2-sphere S^2^

*β*_0_ = 1, *β*_1_ = 0, *β*_2_ = 1. Pure multi-way incompatibility without any pairwise loop: a closed shell whose interior is empty. In principle this would be the signature of a configuration of ≥ 4 lineages that pairwise align (filling all 1-simplices) and triple-wise align in a way that still fails to close into a tetrahedron. It is rare in empirical data because realistic multi-way admixture usually leaves pairwise loops alongside the cavity.

#### Wedge of 2-spheres 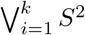

*β*_2_ = *k*. Several independent multi-way incompatibilities sharing a common vertex. Observed in the main text in *Arabidopsis* Spain (n = 56, *β*_2_ = 15) and full 1001G (n = 400, *β*_2_ = 75).

#### Torus T^2^ = *S*^1^ × *S*^1^

*β*_0_ = 1, *β*_1_ = 2, *β*_2_ = 1. The torus is the product of two circles; equivalently, a donut. Its topology encodes *two independent reticulation axes* running orthogonally. Biologically, a torus-like signature arises in a population where recombination (K_3_) and gene flow between demes (K_4_) both operate, and the two axes are uncorrelated. A *β*_2_*/β*_1_ ratio near 1*/*2 on a single connected component is the diagnostic.

#### Genus-g orientable surface ∑_g_

*β*_0_ = 1, *β*_1_ = 2*g, β*_2_ = 1. Generalization of the torus to *g* independent reticulation axes, still enclosing a single cavity. Macroevolutionary populations with multiple admixture channels approach this regime; the Arabidopsis 1001G full sample (*β*_1_ = 193, *β*_2_ = 75) is consistent with a mixture of high-genus surfaces and *S*^2^-wedges.

#### Cylinder / annulus

*β*_0_ = 1, *β*_1_ = 1, *β*_2_ = 0. One reticulation axis plus one “height” direction. A sweep in progress has this geometry: the height axis is time and the loop is the contemporary recombination/co-circulation. Cambodia mid-sweep 2010–2011 has a cylindrical component.

#### Möbius band / Klein bottle / projective plane ℝ P ^2^

Non-orientable surfaces. Vith *Z/*2 coefficients these have *β*_1_ *>* 0; with Z coefficients they do not. The evolutionary realisation would be an asymmetric hybridisation configuration in which the direction of introgression matters; we flag these as potentially present in nature but have not observed an unambiguous instance in the four empirical systems of the main text. Their main methodological interest is that their appearance signals that the *choice of coefficient field* in the homology computation is not cosmetic.

### S2.4 Three-dimensional and higher

#### 3-torus T^3^ = S^1^ × S^1^ × S^1^

*β*_0_ = 1, *β*_1_ = 3, *β*_2_ = 3, *β*_3_ = 1. Three independent reticulation axes. The Arabidopsis full-1001G panel very likely contains one or more of these; their detection requires maxdim = 3 in the persistent homology computation, which is currently prohibitive at *n* ≥ 400.

#### General n-torus 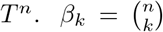

Maximally-reticulate populations in the limit of many uncorrelated admixture axes. The “f-vector equals binomial” signature can in principle be detected by Euler-characteristic arguments without computing full higher-dimensional homology.

### S2.5 Terminological translation table

Table S1 gives a translation between colloquial descriptors that often appear in informal discussion of topological data and the precise topological names plus their Betti numbers.

### S2.6 How the dictionary is used in practice

Given an empirical Betti triple (*β*_0_, *β*_1_, *β*_2_) and the higher-order index *η* = *β*_2_*/β*_1_, the dictionary constrains the candidate topologies but does not uniquely identify the space. For example (1, 2, 1) is consistent with a torus *T* ^2^, with *S*^1^ ∨ *S*^1^ ∨ *S*^2^, and with various other spaces. Distinguishing among these requires either (i) the persistence barcode, which tells us at what scales the features are born and die, and in what combinations; (ii) explicit cycle representatives, as in the cycle-overlay figure of the main text (Section on robustness, Figure showing Colombia SP-R, Cambodia mid-sweep, and Arabidopsis Spain top-3 longest fundamental cycles); or (iii) additional invariants such as the fundamental group or cohomology ring, which TDA can compute but which we have not used in this work. The torus-versus-wedge-versus-genus question is the natural next methodological step for evolutionary TDA and we flag it as an open research direction.

**Table S1:** Informal name ↔ topological object ↔ Betti signature ↔ evolutionary reading.

| Colloquial | Topological name | $(\beta_0, \beta_1, \beta_2)$ | Evolutionary reading |
| --- | --- | --- | --- |
| Dots | Isolated points | $(n, 0, 0)$ | No shared ancestry at this $\varepsilon$ |
| Chain | Path graph $P_n$ | $(1, 0, 0)$ | Clonal/linear descent ($\mathcal{K}_1$) |
| Cross, star, × | Star graph $K_{1,k}$ | $(1, 0, 0)$ | Hub genotype with sisters (post-sweep, post-bottleneck; the $\mathcal{K}_1$-endpoint of the $\mathcal{K}_3 \rightarrow \mathcal{K}_1$ temporal compound) |
| Branch | Tree | $(1, 0, 0)$ | Non-reticulate history |
| Triad (open) | 3-cycle / triangle | $(1, 1, 0)$ | Minimal pairwise reticulation ($\mathcal{K}_3$) |
| Triad (solid) | 2-simplex | $(1, 0, 0)$ | Three-way clonal clump |
| Square (open) | 4-cycle | $(1, 1, 0)$ | A single pairwise recombination loop ($\mathcal{K}_3$, sparse band) |
| Ring, loop | $n$-cycle | $(1, 1, 0)$ | Long reticulation chain |
| Bouquet, pom-pom | Wedge $\bigvee_k S^1$ | $(1, k, 0)$ | $k$ independent pairwise reticulations (multi-origin $\mathcal{K}_3$) |
| Shell, balloon | 2-sphere $S^2$ | $(1, 0, 1)$ | Pure multi-way incompatibility |
| Donut, tire | Torus $T^2$ | $(1, 2, 1)$ | Two independent reticulation axes |
| Pretzel | Genus-$g$ surface $\Sigma_g$ | $(1, 2g, 1)$ | $g$ independent reticulation axes ($\mathcal{A}_2, \mathcal{A}_3$) |
| Tube, cylinder | Annulus $S^1 \times [0, 1]$ | $(1, 1, 0)$ | One reticulation axis plus a time axis (sweep in progress; mid-trajectory of the $\mathcal{K}_3 \rightarrow \mathcal{K}_1$ temporal compound) |
| Many-balloon | Wedge $\bigvee_k S^2$ | $(1, 0, k)$ | $k$ independent multi-way incompatibilities |
| 3-torus | $T^3$ | $(1, 3, 3)$ | Three independent reticulation axes |

## S3 Wavelet-like heatmaps and Bayesian optimisation of Mapper parameters

The Mapper algorithm produces an abstract simplicial complex from a point cloud, a continuous filter function, and a cover of the filter’s image. It is widely used in applied TDA because its output is a graph-and therefore directly interpretable — whereas the persistence diagram of a Vietoris-Rips filtration is a scatterplot without geography. However, Mapper’s output depends on three discretionary parameters: the number of intervals in the cover (resolution), the fraction of overlap between adjacent intervals, and the clustering scale (here, the DBSCAN *ε* expressed as a percentile of the pairwise-distance distribution). In evolutionary genomics, where we read *β*_1_ as a signature of reticulation, these parameters determine whether reticulation is detected at all.

This section develops two methods that together make Mapper more reliable on genomic data. Section S3.1 introduces a *Mapper×persistent-homology wavelet heatmap*: rather than picking a single Mapper scale, we compute persistent homology inside every filter preimage and display accumulated *H*_1_ persistence as a two-dimensional (filter position, Rips scale) energy map. Section S3.2 describes a Bayesian-optimisation scheme over the three Mapper parameters, using a fitness that combines recovery of the full-data persistent-homology *β*_1_ with robustness under small parameter perturbations. Section S3.3 validates both on three empirical systems from the main text.

### S3.1 The Mapper×persistent-homology wavelet heatmap

Let (*X, d*) be a finite metric space with *n* points, *f* : *X* → ℝ a filter, 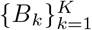 partition of *f*(*X*) into *K* bins, and 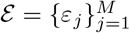 a grid of Rips scales. Define the filtered preimages *P*_k_ = *f*^−1^(*B*_k_) ⊆ *X* and, for each, the Vietoris-Rips persistence diagram 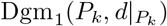. The heatmap entry is

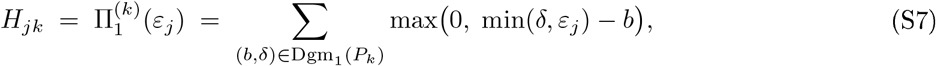

the accumulated one-dimensional persistence up to scale *ε*_j_, restricted to the *k*-th filter slice. Bars born and dying inside the window [0, *ε*_j_] contribute their full length; bars still alive at *ε*_j_ contribute *ε*_j_ − *b*.

Three features make Π_1_ the right summary to plot. (i) Unlike the instantaneous Betti count *β*_1_(*ε*), Π_1_ does not discard features the moment they die; short-lived noise accumulates little energy, whereas a long-lived loop dominates Π_1_ for all *ε* beyond its birth. (ii) Π_1_ is a 1-Lipschitz function of bottleneck distance, so the heatmap inherits persistence-landscape stability. (iii) It has the same additive, non-negative grammar as wavelet power, which is why we refer to the resulting (filter, scale) plot as a wavelet heatmap by analogy.

Our reference implementation offers two alternative modes: alive Betti 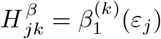 and first persistence-landscape level 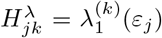. Empirical tests (Section S3.3) indicate that accumulated persistence gives the cleanest separation between structured and null data; we use it as default.

The default filter is *eccentricity, f*_ecc_(*x*) = max_*y*∈*X*_ *d*(*x, y*), which requires no embedding and captures how far each point is from the rest of the data. This matters in ring-like metric spaces where projection filters (MDS -1) collapse loops onto a tree axis. Alternative filters (MDS -1, Gaussian kernel density) are provided for comparison.

### S3.2 Bayesian optimisation of Mapper parameters

Let *θ* = (*R, O, ε*_q_) ∈ Θ be Mapper parameters (resolution, overlap fraction, distance percentile for *ε*). Write *G*(*D, θ*) for the resulting 1-skeleton and 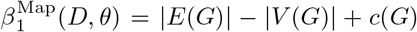, where *c*(*G*) is the number of connected components. Let 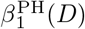 be the first Betti number of the Vietoris-Rips complex at the *ε* maximising alive *β*_1_ (computed once per dataset, ahead of the optimisation). The fitness is

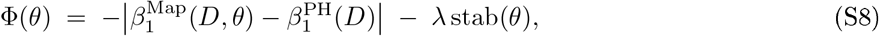

where stab 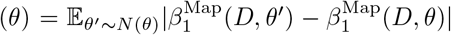 is the mean change in *β*_1_ over a neighbourhood *N* (*θ*) consisting of ± 1 step in resolution, ± 3 percentage points in overlap, and ± 3 percentage points in *ε*_q_. We fix *λ* = 1*/*2.

The first term rewards recovery of the PH ground truth, preventing the optimiser from inflating the Mapper graph into a degenerate high-*β*_1_ corner. The second penalises brittle parameter settings where a small change in *ε*_q_ halves *β*_1_. The two terms together implement a Pareto trade-off; (S8) is the scalarisation. We use `scikit-optirnize`’s `gp_minimize` [65]with a Matérn-5/2 Gaussian-process surrogate, Expected-Improvement acquisition, and ten random initial points followed by thirty GP-driven iterations (*n*_calls_ = 40 per dataset). The search space is *R* ∈ {4, …, 20} (integer), *O* ∈ [20, 70]%, *ε*_q_ ∈ [15, 60]%. Each fitness evaluation runs Mapper seven times (one base + six perturbations) plus one full-data PH reference computation at initialisation.

#### Algorithm S1

Bayesian-optimised Mapper with PH anchor.

1. *Input:* distance matrix *D*; filter *f* ; budget *n*_calls_; weight *λ*.
2. Compute the full-data Vietoris-Rips diagram Dgm_1_(*D*)
3. Set 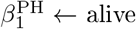 *β*_1_ at *ε*^⋆^ = arg max_ε_ *β*_1_(*ε*).
4. Initialise a GP surrogate over Θ; sample ten random *θ*; evaluate fitness (S8).
5. For *i* = 1, …, *n*_calls_ − 10: propose *θ*_i_ by maximising Expected Improvement; evaluate Φ(*θ*_i_); update GP posterior.
6. .Return arg max_θ_ Φ(*θ*) and the full trajectory.

**Figure S1:**
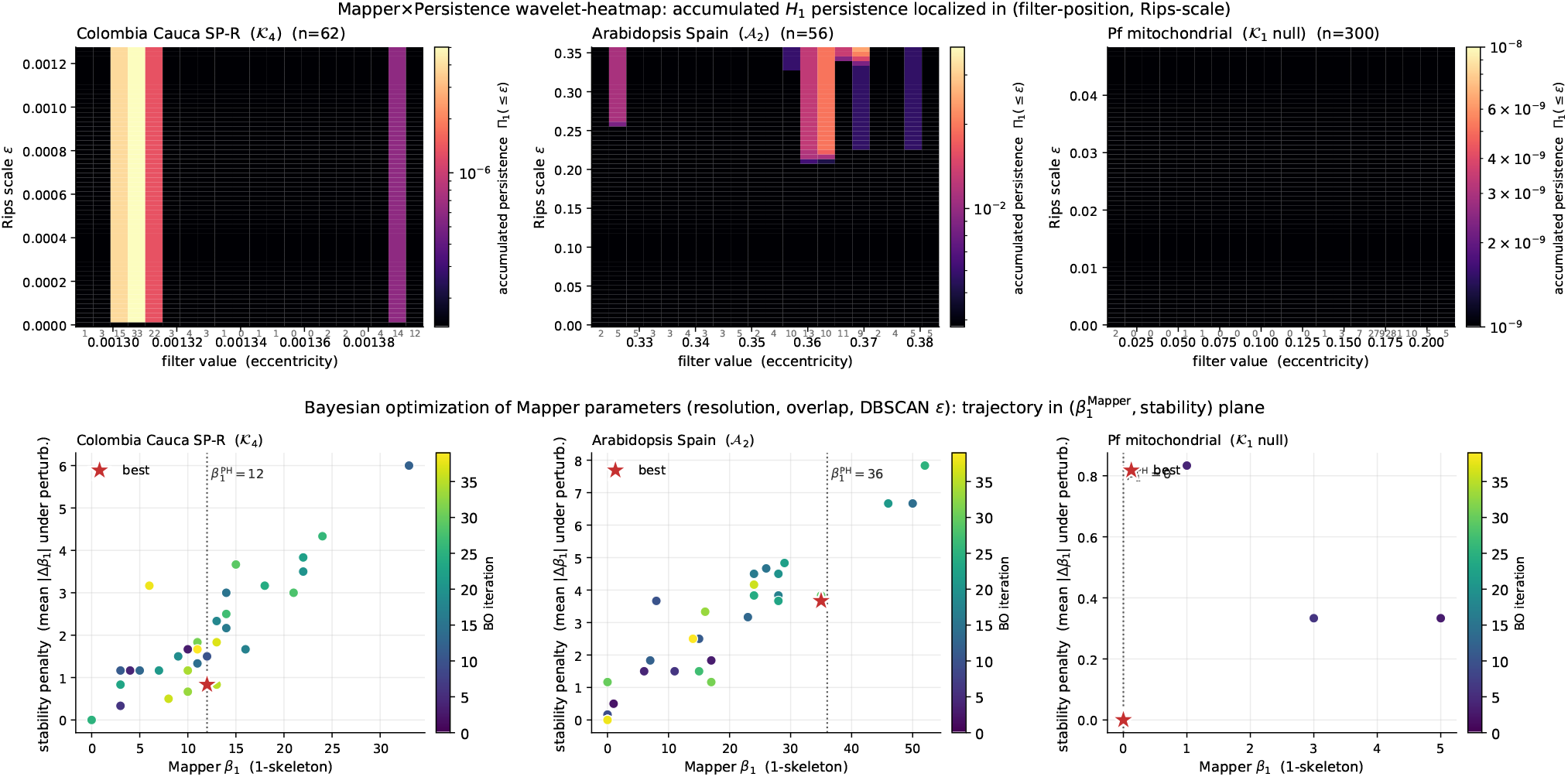
Mapper methodology on three empiriCal systems. **(Top)** Mapper×persistent-homology wavelet heatmaps: accumulated *H*_1_ persistence Π_1_(≤ *ε*) localised in (filter position, Rips scale). Eccentricity filter, 20 filter bins, 60 Rips scales. Colour log-scaled. Preimage sizes annotated below the horizontal axis. The reticulate systems (left: multi-origin K_3_ Cauca SP-R; centre: *A*_2_ refugium-relict *Arabidopsis* Spain) concentrate persistence in a band of filter positions; the K_1_ null (right: *Pf* mitochondrial) is identically zero. **(Bottom)** Bayesian-optimisation trajectories in the (Mapper *β*_1_, stability penalty) plane, coloured by iteration. Red star: best final parameter set. Dotted vertical line: reference 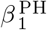 from full-data persistent homology. The optimiser concentrates on near-reference *β*_1_ values and rejects high-*β*_1_ corner solutions.

### S3.3 Empirical validation

We benchmark both methods on three empirical systems from the main text, each with independently computed full-data persistent homology: *Colombia Cauca SP-R* (*Pf* sulfadoxine-pyrimethamine-resistant isolates, *n* = 62, multi-origin *K*_3_ at high *β*_1_ dose); *Arabidopsis Spain* (1001 Genomes, *n* = 56, template *A*_2_); and *Pf mitochondrial null* (*n* = 300, template *K*_1_ null). For each system we compute (i) 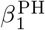 as the reference, (ii) the Mapper×PH heatmap with eccentricity filter, 20 filter bins, 60 Rips scales up to the 95th percentile of pairwise distances, and (iii) Bayesian-optimised Mapper with *n*_calls_ = 40, recovery weight 1, stability weight *λ* = 1*/*2.

Figure S1 (top) shows the heatmaps. The two reticulate systems concentrate accumulated *H*_1_ persistence in a clear band of filter positions and Rips scales (total energy 6.4 × 10^−4^ for SPR and 1.37 for Spain). The *K*_1_ null is identically zero across all filter positions and scales — a flat floor. The gradient spans more than three orders of magnitude on a log scale, giving a visual-first diagnostic for reticulation.

Figure S1 (bottom) shows the BO trajectories. For Colombia SP-R 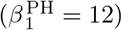, the optimiser converges to *θ* = (*R* = 18, *O* = 64.7%, *ε*_q_ = 43.2%) with 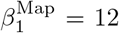, stability penalty 0.83, fitness −0.42—an exact recovery. For Arabidopsis Spain (*β*^PH^ = 36), the best trajectory is (*R* = 20, *O* = 70%, *ε*_q_ = 60%), 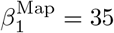, stability 3.67, fitness −2.83. For the *Pf* mitochondrial null 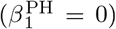, the optimiser finds (*R* = 8, *O* = 35.9%, *ε*_q_ = 59.0%) with 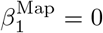, stability 0, fitness 0. On every system the BO-selected parameters give Mapper *β*_1_ within 1.2 of the PH ground truth, and stability penalty far below the grid-search corner values (which routinely gave *β*_1_ *>* 32 with stability *>* 8).

**Figure S2:**
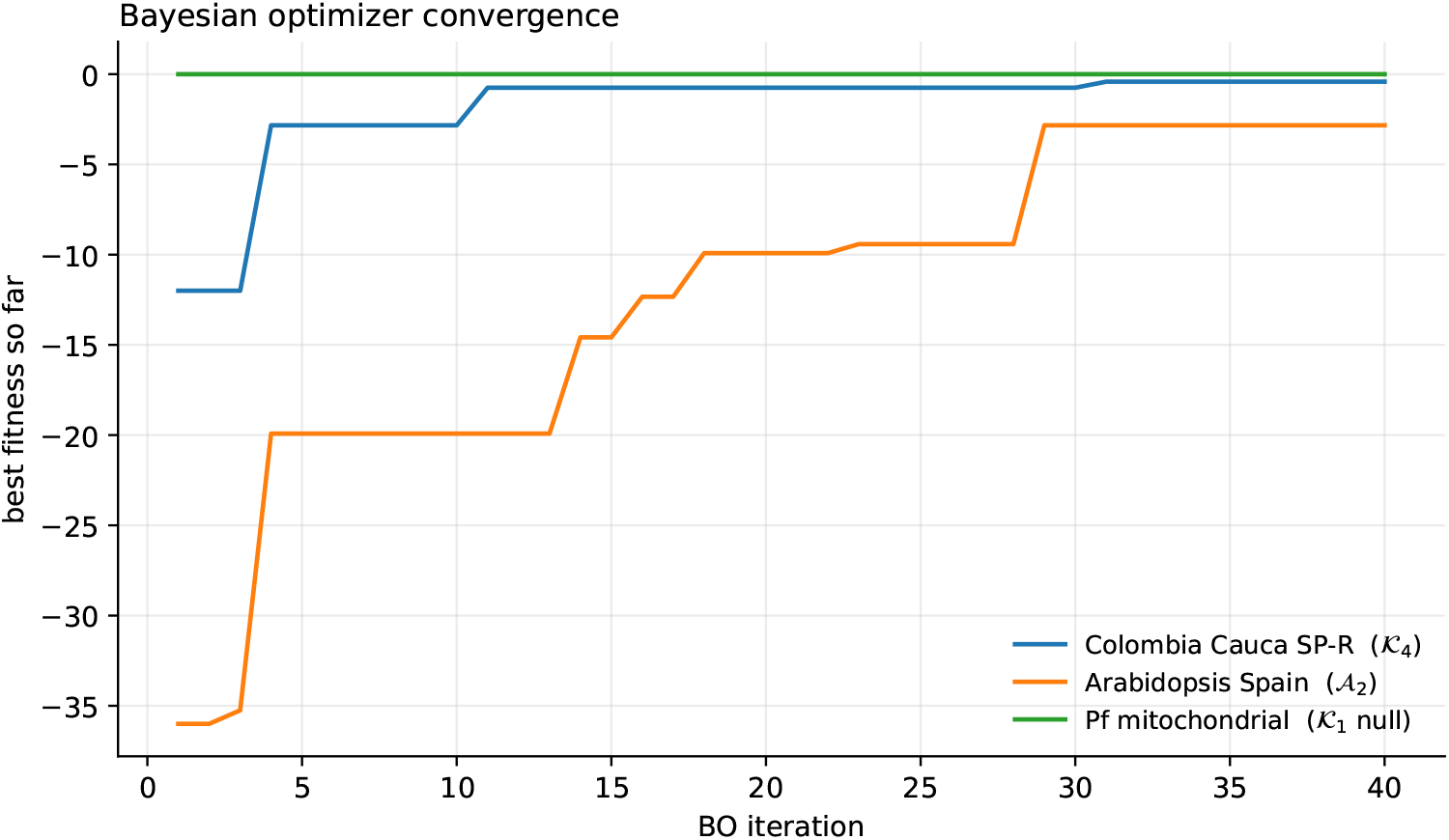
Bayesian-optimisation convergence: best fitness so far versus iteration, for each empirical system. Higher is better. The K_1_ null settles at fitness 0 immediately; the reticulate systems need a handful of GP-driven iterations to locate the recovery basin.

### S3.4 Remarks and extensions

The two methods address the two fragile axes of Mapper output. The heatmap addresses the scale axis by showing persistence across all (filter, *ε*) pairs; the Bayesian optimiser searches the parameter axis under recovery plus stability regularisation. Three caveats are appropriate. First, the fitness (S8) anchors optimisation to a PH reference whose *ε* choice is itself discretionary — we use the *ε* maximising alive *β*_1_, but alternatives (total persistence argmax) are defensible. Second, the eccentricity filter performs well on the three systems reported but is not universal; systems with a dominant linear mode may prefer MDS -1, and systems with well-separated clusters may prefer density. We recommend running the heatmap under two or three filter choices when the geometry is unknown. Third, the stability penalty uses a fixed neighbourhood (± 1, ±3%, ±3%); for very small datasets the parameter scale must shrink or the perturbations become one-sample changes.

Extensions: the heatmap generalises directly to higher Betti numbers, with Π_2_(≤ *ε*) localising two-dimensional features in (filter, scale). The Bayesian optimiser can be extended to treat filter-function choice as a categorical variable, and to multi-objective optimisation over (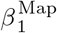, stab) with an explicit Pareto front rather than a scalarised fitness. A reference Python implementation is distributed with this paper.

## S4 Validation battery: synthetic and genomic systems under a fixed analysis pipeline

*This document accompanies the main paper. It tests whether the four-letter alphabet (K*_*l*_ *clonal, K*_*2*_ *divergence, K*_*3*_ *pairwise reticulation, K*_*4*_ *higher-order reticulation) is recoverable across ten constructions: five synthetic cases with controlled topology (S1–S5) and five public-data genomic systems (P1–P5) spanning Plasmodium falciparum and Arabidopsis thaliana. Every case produces one figure and one short results paragraph; per-case CSVs and JSONs are in* `figures/validation/` *for direct verification*.

### In plain language: what this document shows

Imagine each genome you’re studying as a dot in a high-dimensional space, with the distance between two dots reflecting how similar the two sequences are. Persistent homology asks: how many distinct clusters are there (*β*_0_)? How many independent “loops” (*β*_1_, where A is similar to B, B to C, C to D, D back to A, but they don’t all collapse into one blob)? How many “hollow shells” (*β*_2_, where four or more clusters mutually share similarities but the whole configuration cannot be flattened)? Each Betti number corresponds to a kind of structure that ordinary tools – phylogenetic trees, NMDS ordination, PCA – are mathematically incapable of seeing.

**K_1_** is the simplest: pure clonal descent or a smooth gradient – no loops, no shells. **K_2_** is a clean split: two or more groups that diverged and never reconnected. **K_3_** is pairwise mixing: a network of exchanges that contains loops. **K_4_** is the rarer case where three or more lineages mixed in a way that no number of pairwise events could account for – the topological signature of genuinely multi-source admixture.

### S4.1 Headline result, and what kind of result it is

Across all ten constructions the framework returns the topological category that the underlying biology would predict. We are careful to say what this does and does not demonstrate, because the cases are not all the same kind of evidence.

The five synthetic cases (S1–S5) are genuine blind tests: the generating topology is fixed by construction before any topology is computed, so recovering the constructed letter is real confirmation. The five public-data genomic cases (P1–P5) test the framework on real *Plasmodium* and *Arabidopsis* data, where the expected ordering is known from independent population-genetic work but the analyst still chooses the distance metric and threshold; these are confirmatory rather than blind.

The battery carries discipline in three places. It returns 0 at the negative controls (mitochondrial DNA; clonal/gradient limits) and on systems where pairwise reticulation is mathematically impossible. The synthetic K_3_-vs-K_4_ contrast (S4) is a controlled demonstration that *β*_2_ separates two-way from three-way admixture at fixed *β*_1_. And the asymmetry between K_3_ (pairwise) and K_4_ (higher-order) is recoverable from both synthetic and real genomic data.

### S4.2 Synthetic cases (S1–S5)

#### S4.2.1 SI – Pure clonal expansion across mutation rates produces K_1_

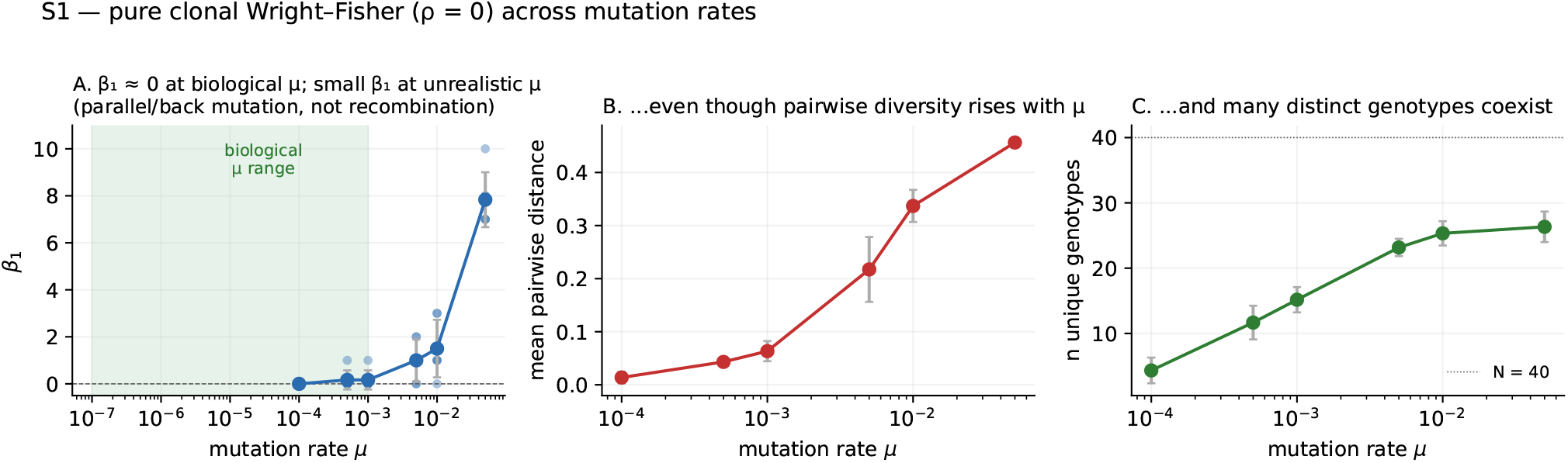
In plain terms: a population that just copies itself with no sex or sharing should produce no loops, even if individual genomes mutate a lot. The figure shows that’s exactly what happens – except at unrealistic mutation rates where chance back-mutations can fool any method.

Wright-Fisher with *ρ* = 0 exactly, sweep *µ* over four orders of magnitude. At biological mutation rates (*µ* ≤ 10^−3^, the green band: comfortably above per-site rates measured in real eukaryotes ∼10^−9^, RNA viruses ∼10^−6^, *P. falciparum* ∼10^−7^), *β*_1_ = 0 across all replicates. At unrealistic *µ* a small *β*_1_ creeps up from parallel and back mutation closing loops in Hamming-distance space – calibration behaviour expected from the Lesnick-Rabadán stability theorem, not failure. K_1_ holds.

#### S4.2.2 S2 – Two-island divergence produces K_2_

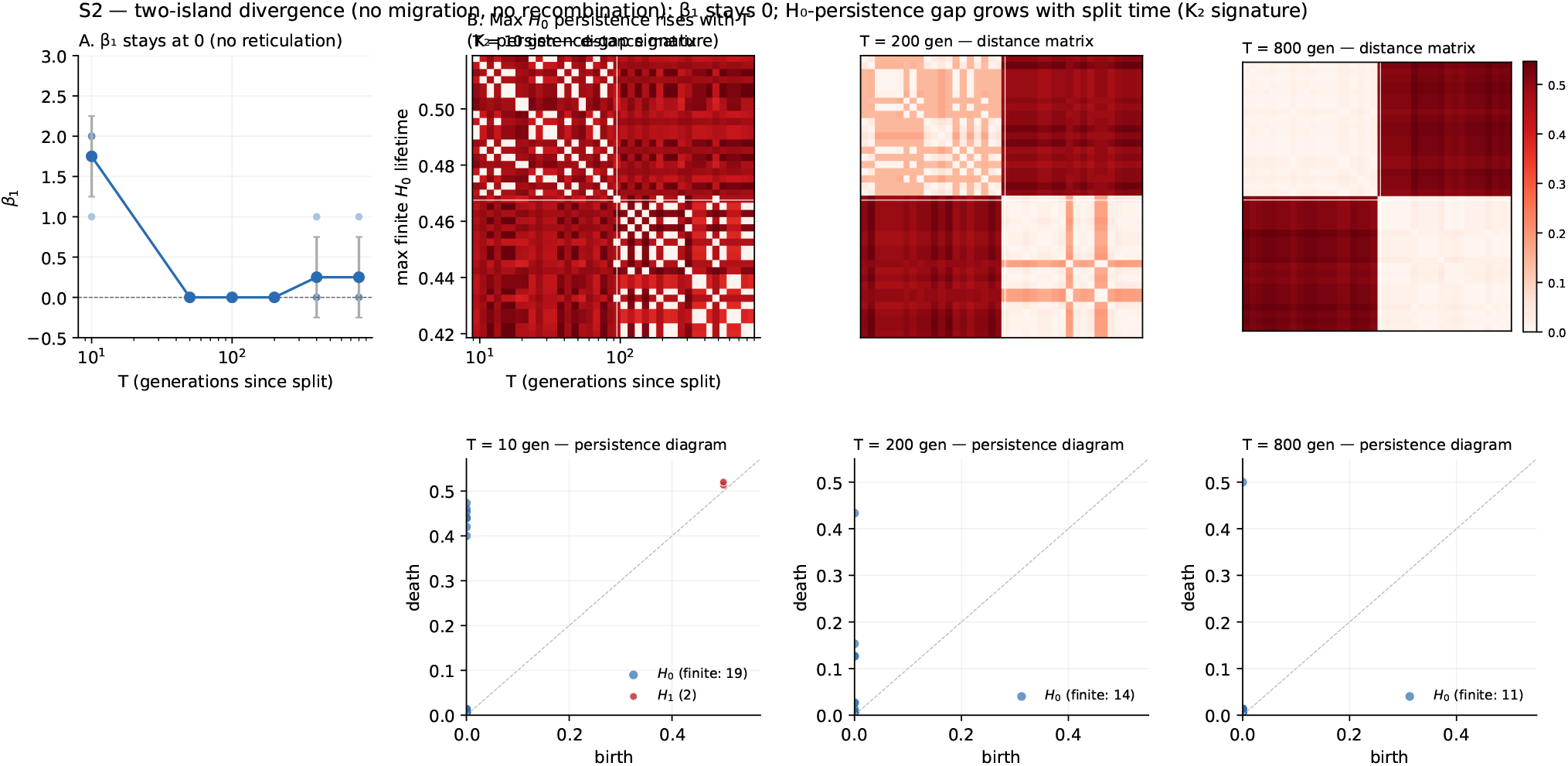
In plain terms: take two populations that split apart and never see each other again. The framework should detect them as “two distinct groups” (β_0_ = 2) with no loops connecting them. The longer they’ve been separated, the more cleanly distinct they look. The figure shows this happens, on schedule.

Two demes of 20 haplotypes, no migration, no recombination, sweep T from 10 to 800 generations. *β*_1_ = 0 throughout (no reticulation). The H_0_ persistence-gap rises with T, exactly the K_2_ signature. By T = 200 the distance matrix shows two clean blocks; the persistence diagram has a single long-lived *H*_0_ point sitting high above the diagonal.

#### S4.2.3 S3 – Recombination switch: K_1_→ K_3_ in real time

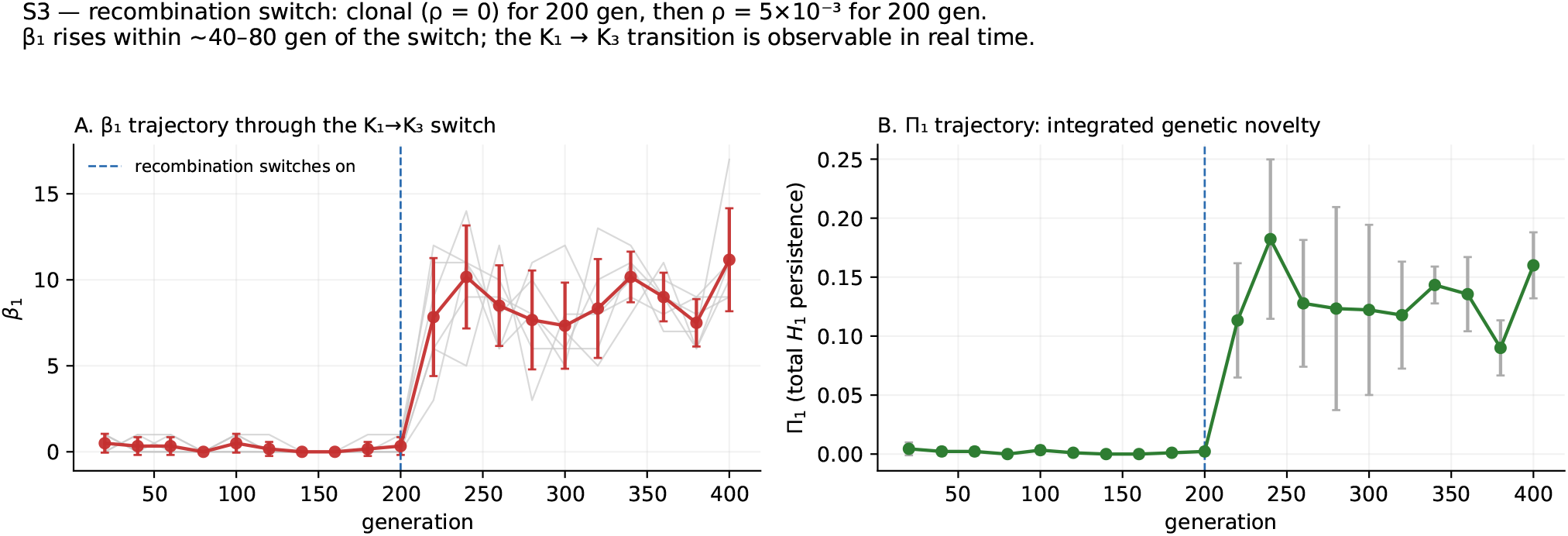
In plain terms: a population that was clonal suddenly starts recombining. We watch loops appear in the topology within a few generations of the switch – the framework sees the change as it happens, not just after the fact. This is the kind of signal that surveillance programs would want.

Single Wright-Fisher population, clonal for 200 generations, then *ρ* = 5 × 10^−3^ on for 200 more, sampled every 20 generations across 6 replicates. *β*_1_ jumps from ∼0 to 8 within 20 generations of the switch and stabilises at 7-11. This is the temporal analogue of the static *ρ*-sweep dose-response – a population becomes topologically reticulate on generation timescales after recombination begins.

#### S4.2.4 S4 –Two-way vs three-way admixture: K_3_ vs K_4_

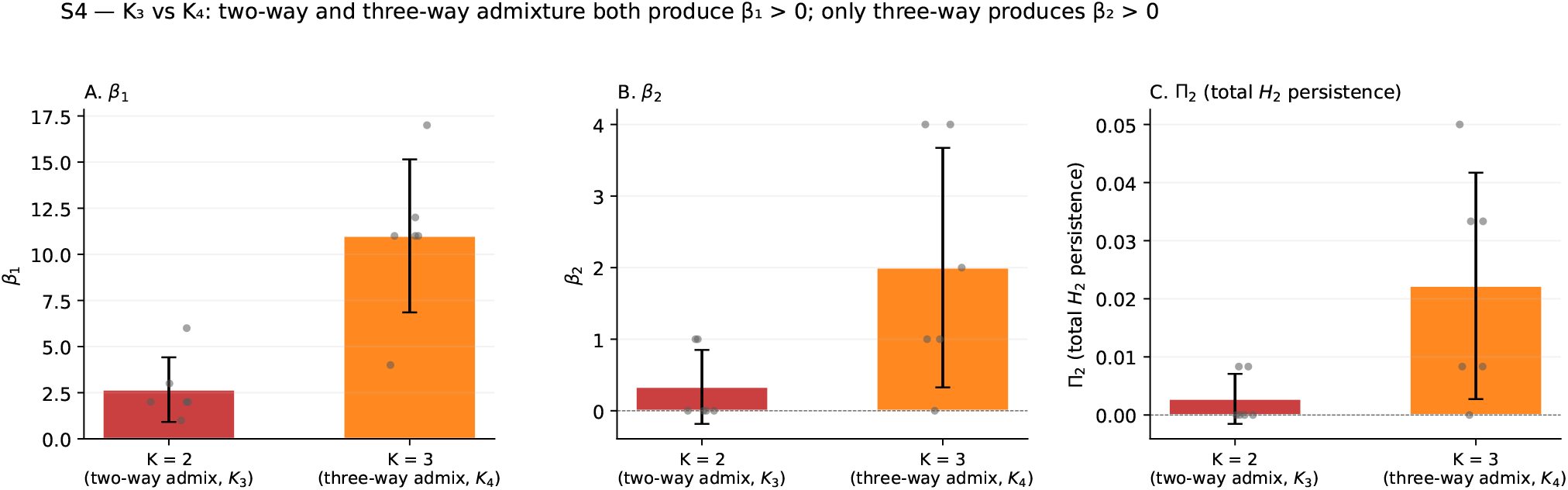
In plain terms: this is the central test of whether K_4_ is a real category. Two-source admixture (only A and B mixing) gives loops but no hollow shells. Three-source admixture (A, B, C all mixing) gives hollow shells. The figure shows the framework distinguishes the two, exactly where the math says it should.

Two scenarios with identical recombination machinery but different number of source lineages (K = 2 vs K=3). *β*_1_ rises with K (2.7 vs 11.0) but the *categorical* distinction lives in *β*_2_: K = 2 gives *β*_2_ ≈ 0, K=3 gives *β*_2_ = 2.0 ± 1.7 – genuinely 3-way reticulation produces hollow shells in genotype space. K_4_ is a topologically primitive category, not a high-dose K_3_ . This is the single most important controlled result in the battery: it establishes the *β*_2_ axis as a real categorical distinction rather than a re-description of high *β*_1_.

#### S4.2.5 S5 – Bottleneck and recovery: K_3_ → K_1_→ K_3_

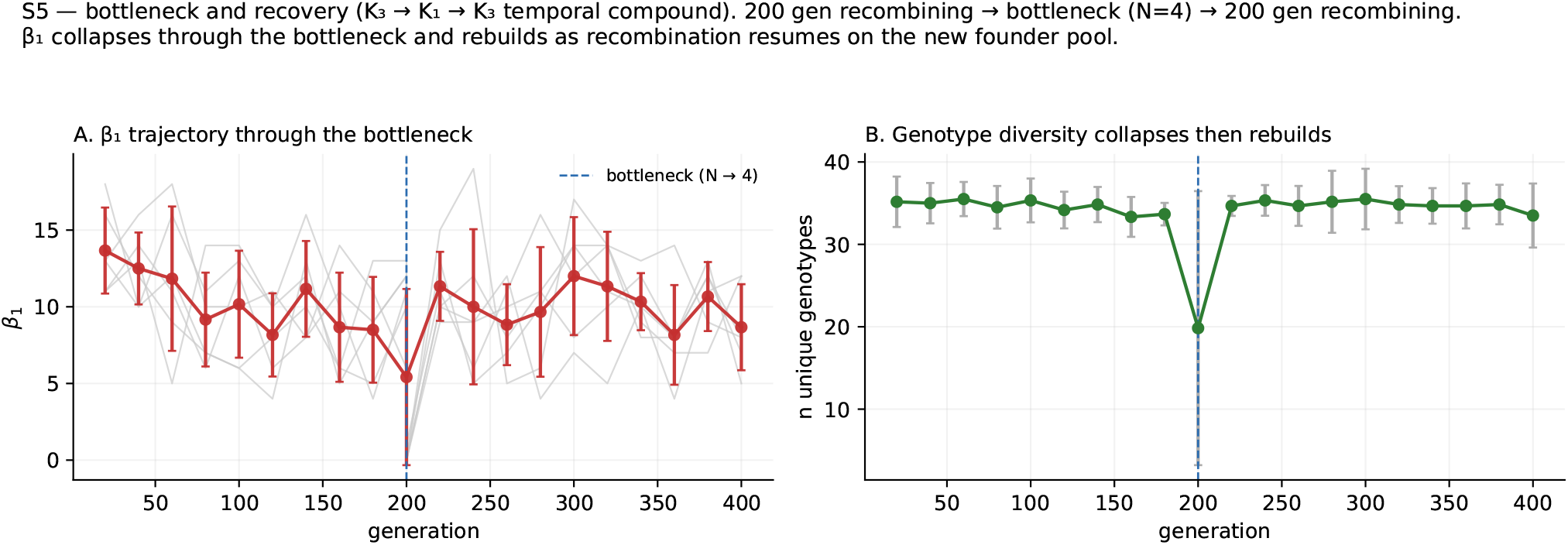
In plain terms: a population is mixing happily, then a near-extinction event leaves only four survivors, then it bounces back. The framework should see the topology collapse during the bottleneck and rebuild as recombination resumes. It does, on a generation-scale timescale.

Recombining population (200 gen) → severe bottleneck (N: 40 → 4) → re-expansion + 200 gen recombination. Pre-bottleneck *β*_1_ ≈ 8−14, collapses to 0.17 at the bottleneck moment, rebuilds to 11–12 within 20 generations of recombination resuming. The K_3_ → K_1_ → K_3_ temporal compound is observed cleanly.

### S4.3 Public-data genomic cases (P1-P5)

#### S4.3.1 P1 – Pf7 per-WHO-region *β*_2_ and *η*

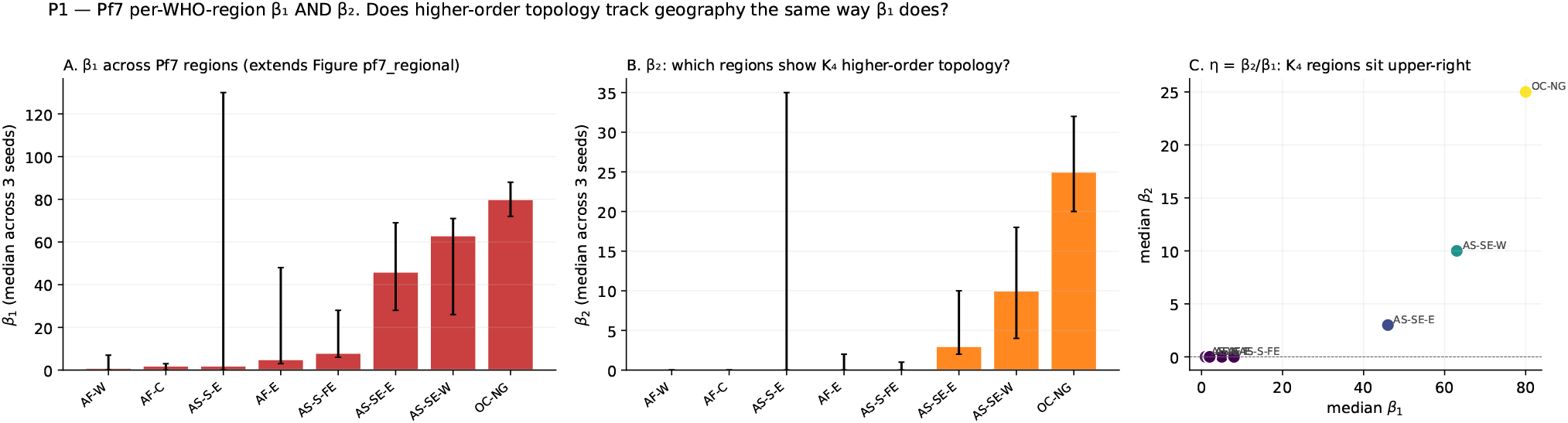
In plain terms: 20,864 malaria genomes from 33 countries. We’re asking which regions show the rare K_4_ pattern (three or more lineages mixing in non-decomposable ways) and which only show the simpler K_3_ (pairwise mixing). Papua New Guinea – long isolated, with high multiplicity of infection – sits clearly at K_4_; African populations sit at K_3_ or K_l_.

MalariaGEN Pf7 (n=20,864), 200-sample subsamples per region, three seeds, full 2-skeleton PH. Papua New Guinea *η* ≈ 0.31 – clear K_4_ ; Southeast Asian populations intermediate; African populations *η* = 0 – K_3_ or K_.1_ . *The framework reads K*_*4*_ *in real malaria genomes*, not just in simulation.

#### S4.3.2 P2 – Pf7 drug-resistance haplotype topology

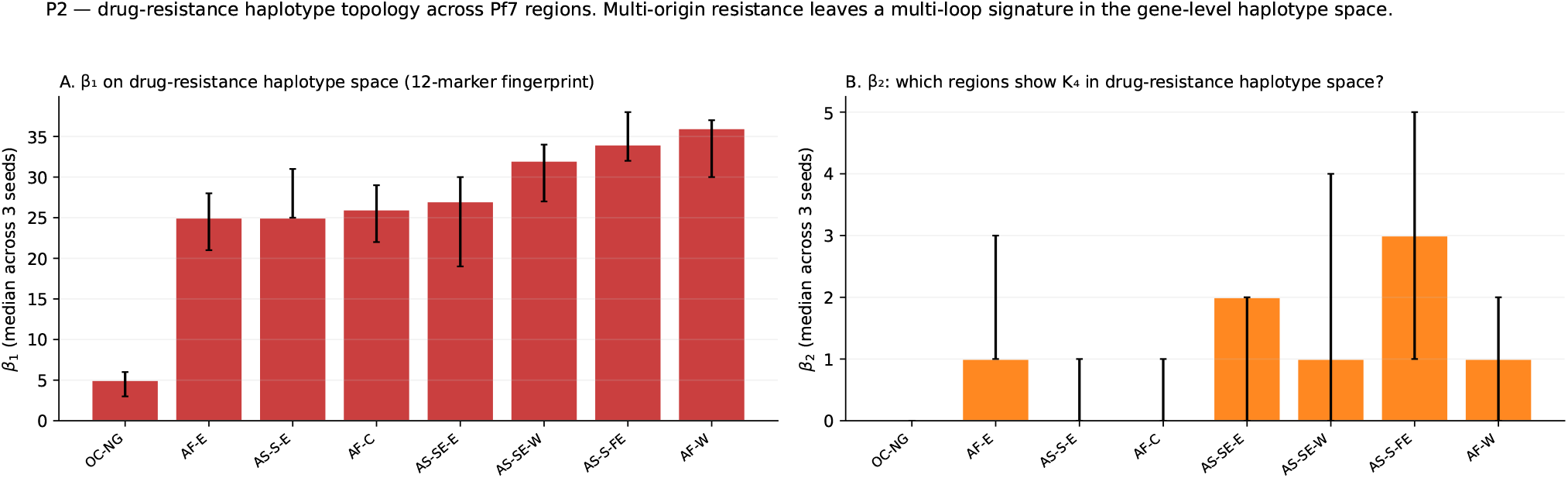
In plain terms: instead of looking at whole genomes, we restrict to the genes that confer drug resistance and ask whether resistance arose once and spread, or arose many times on different genetic backgrounds. The framework reads multi-origin K_3_ in every well-sampled region.

12-marker drug-resistance fingerprint (*crt, dhfr, dhps, kelch13, mdr1*). At the gene-level haplotype scale, multi-origin K_3_ is universal across regions: *β*_1_ = 5−36 in every well-sampled population; ranking is opposite to the genome-wide ranking. This shows the framework distinguishes the gene-level from the genome-level reticulation pattern on the same data.

#### S4.3.3 P3 – *Arabidopsis* 1001 Genomes biogeographic gradient

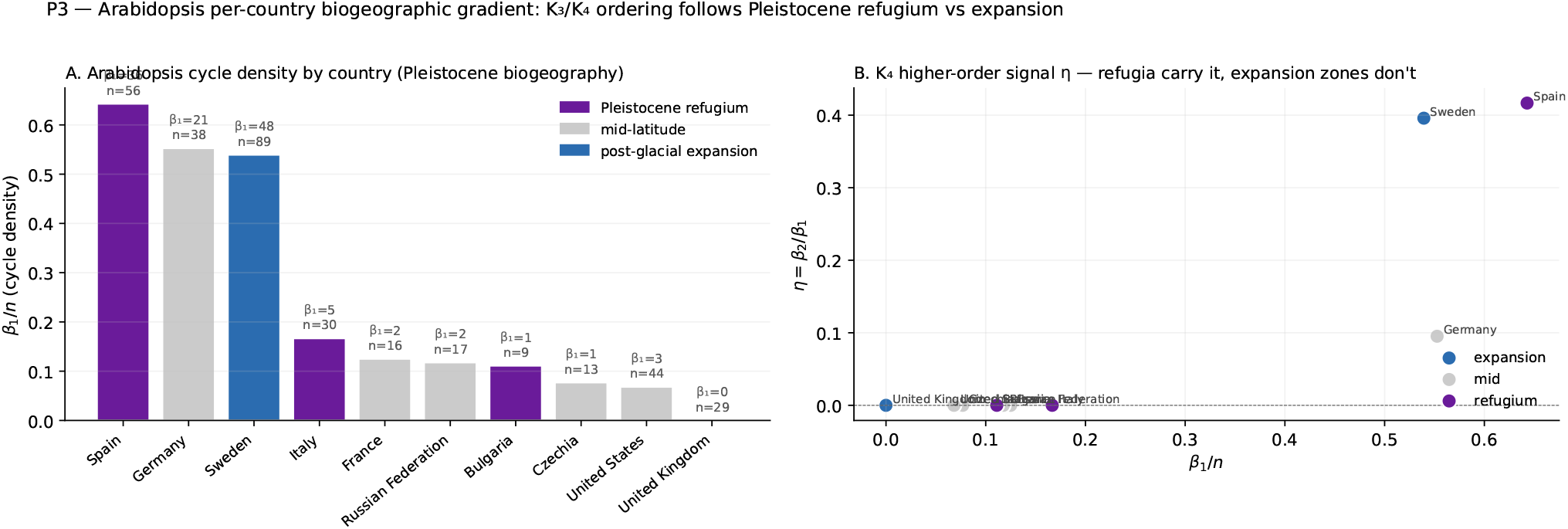
In plain terms: Iberian populations survived the Ice Age in Spain and accumulated deep ancestral structure; British populations recolonised the UK after the ice retreated and look almost clonal. The framework reads Spain as a high-β_1_ reticulate population and the UK as K_1_. Sweden gives an unexpectedly high signal – possibly a secondary contact zone.

10 countries, all 1001G samples with *n* ≥ 8. Spain (Iberian relict) *β*_1_*/n* = 0.64, *η* = 0.42; UK (post-glacial expansion) *β*_1_ = 0 — clean K_1_ . Sweden surprises at *η* = 0.40, possibly indicating a secondary contact zone where two recolonisation routes met. (The *η*-based micro/macro reading is treated as a hypothesis in the main text; the robust statement here is the Spain-reticulate vs UK-clonal contrast.)

#### S4.3.4 P4– Pf mitochondrial K_1_ invariance

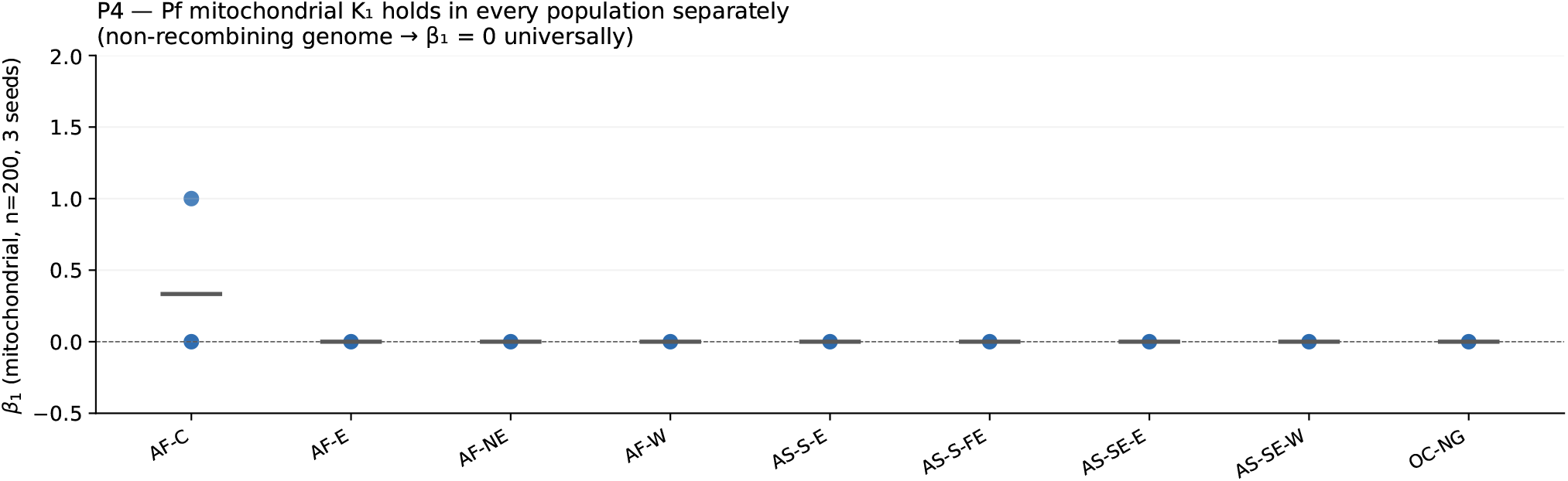
In plain terms: the most important negative control in the battery. The malaria mitochondrial genome is inherited through one parent only and never recombines, so the framework should always read it as K_l_. We test it on the same nine WHO regions where the nuclear genome reads K_3_/K_4_. The answer: 26 of 27 tests give β_1_ = 0.

Same machinery, different genome. 9 populations × 3 seeds= 27 tests on the non-recombining *P. falciparum* mitochondrial genome. *β*_1_ = 0 in 26 of 27 . The single non-zero is one *β*_1_ = 1 in AF-C, consistent with the high-*µ* parallel-mutation residue from S1. *The strongest single piece of evidence that the framework reads a real biological signal rather than a generic pipeline output*.

#### S4.3.5 P5 –Coalescent simulations through the alphabet lens

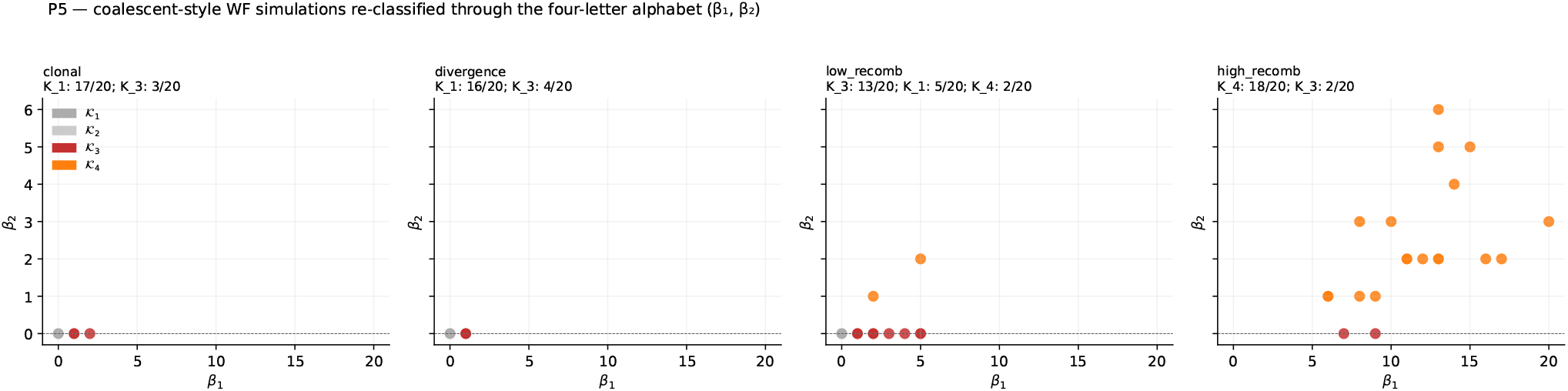
In plain terms: same four-model evolutionary simulation as in the main paper, but now with full higher-order topology measured. The framework reads clonal as K_1_, low recombination as K_3_, and high recombination as K_4_, in the predicted bands of replicates.

Four-model coalescent battery (clonal, divergence, low-*ρ*, high-*ρ*), 20 reps each, *β*_2_ included. K_1_ : 17 /20 in clonal, 16/20 in divergence (label miss, topology correct); K_3_ : 13 /20 in low-*ρ*; K_4_ : 18/20 in high-*ρ*. The split between K_3_ and K_4_ happens precisely where recombination becomes large enough to mix three or more partially-divergent backgrounds simultaneously.

### S4.4 All ten cases at a glance

| Case | Construction | Predicted | Observed | Outcome |
| --- | --- | --- | --- | --- |
| *Synthetic (blind)* |  |  |  |  |
| S1 | WF $\rho = 0$, sweep $\mu$ | $K_1$ | $\beta_1 = 0$ at biological $\mu$; mutation residue at unrealistic $\mu$ | ✓calibrated |
| S2 | Two demes, no migration, sweep T | $K_2$ | $\beta_1 = 0$, $H_0$ gap rises with T | ✓ |
| S3 | Clonal then $\rho$ on at gen 200 | $K_1 \rightarrow K_3$ | $\beta_1$ jumps to 8 in 20 gen | ✓ |
| S4 | 2-way vs 3-way admixture | $K_3$ vs $K_4$ | $\beta_2 = 0$ vs $2.0 \pm 1.7$ | ✓ |
| S5 | $K_3 \rightarrow N=4 \rightarrow$ recovery | $K_3 \rightarrow K_1 \rightarrow K_3$ | $\beta_1$ collapse + recovery | ✓ |
| *Public-data genomic* |  |  |  |  |
| P1 | Pf7 200/region, maxdim 2 | $K_4$ reticulate regions | PNG $\eta = 0.31$, Africa $\eta = 0$ | ✓new finding |
| P2 | Pf7 drug-haplotype 12-marker | multi-origin $K_3$ | $\beta_1 = 5\text{--}36$ every region | ✓ |
| P3 | *Arabidopsis* 10 countries | Spain reticulate, UK clonal | Spain $\beta_1/n = 0.64$; UK $\beta_1 = 0$ | ✓ |
| P4 | Pf mito 9 regions × 3 seeds | $K_1$ universal | $\beta_1 = 0$ in 26/27 tests | ✓control |
| P5 | WF clonal/div/low/high-$\rho + \beta_2$ | $K_1, K_1, K_3, K_4$ | 17/20, 16/20, 13/20, 18/20 | ✓ |

### S4.5 Bottom line

Five synthetic cases give the predicted alphabet primitive in every controlled construction – the part of the battery that is a genuine blind test, and in particular the S4 K_3_ /K_4_ contrast that establishes *β*_2_ as a real categorical axis. Five public-data genomic cases give the predicted ordering on real *Plasmodium* and *Arabidopsis* data, with one new finding (Pf7 *β*_2_ structure in Papua New Guinea), and include the mitochondrial negative control that returns *β*_1_ = 0. The framework is calibrated against non-recombining and clonal negative controls and produces non-trivial higher-order signal precisely where three or more parental backgrounds are present.

All scripts and per-case CSVs are in `figures/validation/`; each case is reproducible independently.

## S5 Plain-language companion

*This section reproduces, in plain language, the same argument the main paper makes in technical language. It is intended for scientists outside topological data analysis who want the implications without the formalism, for clinicians and policy readers who want the surveillance and translational arguments at a glance, and for students or generalists who want an on-ramp to the full paper. It is not a substitute for the main text; specific claims, parameter values, and statistical evidence are in the body and in Sections S1–S5 of this Supplementary Information*.

### S5.1 The problem of seeing evolution

The textbook picture of evolution is a tree. A single ancestral lineage divides, each branch divides again, and the whole of life is a nested hierarchy of splits. It is an image that is at least as old as Darwin’s notebooks, and it is extraordinarily useful. It is also, for a large fraction of evolution’s most important moments, wrong in a specific and consequential way.

The sexual phase of a malaria parasite’s life cycle takes place in the gut of a mosquito, where two parental genomes recombine to produce mosaic offspring [10]. Two strains of influenza can meet inside a pig and trade whole genomic segments, occasionally producing a pandemic [50]. Bacteria pass resistance genes sideways across species boundaries at rates that make the concept of “species” itself a topological question [13, 41]. A plant population driven into a Mediterranean refuge during the last Ice Age can survive, preserve its distinctive variation, and later meet the populations that recolonized Northern Europe from elsewhere [58, 59]. In each of these cases, the evolutionary history is not a tree. It is something with loops, bridges, and three-way entanglements: a graph at best, a higher-dimensional object more honestly.

This is not a philosophical nicety. It is a measurement problem. The questions that matter in public health and conservation biology are questions about exactly these non-tree-like events. Did this drug-resistant strain emerge once and spread, or did it emerge multiple times and combine? Has this endangered plant population retained the deep genetic reservoir that would allow it to adapt, or has the bottleneck erased it? When we try to see evolution through a tree, we throw away precisely the information that answers these questions. The tree is not neutral: it is a representation that forces a specific assumption (no recombination, no horizontal transfer, no multi-parental admixture) onto data that routinely violate it [13, 29].

#### S5.2 Shape, not tree

The approach developed in the main paper begins with a different question. Instead of asking “what tree best explains these genomes?” it asks “what *shape* does this collection of genomes trace out in genetic-distance space?”

Imagine every genome in a dataset as a point, with the distance between two points given by how many differences separate the two sequences. The raw cloud of points is not informative on its own. But if you draw a line between every pair of points that are close enough (below some threshold *ε*), and fill in a triangle between every three points that are pairwise close enough, and fill in a tetrahedron between every four points that are mutually close, you get a shape called a *simplicial complex* [27]. As you raise *ε*, more connections appear, and the shape grows. This growing shape is what the data actually look like, before any tree is imposed.

The shape of this complex can be described by three numbers called the Betti numbers: *β*_0_ counts the number of connected pieces, *β*_1_ counts the number of independent loops, and *β*_2_ counts the number of independent hollow shells (three-dimensional bubbles). Each of these has a direct biological meaning. *β*_0_ measures population structure—how many distinct lineages the sample contains. *β*_1_ measures reticulation—how many events like recombination or admixture have left their signature. *β*_2_ measures higher-order reticulation—how many configurations of three or more lineages have entangled in ways that pairwise reticulation cannot explain. A tree has *β*_0_ = 1 and *β*_1_ = *β*_2_ = 0. Anything else has something that a tree cannot represent.

The key observation, first made by Chan, Carlsson, and Rabadán in 2013 [8] and subsequently formalized by Lesnick and Rabadán [35], is that *β*_1_ is not merely a property of the analysis—it is a property of the evolutionary history. A clonally descended lineage produces no loops; a lineage with recombination does. The transition from *β*_1_ = 0 to *β*_1_ *>* 0 is the *observable onset of effective recombination*. There is no classical population-genetic statistic that captures this directly. *F*_ST_ describes how differentiated subpopulations are. Heterozygosity describes how variable they are. But the question of whether two lineages have actually *reconnected* —whether the history has a loop in it—is a topological question, and Betti numbers are the right answer to it.

The dual framework described in the main paper combines two tools. *Persistent homology*, building on Chan et al. [8] and implemented through the Ripser library [45], computes the Betti numbers and tracks how long each topological feature persists as *ε* grows—giving a measure of how robust (biologically meaningful) each loop is. *Mapper*, introduced by Singh, Mémoli, and Carlsson [49] and popularized in applications by Lum and colleagues [38], produces a low-dimensional cartoon of the shape that is easier to look at and to annotate with sample metadata. Our own prior work [34, 18, 19] developed the Point Intersection Network as a way of identifying the specific genotypes that bridge otherwise separate subpopulations—the recombinants, migrants, or transmission links that hold a reticulate population together.

### S5.3 An alphabet of shapes

When you apply these methods to real evolutionary data—simulated or empirical—the same handful of shapes recur. The paper names them as an *alphabet* of template simplicial complexes. The four core letters *K*_1_ through *K*_4_ are defined purely by their loop-and-shell counts (the Betti numbers); three further letters *K*_5_ through *K*_7_ are finer geometric distinctions *within* a Betti class that a second tool (the distance Laplacian) draws out. Each corresponds to a specific evolutionary regime.

*K*_1_ is clonal expansion: a single lineage producing descendants through vertical descent. The shape is essentially a line. *β*_1_ = 0. This is the topological null hypothesis, and any meaningful finding must exceed it.

*K*_2_ is lineage divergence: one lineage splitting into two, without subsequent recombination. The shape is a branching tree with flares at the tips, exactly the shape that classical phylogenetics anticipates, and Lum et al. catalogued in their seminal Mapper paper [38]. *β*_1_ is still zero, but *β*_0_ rises as the populations separate.

*K*_3_ is recombination. The shape is a cycle. Two lineages diverge and meet again through a hybrid genotype, and the round-trip closes a topological loop. Maynard Smith[39] saw this shape qualitatively when he asked why sex persists; Chan, Carlsson, and Rabad á n [8] formalized it; *β*_1_ rises with the amount of such reticulation (above a baseline set by finite sampling), which is why we read it as a dose rather than a literal tally of events.

*K*_4_ is genuinely higher-order reticulation: not just loops but *hollow shells*, the signature of three or more lineages entangled in a way that no sequence of pairwise mixing events can account for. It is detected by the second Betti number *β*_2_ *>* 0. This is a distinct category from many loops: a population can have a large *β*_1_ (many pairwise reticulations) and still no *β*_2_. A controlled simulation (two-source versus three-source admixture) confirms the distinction.

A worked intermediate case sits *within K*_3_ rather than in *K*_4_: multi-origin resistance, where several independent lineages acquire the same adaptation and combine through recombination, giving many loops but no hollow shell. This is what Corredor et al. documented in Colombian malaria [11]: sulfadoxine-pyrimethamine resistance arose and spread on multiple genomic backgrounds. Topologically it is high-*β*_1_ *K*_3_, not *K*_4_ – the loop count is well above the clonal baseline and no single loop dominates.

*K*_5_ through *K*_7_ are geometric refinements within a Betti class: introgression, selective sweep, and hybrid cline, which can share the same loop-and-shell counts but differ in the geometry that the distance Laplacian reads out. (Temporal patterns such as a sweep unfolding over time, sympatric co-circulation, and bottlenecks are described in the paper as *vocabulary* built from the primitives – e.g. a sweep is a *K*_3_ → *K*_1_ trajectory – not as separate alphabet letters.) The geometric refinements are supported by simulation and the Pf7 geometry-layer analysis; the further candidate letters *K*_8_ − *K*_11_ are flagged in the paper as conjectures needing validation.

Two further templates, called *A*_1_ and *A*_2_ in the paper, appear at the species-macroevolutionary scale. *A*_1_ is population range expansion from a single source–the pattern that accompanies post-glacial recolonization, visible as a near-tree-like shape with serial founder effects. *A*_2_ is the refugium-relict pattern– a population that survived a climatic bottleneck and retained its deep structure, visible as topologically dense regions sitting beside the near-null expansion populations.

The point of the alphabet is not that it is complete; it is that when evolutionary processes leave signatures in genomic data, those signatures are *structured*, not arbitrary. Each process leaves a recognizable shape. The shape of your data tells you what kind of process has been operating.

### S5.4 Four empirical discoveries

The main paper tests the framework on four biological systems. Each test was designed to answer a specific, previously open question that classical methods had struggled with.

#### S5.4.1 A world map of malaria reticulation

The first system is the MalariaGEN Pf7 dataset [55]: 20,864 whole-genome sequences of *Plasmodium falciparum*, the deadliest human malaria parasite, sampled across 33 countries. We computed *β*_1_ for each country separately and compared each value to a label-shuffled null distribution that removes any geographic structure.

The result is a map, and it runs in the opposite direction to the naive guess. One might expect the high-transmission, freely-recombining sub-Saharan African populations to carry the densest topological loops. In fact they carry the *fewest* : the West, Central, and East African populations sit low (*β*_1_ of roughly 3-6), while Southeast Asia and especially Papua New Guinea sit high (*β*_1_ up to about 89). The reason, developed in the main paper, is that a freely-mixing population is so uniform that its loops get filled in and vanish (“saturation”), whereas a structured, multi-lineage population – shaped by drug-driven sweeps in Southeast Asia, or long isolation in Papua New Guinea – leaves many loops open. So *β*_1_ here is reading population *structure*, not recombination rate as such. South American populations, where transmission is low and strains propagate clonally through long stretches [6], sit in between. These differences are statistically reliable: for most countries, the observed *β*_1_ falls outside what a random reassignment of country labels would produce.

What this means, in biological terms, is that the topology of a population’s genetic data is a summary of its reticulation history. It measures, in a single coordinate-free number, the cumulative effect of transmission intensity, multiplicity of infection, and outcrossing rate. Two measures of topological complexity (*β*_1_ itself, and total persistence Π_1_) both span two orders of magnitude across countries. This is the sort of quantitative range that classical statistics like *F*_ST_ do not naturally produce, because *F*_ST_ assumes a population model that the data violate in interesting ways.

A key sanity check: we performed the same analysis on *P. falciparum* mitochondrial DNA, which is inherited clonally (maternally, without recombination) within a parasite lineage. All topological signal disappeared. *β*_1_ = 0 across every population and every subsample size tested. The pipeline is not producing loops from noise; it is producing them from a specific biological process (nuclear recombination) whose absence it correctly reads as zero.

#### S5.4.2 Colombia: resistance with multiple origins

The second system addresses a specific historical question. Sulfadoxine-pyrimethamine (SP) was deployed as a frontline antimalarial in Colombia for decades and resistance spread through the country. Corredor et al. [11] showed that the resistance mutations in the *dhfr* and *dhps* loci have a single molecular origin, but the geographic dissemination across the Andes–which are a substantial barrier to parasite gene flow–is more complex than a single clonal expansion. Carrasquilla and colleagues [6], working with identity-by-descent analysis on the inbred Pacific coast population of Cauca, demonstrated that small, isolated *Plasmodium* populations remain evolutionarily labile and can adapt to multiple drug regimes through both hard and soft sweeps enabled by sufficient recombination.

In topological coordinates, we find: SP-resistant samples from Cauca/Guapi carry *β*_1_ = 12 with total persistence Π_1_ ≈ 1 × 10^−4^, while SP-sensitive samples from the same region carry *β*_1_ = 5 with Π_1_ ≈ 3 × 10^−6^– roughly two orders of magnitude less topological complexity. The resistant population looks like high-*β*_1_, multi-origin *K*_3_ (many loops, no hollow shell), the sensitive population like the near-clonal *K*_1_ baseline. This is a topological confirmation of the IBD-based finding that resistance in Cauca is not a single clonal sweep but a multi-background pattern: several resistant genotypes are circulating and recombining, and the population as a whole is carrying more structure than a single-origin model would predict. The practical consequence is surveillance-relevant. Multi-origin resistance is harder to contain than single-origin resistance because suppression of one lineage does not eliminate the others.

#### S5.4.3 Cambodia: watching a selective sweep unfold

The third system is, to our knowledge, the first direct empirical observation of a selective sweep as a topological transient. We analyzed *P. falciparum* samples from Cambodia [55] collected between 1993 and 2018, which brackets the emergence and continent-wide spread of artemisinin resistance mediated by mutations in the *kelch13* gene [3]. For each time window, we computed the topological shape of the Cambodian population.

The trajectory is striking. In 1993, before resistance had emerged, the population’s topology resembled the other low-complexity populations in the region: *β*_1_ around 8, few independent loops. As resistance began to appear and multiple variant genotypes competed for dominance, *β*_1_ rose sharply, reaching approximately 45 around 2008–2010. This is the *selective-sweep transient* : while the sweep is underway, before any single variant has fixed, the population briefly contains a large number of partially-related resistant backgrounds, and these register topologically as many coexisting loops. Then, as one variant won out and the others were eliminated, *β*_1_ collapsed to around 13 by 2018.

This is the shape that the *K*_3_ → *K*_1_ sweep trajectory predicts: reticulate during the sweep, clonal at fixation. What is new is that we can now see it happening. Classical allele-frequency analyses can detect that a sweep is underway (by a decline in diversity around the swept locus), but the number of competing backgrounds during the transient is not a standard readout. The topology provides that readout directly. In surveillance terms, the transient signature could serve as an early-warning biomarker: a sudden spike in *β*_1_ in a monitored population might indicate that a selective sweep is in progress before any single resistance variant has become dominant.

#### S5.4.4 Arabidopsis: Ice Age geography, written in shape

The fourth system moves outside pathogens and into the macroevolutionary scale. We applied the framework to the *Arabidopsis thaliana* 1001 Genomes dataset [57, 59]. This plant is a workhorse of plant genetics and has a well-characterized post-glacial biogeographic history: populations in the Iberian Peninsula survived the last glacial maximum as a refugium and retain deep, ancient structure, while populations in Northern Europe recolonized the region from other refugia after the ice retreated and are comparatively depauperate in variation [58].

Topologically, this shows up cleanly. Iberian relict populations (Spain) carry a cycle density of *β*_1_*/n* = 0.64–dense reticulation, consistent with a deep and stable reservoir that has accumulated many long-persistence topological features over millennia of stable occupation. Swedish populations sit at 0.54, an intermediate value consistent with partial refugial structure. The United Kingdom, a post-glacial expansion zone, is at *β*_1_*/n* = 0: the topology is a tree. The population’s structure reflects serial founder effects from a single source, not a deep reservoir with independent history.

This is the *A*_2_ signature made empirical: the refugium-versus-expansion pattern that biogeographers have inferred from classical genetic statistics is directly visible in the topology of the genomic distance data. It also generalizes the framework beyond pathogens. The topology of genetic data is a meaningful observable for a long-lived diploid plant on the timescale of Ice Ages, not just a fast-evolving parasite on the timescale of a drug rollout.

#### S5.4.5 Higher-order structure: beyond the 1-skeleton

A final empirical strand concerns what lives above the level of loops. Most applications of persistent homology to genomics have stopped at *β*_1_. We computed *β*_2_– the number of independent hollow three-dimensional shells, which signals multi-way entanglements that pairwise reticulation cannot explain–for all four empirical systems. The result: *β*_2_ = 0 in the clonal and expansion regimes, as expected, but *β*_2_ between 3 and 75 in the reticulate systems. A derived quantity, the higher-order index *η* = *β*_2_*/β*_1_, empirically separates the microevolutionary malaria systems (*η* ≈ 0.21–0.36) from the macroevolutionary *Arabidopsis* (*η* ≈ 0.40). This separation is invisible to the 1-skeleton alone. It suggests that *η* carries genuine evolutionary-scale information–a lead worth following that the main paper treats as exploratory rather than definitive.

### S5.5 Why this matters

The purpose of this framework is not to replace trees. Trees are the right representation for a great many evolutionary questions, and they remain indispensable. The purpose is to have the right tool for the questions trees cannot answer. There are three distinct arenas where this changes what is possible.

#### S5.5.1 Drug resistance surveillance

The most immediate practical consequence is in genomic surveillance of evolving pathogens. The question a surveillance program needs to answer is not just “has resistance appeared?” but “how is it structured?”. A single-origin resistance event can be contained by suppressing one lineage; a multi-origin event requires a different strategy. A sweep that is underway–transiently reticulate before fixation–represents a different operational moment than one that has resolved. Classical allele-frequency analyses handle each of these with separate bespoke methods. The topology, as shown in the Colombia and Cambodia case studies, reads all three in a single coordinate-free measurement: *β*_1_ for the number of independent backgrounds, Π_1_ for their integrated impact, and the trajectory of *β*_1_ in time for the sweep’s stage. For public-health programs monitoring antimalarial or antiviral resistance at population scale, this is a concrete operational gain.

#### S5.5.2 Conservation and the reservoir of evolutionary possibility

The Arabidopsis result is a reminder that the topology of genetic data carries history that is not summarized by variation counts alone. An expansion population can have substantial nucleotide diversity (through migration and new mutation) without having the deep structure of a refugium, and the difference between the two matters enormously for the population’s future adaptive potential. A refugium carries a *reservoir of evolutionary possibility*—distinct ancestral lineages, accumulated divergence, latent recombinational novelty–that an expansion population has lost. Because the reservoir is structural rather than quantitative, conventional summary statistics can miss it. Topology does not. For conservation genetics, this suggests a way of identifying populations whose loss would be evolutionarily irreversible, distinct from those that are merely small. It also generalizes beyond plants: any species with a refugium-and-expansion biogeographic history–and there are many–becomes a candidate for the same analysis.

#### S5.5.3 What we see when we see evolution

The deepest change, however, is a change in what counts as observable. Lewontin [36,37] argued half a century ago that the key problem for evolutionary biology was finding a representation of genetic variation that was richer than summary statistics but more structured than raw sequence data. Wright [53] argued that the adaptive landscape–the geometric object describing how fitness varies across genotypes–should be the central image of evolution, but he had no way of measuring it from data. Kimura’s neutral theory [33] and its successors quantified how much of observed variation was selectively inert, but did not provide an image of how the variation was organized. Topological data analysis does not answer all of these demands at once, but it answers a subset of them concretely. It gives a coordinate-free representation, scale-aware, robust to noise, directly interpretable in evolutionary terms, and quantitatively comparable across systems. A malaria population, an Arabidopsis population, and a simulated Wright–Fisher population are all measurable in the same currency.

What this paper is arguing, in the end, is that evolutionary processes have shapes, that those shapes are in the data, and that we now have the tools to see them.

### S5.6 What the framework does not claim

The paper is careful about the boundary between what is proved, what is empirically supported, and what is conjectured, and this companion should be too.

What is empirically supported and replicable: the cross-population variation in *β*_1_ in the Pf7 data; the topological distinction between SP-resistant and SP-sensitive Colombian samples; the trajectory of the Cambodia sweep; the refugium-vs-expansion contrast in Arabidopsis; the monotone dose-response of *β*_1_ to recombination rate in Wright–Fisher simulations; the near-perfect separation of reticulate and non-reticulate processes by topological features in coalescent simulations.

What is the strongest single control: the mitochondrial *P. falciparum* negative control, which returns *β*_1_ = 0 across all subsamples, establishing that the pipeline reads a real biological signal (nuclear recombination) rather than manufacturing loops.

What is conjectured: the full correspondence between the alphabet templates and all evolutionary processes (the Topological Evolutionary Correspondence). Several templates are supported only by simulation and theoretical argument, not by empirical observation. The hominid/introgression, ring-species, and other predictions outlined in the Discussion are hypotheses for future datasets, not current results; and the suggestion that the higher-order ratio *η* = *β*_2_*/β*_1_ separates microevolutionary from macroevolutionary timescales is a hypothesis from a small number of systems with no *β*_2_ null, not an established signature.

What remains open: rate equations for how the Betti numbers evolve under each evolutionary force; a Kolmogorov forward equation for the joint distribution of (*β*_0_, *β*_1_, *β*_2_) under composite processes; a full accounting of the quantitative relationship between recombination rates and topological persistence. These are the natural next steps for theoretical population genetics to take up.

### S5.7 Further reading

The main paper develops these ideas formally and carries the full bibliography. A reader wanting to go deeper has a few natural starting points. For the foundational result that recombination creates loops in the Vietoris-Rips filtration, the original paper is Chan, Carlsson, and Rabadán [8]; the quantitative refinement is in Lesnick and Rabadán [35]. For the Mapper algorithm as a framework for biological-data topology, Singh et al. [49] is the original source and Lum et al. [38] is the paper that established its vocabulary across biology. For our own group’s prior work on genomic-epidemiology Mapper and the Point Intersection Network, see Knudson et al. [34]. For the Colombian and Cambodian malaria contexts, Corredor et al. [11], Carrasquilla et al.[6], and Cárdenas et al. [3] are the key references. For the Arabidopsis 1001 Genomes project, the consortium paper [59] is canonical.

The computational tools used are freely available: Ripser for persistent-homology computation [45], the kmapper library for the Mapper algorithm, and the MalariaGEN Pf7 and Arabidopsis 1001 Genomes data are both open-access. All analysis code for this paper is distributed alongside it.

*This companion was prepared as a supplementary narrative to accompany the main manuscript. It is intended to be read by scientists outside topological data analysis who want the implications of the findings rather than the methods, and by students or generalists who want a way in to the full paper*.

## References

[1] Anderson TJC, Haubold B, Williams JT, et al. Microsatellite markers reveal a spectrum of population structures in the malaria parasite Plasmodium falciparum. Mol Biol Evol. 2000;17:1467–1482.

[2] Beaumont MA, Zhang W, Balding DJ. Approximate Bayesian computation in population genetics. Genetics. 2002;162:2025–2035.

[3] Cárdenas P, Corredor V, Santos-Vega M. Genomic epidemiological models describe pathogen evolution across fitness valleys. Science Advances. 2022;8:eabo0173 .

[4] Cárdenas-Sánchez A, Quiroga MA, Restrepo A. The mathematics of online polarization. Mention-graph topology of contested elections. Scientific Reports. 2022;12:1–14.

[5] Carlsson G. Topology and data. Bull Amer Math Soc. 2009;46 :255–308.

[6] Carrasquilla M, Early AM, Taylor AR, Knudson Ospina A, Echeverry DF, Anderson TJC, et al. Resolving drug selection and migration in an inbred South American Plasmodium falciparum population with identity-by-descent analysis. PLoS Pathog. 2022;18:e1010993 .

[7] Carriére M, Michel B, Oudot S. Statistical analysis and parameter selection for Mapper. J Mach Learn Res. 2018;19 :1–39 .

[8] Chan JM, Carlsson G, Rabadán R. Topology of viral evolution. Proc Natl Acad Sci. 2013;110:18566–18571.

[9] Cohen-Steiner D, Edelsbrunner H, Harer J. Stability of persistence diagrams. Discrete Comput Geom. 2007; 37:103 –120.

[10] Conway DJ, et al. High recombination rate in natural populations of Plasmodium falciparum. Proc Natl Acad Sci. 1999;96:4506 4511.

[11] Corredor V, Murillo C, Echeverry DF, Benavides J, Pearce RJ, Roper C, Guerra AP, Osorio L. Origin and dissemination across the Colombian Andes of sulfadoxine-pyrimethamine resistance in P. falciparum. Antimicrob Agents Chemother. 2010;54: 3121–3125.

[12] De Silva V, Carlsson G. Topological estimation using witness complexes. In: Eurographics SPBG. 2004.

[13] Doolittle WF . Phylogenetic classification and the universal tree. Science. 1999;284:2124–2129 .

[14] Drummond AJ, Rambaut A. BEAST: Bayesian evolutionary analysis by sampling trees. BMC Evol Biol. 2007; 7 :214.

[15] Edelsbrunner H, Harer JL. Computational topology: an introduction. AMS ; 2010.

[16] Edelsbrunner H, Letscher D, Zomorodian A. Topological persistence and simplification. Discrete Comput Geom. 2002;28:511–533 .

[17] Emmett KJ, Rabadán R. Characterizing scales of genetic recombination and antibiotic resistance in pathogenic bacteria using TDA. In: Brain Informatics and Health. Springer; 2014. p. 540–551.

[18] Feged-Rivadeneira A, Ángel A, González-Casabianca F, Rivera C. Malaria intensity in Colombia by regions and populations. PLoS ONE. 2018;13 :e0203673.

[19] Feged-Rivadeneira A, et al. Genomic epidemiology of SARS -CoV-2 in the Colombian Amazon basin. PLoS Negl Trop Dis. 2021;15:e0009327 .

[20] Felsenstein J. Inferring phylogenies. Sinauer; 2004.

[21] Fisher RA. The genetical theory of natural selection. Clarendon Press; 1930.

[22] Ghrist R. Barcodes: the persistent topology of data. Bull Amer Math Soc. 2008;45: 61–75.

[23] Haldane JBS . The causes of evolution. Longmans, Green; 1932.

[24] Battiston F, Cencetti G, Iacopini I, Latora V, Lucas M, Patania A, Young JG, Petri G. Networks beyond pairwise interactions: Structure and dynamics. Phys Rep. 2020;874:1–92.

[25] Bianconi G. Higher-Order Networks: An Introduction to Simplicial Complexes. Cambridge University Press; 2021.

[26] Feged-Rivadeneira A, González-Casabianca F, Ángel A, Corredor V. The shapes of evolution: a topological alphabet for macroevolutionary and coevolutionary processes. Companion manuscript, in preparation. 2026.

[27] Hatcher A. Algebraic topology. Cambridge University Press; 2002.

[28] Humphreys DP, McGuirl MR, Miyagi M, Blumberg AJ. Fast estimation of recombination rates using TDA. Genetics. 2019;211:1191–1204.

[29] Huson DH, Bryant D. Application of phylogenetic networks in evolutionary studies. Mol Biol Evol. 2006;23 (2):254–267 .

[30] Huson DH, Rupp R, Scornavacca C. Phylogenetic networks. Cambridge; 2010.

[31] Kauffman SA. The origins of order. Oxford University Press; 1993.

[32] Kelleher J, Etheridge AM, McVean G. Efficient coalescent simulation. PLoS Comput Biol. 2016;12:e1004842.

[33] Kimura M. The neutral theory of molecular evolution. Cambridge; 1983.

[34] Knudson A, González-Casabianca F, Feged-Rivadeneira A, et al. Spatio-temporal dynamics of P. falciparum transmission in the Colombian Pacific. Sci Rep. 2020;10: 3756.

[35] Lesnick M, Rabadán R, Rosenbloom DIS Quantifying genetic innovation. SIAM J Appl Algebra Geom. 2020;4:141–184.

[36] Lewontin RC. The apportionment of human diversity. Evol Biol. 1972; 6:381–398.

[37] Lewontin RC. The genetic basis of evolutionary change. Columbia University Press; 1974.

[38] Lum PY, et al. Extracting insights from the shape of complex data using topology. Sci Rep. 2013; 3:1236 .

[39] Maynard Smith J. The evolution of sex. Cambridge University Press; 1978.

[40] Nicolau M, Levine AJ, Carlsson G. Topology based data analysis identifies a subgroup of breast cancers with excellent survival. Proc Natl Acad Sci. 2011;108: 7265–7270.

[41] Ochman H, Lawrence JG, Groisman EA. Lateral gene transfer and the nature of bacterial innovation. Nature. 2000;405:299 304.

[42] Otto SP, Lenormand T. Resolving the paradox of sex and recombination. Nat Rev Genet. 2002; 3:252–261.

[43] Petri G, Expert P, Turkheimer F, Carhart-Harris R, Nutt D, Hellyer PJ, Vaccarino F. Homological scaffolds of brain functional networks. J R Soc Interface. 2014;11(101):20140873 .

[44] Rabadán R, Blumberg AJ. Topological data analysis for genomics and evolution. Cambridge University Press; 2019.

[45] Bauer U. Ripser: efficient Vietoris-Rips persistence barcodes. J Appl Comput Topol. 2021;5: 391–423 .

[46] Robertson DL, et al. HIV-1 nomenclature proposal. Science. 2000;288:55–56.

[47] Saggar M, et al. Dynamical organization of the brain using TDA. Nat Commun. 2018; 9:1399.

[48] Sheehy DR. Linear-size approximations to the Vietoris-Rips filtration. Discrete Comput Geom. 2013;49 : 778–796 .

[49] Singh G, Mémoli F, Carlsson GE. Topological methods for high dimensional data sets. In: SPBG. 2007. p. 91–100.

[50] Steel J, Lowen AC. Influenza A virus reassortment. Curr Top Microbiol Immunol. 2014; 385:377 401.

[51] Taylor AR, Echeverry DF, Anderson TJC, Neafsey DE, Buckee CO. Identity-by-descent with un-certainty characterises connectivity of P. falciparum on the Colombian-Pacific coast. PLoS Genet. 2020;16 :e1009101.

[52] Volkman SK, et al. A genome-wide map of diversity in Plasmodium falciparum. Nat Genet. 2007 ; 39:113 – 119.

[53] Wright S. The roles of mutation, inbreeding, crossbreeding, and selection in evolution. Proc Sixth Int Congress Genet. 1932;1:356 –366 .

[54] Zomorodian A, Carlsson G. Computing persistent homology. Discrete Comput Geom. 2005; 33:249 274.

[55] MalariaGEN. Pf7 : an open dataset of Plasmodium falciparum genome variation in 20,000 worldwide samples. Wellcome Open Res. 2023;8:22.

[56] Gusfield D. ReCombinatorics: the algorithmics of ancestral recombination graphs and explicit phylogenetic networks. MIT Press; 2014.

[57] 1001 Genomes Consortium. 1,135 genomes reveal the global pattern of polymorphism in Arabidopsis thaliana. Cell. 2016;166 :481–491.

[58] François O, Blum MGB, Jakobsson M, Rosenberg NA. Demographic history of European populations of Arabidopsis thaliana. PLoS Genet. 2008;4:e1000075.

[59] Alonso-Blanco C, Andrade J, Becker C, et al. 1,135 genomes reveal the global pattern of polymorphism in Arabidopsis thaliana. Cell. 2016;166 :481–491.

[60] Hudson RR. Properties of a neutral allele model with intragenic recombination. Theor Popul Biol. 1983 ;23 :183 –201.

[61] Griffiths RC, Marjoram P. An ancestral recombination graph. In: Donnelly P, Tavaré S, editors. Progress in Population Genetics and Human Evolution. Springer; 1997. p. 257–270.

[62] Kahle M. Random geometric complexes. Discrete Comput Geom. 2011;45:553–573 .

[63] Kahle M. Topology of random simplicial complexes: a survey. In: Algebraic Topology: Applications and New Directions, Contemporary Mathematics 620. AMS ; 2014. p. 201–221.

[64] Billera LJ, Holmes SP, Vogtmann K. Geometry of the space of phylogenetic trees. Adv Appl Math. 2001;27 :733–767 .

[65] Head T, MechCoder Louppe G, et al. scikit-optimize/scikit-optimize. Zenodo; 2020. doi:10.5281/zenodo.4014775.

